# A computational proposal for tracking multiple molecules in a multi-focus confocal setup

**DOI:** 10.1101/2022.05.17.492362

**Authors:** Sina Jazani, Lance W.Q. Xu, Ioannis Sgouralis, Douglas P. Shepherd, Steve Pressé

## Abstract

Tracking single molecules continues to provide new insights into the fundamental rules governing biological function. Despite continued technical advances in fluorescent and non-fluorescent labeling as well as data analysis, direct observations of trajectories and interactions of multiple molecules in dense environments remain aspirational goals. While confocal methods provide a means to deduce dynamical parameters with high temporal resolution, such as diffusion coefficients, they do so at the expense of spatial resolution. Indeed, on account of a confocal volume’s symmetry, typically only distances from the center of the confocal spot can be deduced. Motivated by the need for true three dimensional high speed tracking in densely labeled environments, we propose a computational tool for tracking many fluorescent molecules traversing multiple, closely spaced, confocal measurement volumes providing independent observations. Various realizations of this multiple confocal volumes strategy have previously been used for long term, large area, tracking of one fluorescent molecule in three dimensions. What is more, we achieve tracking by directly using single photon arrival times to inform our likelihood and exploit Hamiltonian Monte Carlo to efficiently sample trajectories from our posterior within a Bayesian nonparametric paradigm. A nonparametric paradigm here is warranted as the number of molecules present are, themselves, *a priori* unknown. Taken together, we provide a computational framework to infer trajectories of multiple molecules at once, below the diffraction limit (the width of a confocal spot), in three dimensions at sub-millisecond or faster time scales.

## Introduction

Fluorescence confocal microscopy^1–3^ (FCM) is a non-invasive technique widely used to observe dynamics of molecules both *in vitro* and *in vivo*.^4–10^ One of the advantages of FCM is its ability to detect photons at very rapid time scales, as high as ~10 MHz, limited only by photon emission and detection in a conventional single-focus confocal microscope. Our ability to detect photons at those time scales makes it, in principle, possible to extract dynamical information on fast processes from single photon arrival times approaching data acquisition time scales.^11–13^ A severe drawback in determining trajectories in three dimensions, is that, for a conventional FCM setup consisting of a single confocal volume that is held stationary, we can only acquire information on the distance of molecules from the center of a single stationary confocal volume.^11–13^ This raises the question as to whether we could learn the position of molecules, potentially many molecules simultaneously, with the high temporal resolution afforded by FCM, *i.e.*, in non-scanning mode, at a localization precision exceeding that of the size of a single confocal volume.^2,10,14^

It has been shown in the literature that to learn molecular positions over time, we can either track the targeted molecule by moving multi-focus confocal volumes^15–17^ or using methods such as MINFLUX.^18,19^ The challenge here is that these approaches can currently track *one* molecule at a time. Consequently, the existence of *more* than one molecules within the confocal volume regions which we consider as the region of interest (ROI) leads to biased or altogether incorrect trajectory estimates. These approaches are therefore limited to low density experiments which ensure typical inter-molecular distances far exceeding the dimensions of a single confocal volume. As such, these experiments are carried out using low-density labeling in a dense molecular environment and subsequently infer behavior of the whole population from this labeled sub-population.

To overcome this limitation, critical to many applications requiring high concentrations, here we propose to analyze data from a set of stationary multi-focus confocal volumes (identical to the experimental data we analyzed for a single spot in the past^11–13^) and attempt tracking in more densely labeled environments.

As we will show, our computational approach will also estimate trajectories at the fastest possible time scale available from the raw data, i.e., from single photon arrival times. Tracking from single photon arrival times is relevant to confocal methods currently relying on photon binning at low labeling density.^15-17,20,21^ By virtue of using single photon arrival times to achieve tracking, we will be able to get away with limited light exposure to our sample to determine trajectories and, as such, can limit photo-damage to a light-sensitive sample as well as monitor processes resolved over short periods of time.^11,13^

To achieve multiple molecule tracking, we must address the fundamental “model selection problem” since an unknown number of molecules within one confocal spot may be present at once. That is, depending on how many molecules we estimate to be in the volume, we obtain different trajectory and diffusion coefficient estimates.^11–13^ It is because the number of molecules contributing photons to the detector is unknown that we propose a nonparametric Bayesian framework.^10,22,23^ In doing so, we extend our previous work on single-spot detector with single photon arrival times as its output.^13^ In doing so, we achieve multi-particle tracking with spatial resolution below the width of a single confocal spot.

We illustrate the experimental multi-focus confocal setup in Fig. 1(a) with single-spot illumination and multi-spot detectors. The product of the excitation and detection point spread functions (PSFs) provide four neighboring confocal volumes, Fig. 1(b), and examples of photon arrival times, in each detection channel, are illustrated in Fig. 1(c).

**Figure 1:**
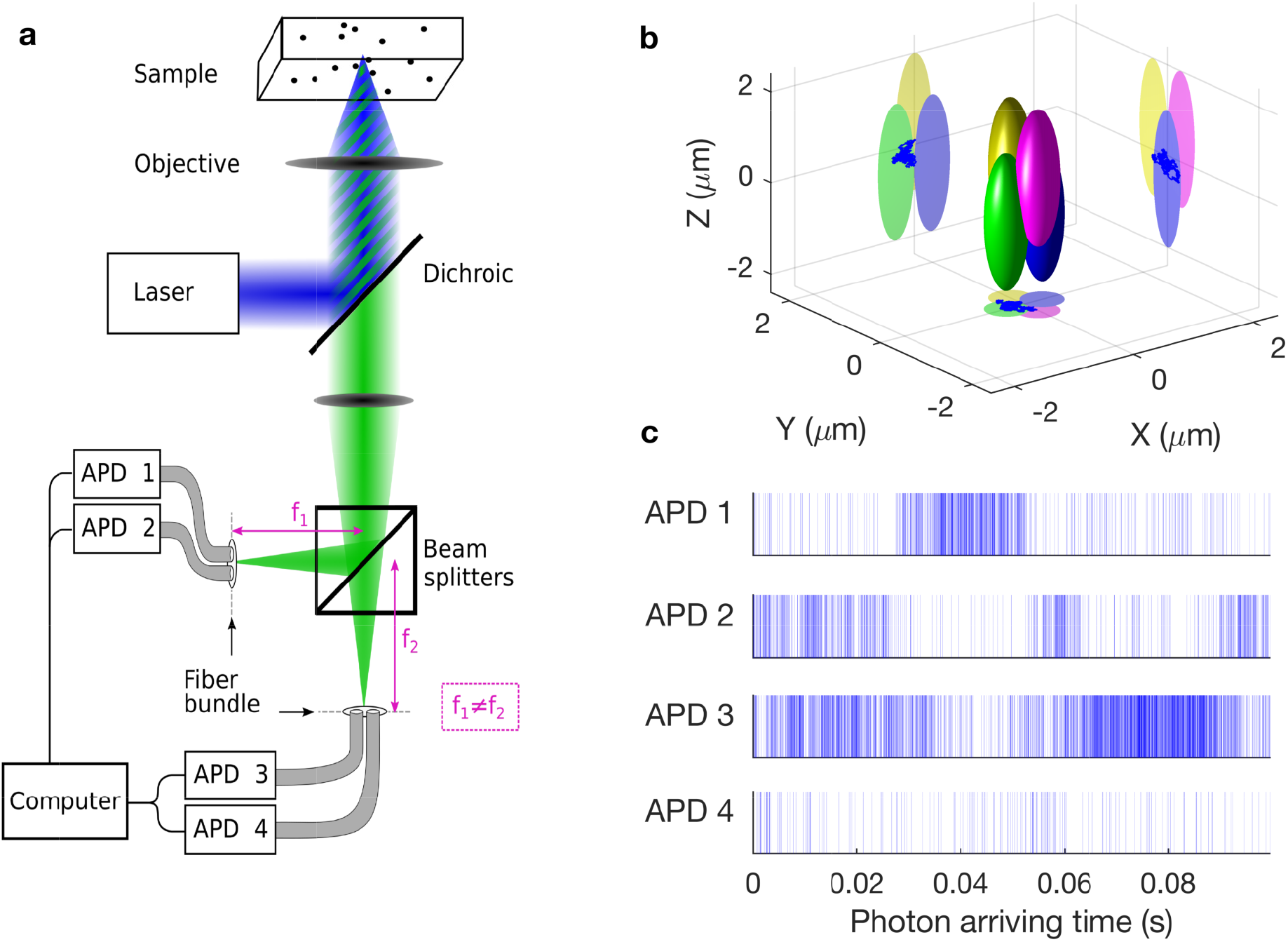
Illustration of a simultaneous multi-focus confocal setup. **(a)** Schematic of a single-spot illumination and multi-spot detector forming a microscope with multi-focus confocal volumes. **(b)** Idealized PSF shapes in the object space (originating from the overlap of the excitation and detection PSFs). The confocal volumes are shown as ellipsoids for visual purposes. **(c)** Single photon arrival time traces, illustrating photon detection times over the course of an experiment by the four axially and laterally shifted detection volumes. Each panel corresponds to a different channel.

By proposing to use single photon arrival times at each detector, just as we had for single detectors in Refs.,^11–13^ we will break symmetry and uniquely learn three dimensional trajectories of many molecules below the conventionally-defined diffraction limit at the fastest time scale available, *i.e*., limited only by single photon arrival times.

While we consider typical asymmetric Gaussian confocal volume shapes, as the framework and computational scheme is general, we can treat any confocal volume shape.

## Results and discussion

### Overview

The method we propose can estimate trajectories of multiple molecules simultaneously from a multi-focus confocal microscope using the mathematics of Bayesian nonparametrics.^10,22,23^ Our method starts from single photon arrival times. The goal is to estimate, i.e., obtain a posterior distribution, over all unknowns. These include: the number of molecules contributing photons whose arrival times constitute our observation, the location of the molecules at each point in time (trajectories), the diffusion coefficient of the molecules, molecular brightnesses and background photon emission rates for each detector.

### Validation under broad parameter ranges

To validate our method’s robustness, we generate synthetic single photon arrival times as our observation, assuming a multi-focus confocal microscopy setup, under a broad range of: (i) diffusion coefficients, Fig. 2; (ii) numbers of labeled molecules, Fig. 3; (iii) molecular brightnesses, Fig. 4; and (iv) positions of detection spots, Fig. 5.

**Figure 2:**
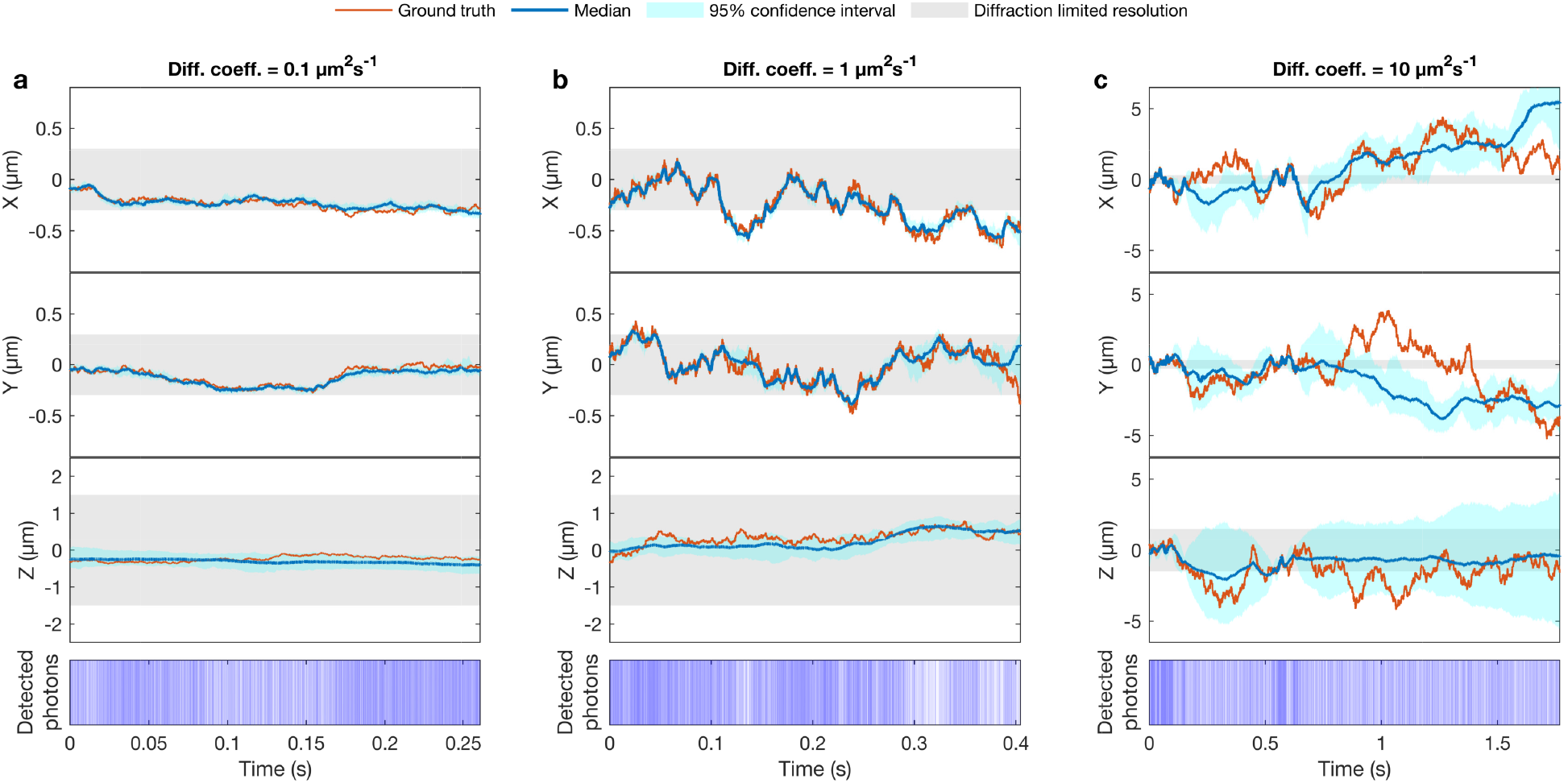
Effect of the diffusion coefficient on the precision of the trajectory estimate. **(a-c)** Trajectory estimate of a single freely diffusive molecule with diffusion coefficient of 0.1 *μ*m^2^s^-1^, 1 *μ*m^2^s^-1^ and 10 *μ*m^2^s^-1^ in 3D, respectively. These values cover the range of protein and nucleic acid diffusion constants *in vivo*.^24,25^ As we note in the text, we remark on the growing error bars as the diffusion coefficient increases. Note that for clarity y-axis on a-c have different bounds. In all three cases, we used molecular brightnesses of 5 × 10^4^ photons s^-1^ and background photon emission rates of 10^3^ photons s^-1^. These values are identical to those drawn from real data in Refs.,^11,13^ have been used to generate the trajectories. The photon detection times from all detectors, are shown with blue lines and the total number of detected photons for all cases is 10^4^. The gray areas represent the width of a single 3D gaussian confocal volume which for X, Y and Z coordinates are equal to 2 × *ω_x_*, 2 × *ω_y_* and 2 × *ω_z_*, respectively.

**Figure 3:**
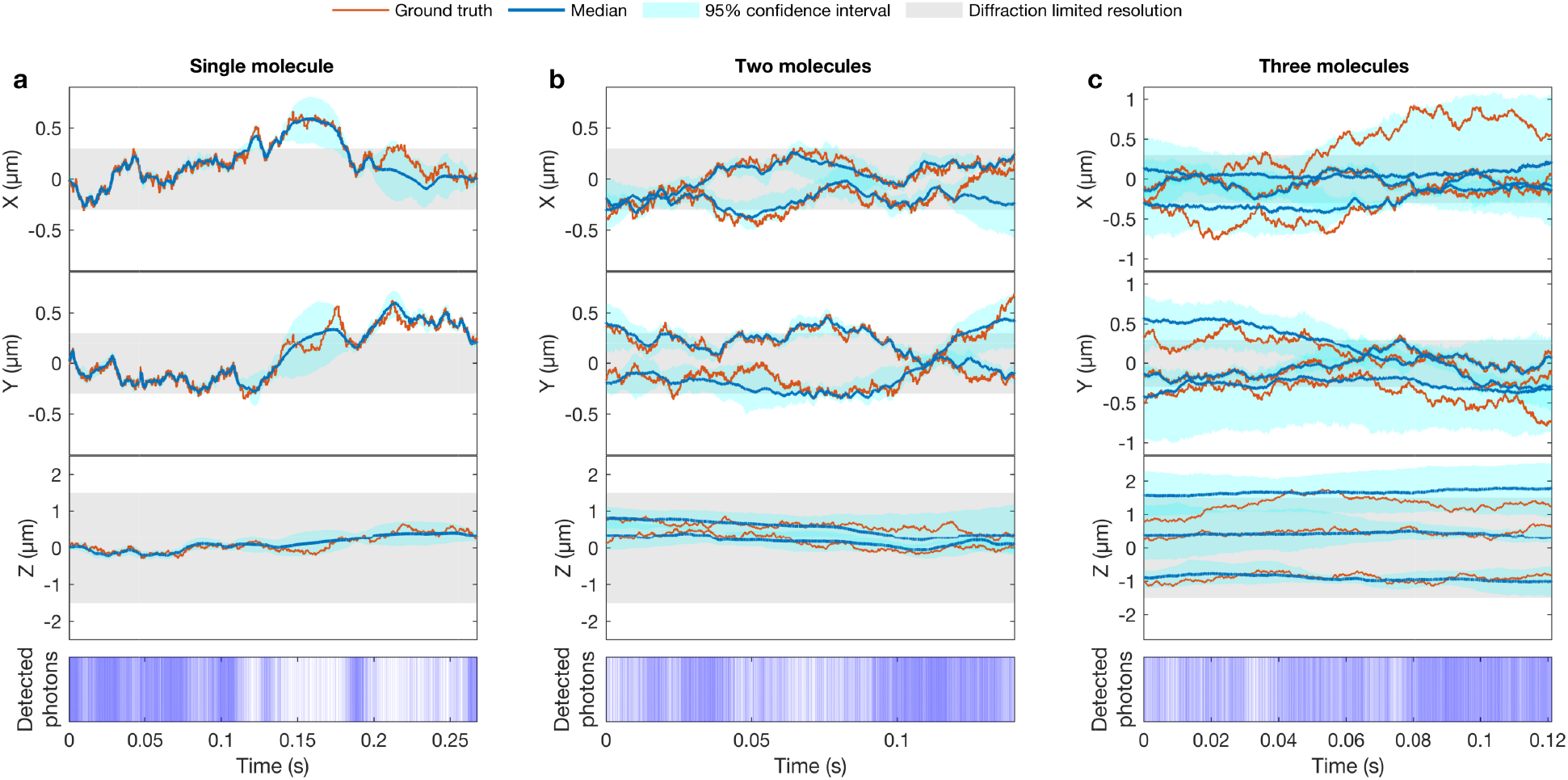
Effect of the number of molecules contributing photons on the precision of the trajectory estimate. **(a-c)** Trajectory estimates for one, two and three freely diffusing molecules, respectively. All three cases were generated with a diffusion coefficient of 1 *μ*m^2^s^-1^, molecular brightness of 5 × 10^4^ photons s^-1^ and background photon emission rate of 10^3^ photons s^-1^. The photon detection times are shown with blue lines and the total number of detected photons for all cases is 10^4^. For approximate confocal volumes of ~ 2.11 fL, dividing the number of molecules by the volume, these coincide with approximate concentrations of 0.79, 1.57 and 2.36 nM.

**Figure 4:**
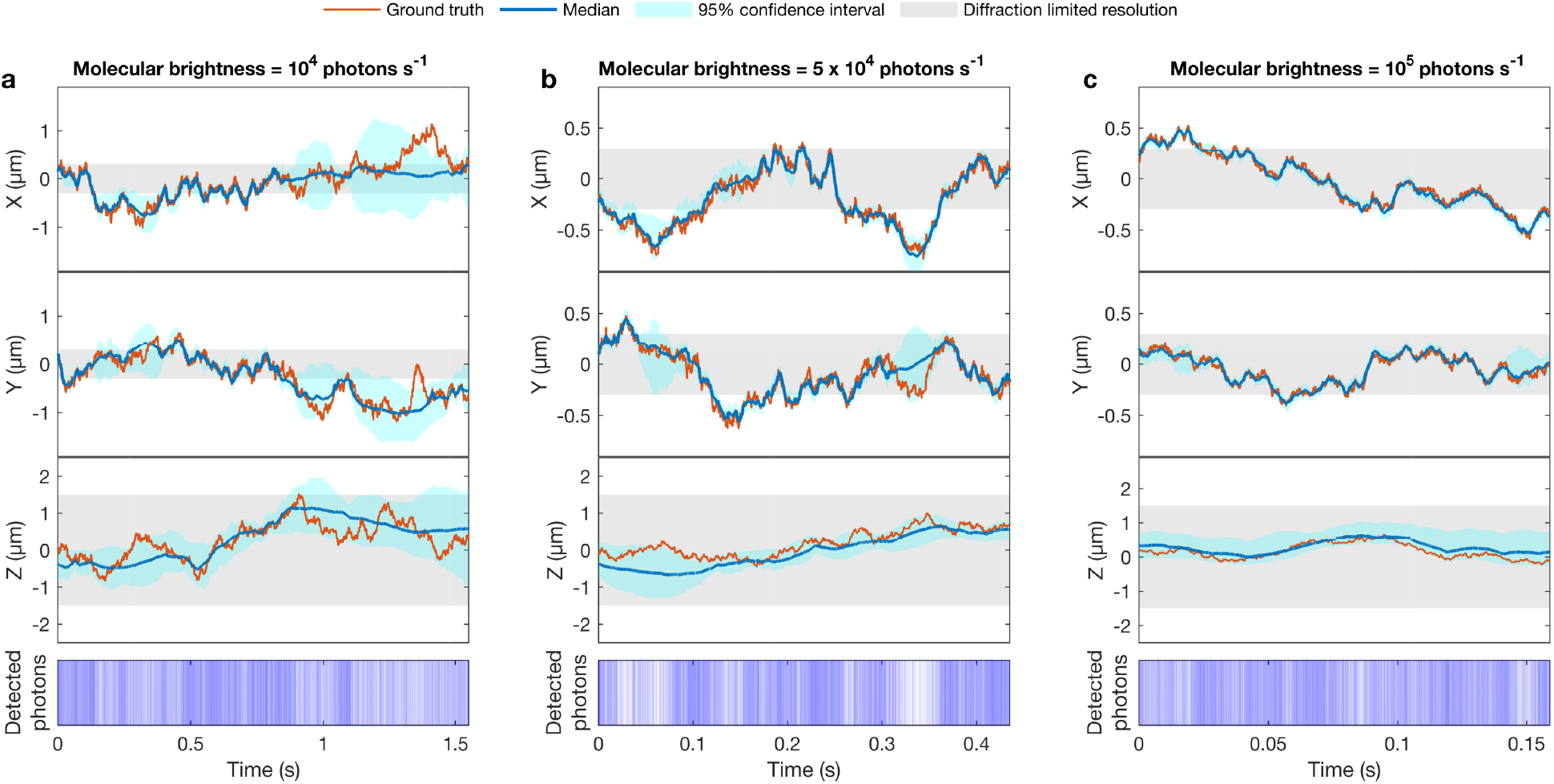
Effect of the molecular brightness on the precision of the trajectory estimate. **(a-c)** Trajectory estimates for a single freely diffusive molecule with molecular brightness of 10^4^ photons s^-1^, 5 × 10^4^ photons s^-1^ and 10^5^ photons s^-1^ for all detection spots, respectively. These values are motivated by Refs.^11,13,26^ For all three cases, diffusion coefficient of 1 *μ*m^2^s^-1^ and the background photon emission rate of 10^3^ photons s^-1^ have been used to generate the trajectories. The photon detection times are shown with blue lines and the total number of detected photons for all cases is 10^4^.

**Figure 5:**
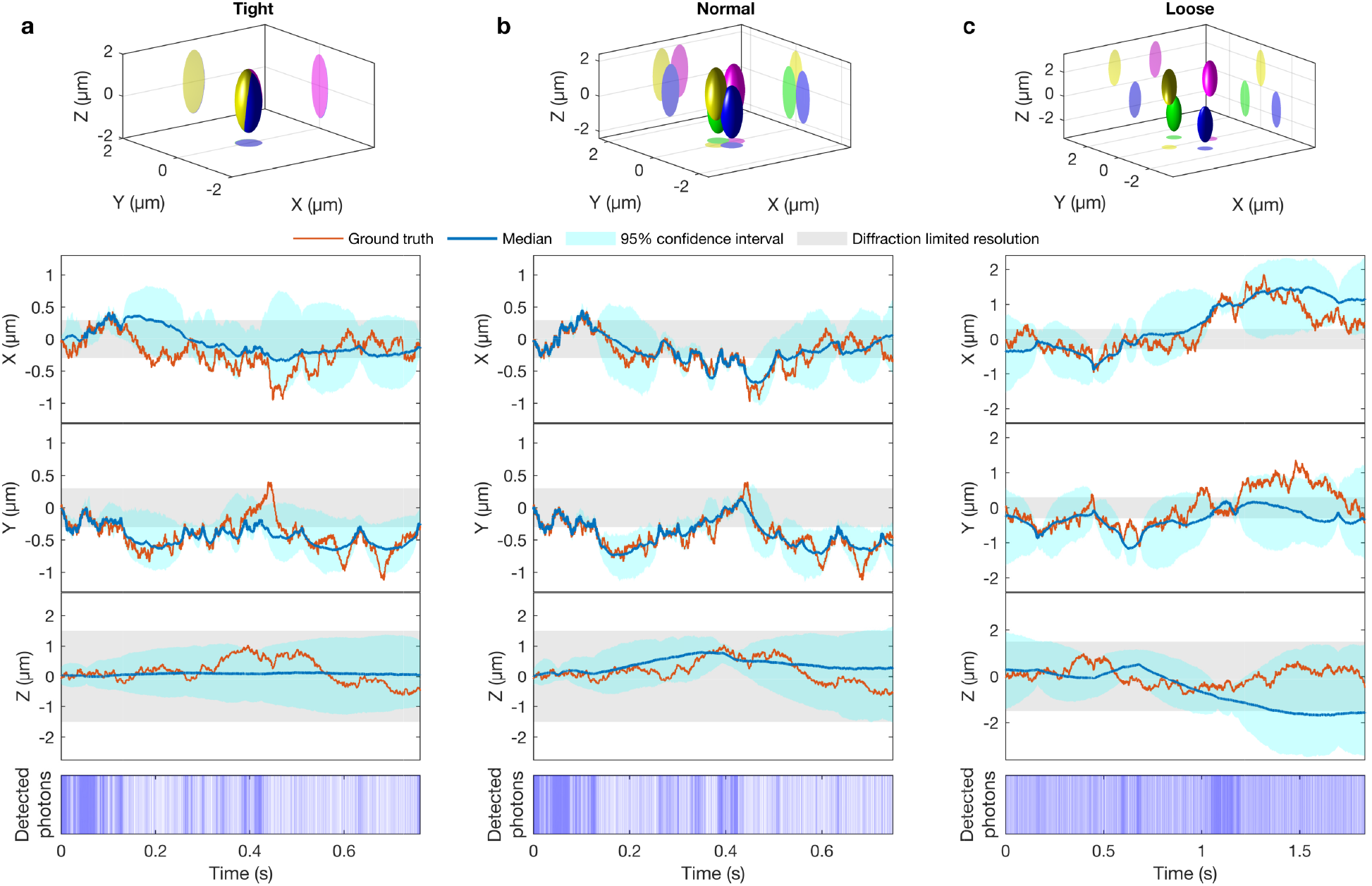
Effect of the configuration of the confocal volumes on the precision of the trajectory estimate. **(a)** Trajectory estimates of a freely diffusive molecule where the confocal volumes overlap substantially and are located at (−0.01, 0.00, −0.01), (0.01, 0.00, −0.01), (0.00, −0.01, 0.01) and (0.00, 0.01, 0.01). **(b)** Trajectory estimates of a freely diffusive molecule with confocal volumes minimally overlapping and located at (−0.3, 0.0, −0.5), (0.3, 0.0, −0.5), (0.0, −0.3, 0.5) and (0.0, 0.3, 0.5). **(c)** Trajectory estimates of a freely diffusive molecule with confocal volumes are far away and located at (−1.0, 0.0, −1.5), (1.0, 0.0, −1.5), (0.0, −1.0, 1.5) and (0.0, 1.0, 1.5). For all three cases, coordinates are with respect to the point of origin (in units of *μ*m) and the diffusion coefficient of 1 *μ*m^2^s^-1^, molecular brightness of 5 × 10^4^ photons s^-1^ and background photon emission rate of 10^3^ photons s^-1^ have been used to generate the trajectories. The photon detection times are shown with blue lines and the total number of detected photons for all cases is 10^4^.

The marginal posteriors over the molecular trajectories that we obtain across all figures are informed from the analysis of one observation trace which is the result of the combination of observation traces collected by each detector channel. In those, the breadth of the posterior (i.e., the standard deviation), which is a measure of the accuracy of our estimate, reflects the uncertainty introduced by the finiteness of the data and the inherent noise in the observation traces. To be able to evaluate the precision of our method, we compared the targeted posteriors of the molecular trajectories with ground truth trajectories. For quick reference, all results are summarized in SI Table. S1.

To begin, in Fig. 2, we consider four 3D Gaussian confocal volumes of size *ω_xy_* = 0.3 *μ*m and *ω_z_* = 1.5 *μ*m at positions (0.3, 0.0, 0.5), (−0.3, 0.0, 0.5), (0.0, 0.3, −0.5) and (0.0, −0.3, −0.5) with respect to the point of origin (in units of *μ*m). These confocal volume sizes are motivated from Refs.^11–13^ and, as we can see from Fig. 5(b), these confocal volumes exhibit limited overlap. We generate synthetic data, just as we had generated and benchmarked with real data in Refs.^11,13^ for a single-focus confocal volume, with different concentration of molecules within ROI. We consider diffusing molecules with diffusion coefficients of 0.1 *μ*m^2^s^-1^, 1 *μ*m^2^s^-1^ and 10 *μ*m^2^s^-1^. These values cover a range of measured protein diffusion coefficients *in vivo*.^24,25^ In all cases, for consistency, we based our analysis on just 10^4^ photons emitted from all molecules and background and detected by all confocal volumes. Thus, the time duration of each trajectory varies based on the molecular brightness of the molecules which, in turn, depends on the molecule’s location within the excitation and detection profile.

Beyond learning the number of molecules and their associated trajectories, from this synthetic data, we also learn diffusion coefficients, molecular brightnesses and background photon emission rates simultaneously; see SI Fig. S1. First off, without prior knowledge of the number of molecules, we recover a posterior over numbers of molecules sharply peaked at 1; see SI Fig. S1. That is, we do not put in this information by hand to obtain the results shown either in this figure or in any other. This information is critical as, without an appropriate estimate of the number of molecules giving rise to the photons, our diffusion coefficient estimates would be inaccurate; see Fig. 2 of our previous work. 13 And, it is fundamentally for this reason, that we must abandon the parametric Bayesian paradigm and use a nonparametric framework. Also, in Fig. 2 we can see that greater diffusion coefficients naturally lead to fewer detected photons (and thus less data) as molecules traverse the ROI. As a result, the posterior over the molecule’s trajectory is broader. Put differently, less information or fewer photons implies less trajectory certainty and thus broader posteriors. Conversely, the slower a molecule diffuses, the more photons are collected, the sharper the posterior estimate of the molecular trajectories, Fig. 2(a).

In Fig. 3 we demonstrate the robustness of our method as we generate synthetic data with an increasing number of molecules in the ROI. Beyond learning the number of molecules and their associated trajectories, from this synthetic data, we also learn diffusion coefficients, molecular brightnesses and background photon emission rates simultaneously; see SI Fig. S3. It is worth mentioning that the posterior over the molecular trajectories is now broader when more molecules are present in the confocal volume.

That is, for two molecules as compared to one, we have greater uncertainty in both trajectories. This is because the same number of photons is now needed to learn twice as many trajectories as we had earlier. Thus the information we have available per trajectory drops. Fig. 3 also allows us to begin addressing how many photons we need to set uncertainty bounds on molecular trajectories given experimental parameters such as confocal volume dimensions, molecular diffusion coefficients and molecular brightness.

In Fig. 4, we demonstrate the robustness of our method with respect to the molecular brightness. That depends on the photon emission rate of the molecules and the quantum yield of detectors. So, for each detector, we have different value for the molecular brightness and background photon emission rate. We provide more detail in SI Sec. S3.2. In particular, due to different quantum yields, we end up with different molecular brightness and background photon emission rate for each detector. Here, we evaluate the method with molecular brightnesses of 10^4^, 5×10^4^ and 10^5^ photons s^-1^ and a background photon emission rate of 10^3^ photons s^-1^; these are values in the range of those previously reported in Refs.^24,25^ As expected, higher molecular brightness leads to sharper posteriors. We also learn diffusion coefficients, molecular brightnesses and background photon emission rates simultaneously; see SI Fig. S5.

Finally, in Fig. 5 we test the robustness of our method as we vary the configuration of the four neighboring confocal volumes. Here, we consider the case that the four confocal volumes are located symmetrically located at (−0.01, 0.00, −0.01), (0.01, 0.00, −0.01), (0.00, −0.01, 0.01) and (0.00, 0.01, 0.01) in Fig. 5(a), (−0.3, 0.0, −0.5), (0.3, 0.0, −0.5), (0.0, −0.3, 0.5) and (0.0, 0.3, 0.5) in Fig. 5(b) and (−1.0, 0.0, −1.5), (1.0, 0.0, −1.5), (0.0, −1.0, 1.5) and (0.0, 1.0, 1.5) in Fig. 5(c) with respect to the point of origin (in units of *μ*m). Except the case that the confocal volumes overlap substantially, these distances are within the realm of what can be achieved in current experimental setups.^15,16,27-30^

There exists an optimal, “goldilocks”, distance between confocal volumes that allows us to minimize trajectory uncertainty. As we see in Fig. 5(a), if the confocal volumes are too close, the information they provide on the molecular positions is redundant. Put differently, in the extreme case that all volumes exactly overlap, then we revert to the highly symmetric case of the single-spot (at which point we can only determine the effective distance of each molecule from the center of the single-focus confocal volume but not its 3D position).^11–13^ Similarly, in the case that the confocal volumes are too far from each other, as in Fig. 5(c), each confocal volume provides independent information from other spots and the position of each molecule cannot be interpolated between confocal volumes. Thus, in this extreme limit as well, we revert to the single-focus confocal volume regime.

Although, it is clear that when confocal volumes overlap, our trajectory estimates improve, it is hard to say precisely what distance is optimal. This chiefly depends on the precise shape of the confocal volumes (we assumed a Gaussian, but this can be replaced by any analytical form), the molecular brightnesses and background photon emission rates. Beyond learning the number of molecules and their associated trajectories, from this synthetic data, we also learn diffusion coefficients, molecular brightnesses and background photon emission rates simultaneously; see SI Fig. S7.

## Conclusion

We have proposed a new analysis strategy using existing stationary multi-focus confocal microscopes to track multiple molecules at once. The use of a multi-focus confocal microscope is required in order to break the symmetry of the single-focus confocal microscope^12^ which allows for multiple molecules tracking from single photon arrival times but only up to a symmetry originating from the symmetry of the confocal volume. Multiple confocal volumes not only break symmetry but also provide non-redundant information at each detector. This allows us to have localization precision as great as ten times that of single-focus confocal volume width (see Figs. S2, S4, S6 and S8); though, the magnitude of the improvement is sensitive to the diffusion coefficient, molecular brightnesses and background photon emission rates; see SI Figs. S1, S2, S3 and S4.

We track many molecules simultaneously within a multi-focus confocal setup by exploiting not just temporal, but also spatial information encoded within photon arrival time traces. The ability to exploit information as it arrives one photon at a time, without binning, is particularly critical in minimizing light exposure or, in the future, to probe rapid processes resolved over rapid time scales spanning the arrival of thousands of photons.

Minimizing light exposure is not the only downstream advantage we see to our method as we envision more complex *in vivo* applications. Beyond tracking, our method retains molecular identity; for example, in Fig. 3 we see that trajectories can be discriminated even if molecular trajectories move closer to each other than the conventionally defined diffraction limit though not indefinitely. Retaining molecular identity is an advantage in resolving single molecule events unique to individual molecules with high localization precision.

To achieve such tracking, we use Beta-Bernoulli processes^31,32^ and other tools from computational statistics that we recently benchmarked on a variety of single confocal volume experiments.^11,13^ As we discuss in the methods section, our method can be generalized to treat alternative forms for the confocal volumes which can be distorted in real experiments or even motion models for the molecules beyond Brownian motion. Assessing, computationally, the efficiency of our posterior sampling for shapes and dynamical motion models beyond that considered, and warranted by alternative multi-focus experiments, is worthy of future study.

The method we propose here differs from existing methods such as multi-focus confocal microscopy,^15–17^ scanning confocal volume^33^ and MINFLUX^18,19,34^ in a number of ways. First existing methods localize and track labeled molecules one at a time and under low density conditions to avoid inaccurate tracking induced by overlapping molecular trajectories. Second, if photobleaching is avoided, they can track molecules over large distances. By contrast, our method tracks over shorter distances (as our method tracks over a fixed volume a few molecules) but is, critically, not limited to low density. What is more, we achieve high temporal resolution without photon binning.

Finally, we add that we considered four stationary confocal volumes and varied their distances; see Fig. 5. However, as few as three confocal volumes, both axially and laterally displaced, would be sufficient to pinpoint positions in 3D albeit with greater uncertainty. While we track over limited distances, it is also conceivable, in the future, that the number of confocal volumes could possibly be increased to span broader areas.^28-30,35-37^

## Method

### Model overview

Here we consider an optical apparatus consisting of multi-focus confocal volumes with, for sake of concreteness, four detectors all simultaneously collecting photons; see Fig. 1. This allows us to break the symmetry imposed by a single-focus confocal volume^11–13^ and consequently track (fluorescing) molecules in 3D.

Based on the graphical model provided in Fig. 6, our goal is to learn the variables: 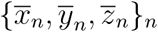 representing the trajectories of the molecules, *D* representing the diffusion coefficient, 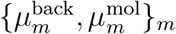 representing the molecular brightnesses and background photon emission rates and {*b_n_*}_*n*_ representing the indicators of molecules which tell us about the population of molecules contributing to the data. We tag the molecules with indices *n* = 1,…, *N* and the detectors with indices *m* = 1,…, *M*.

**Figure 6:**
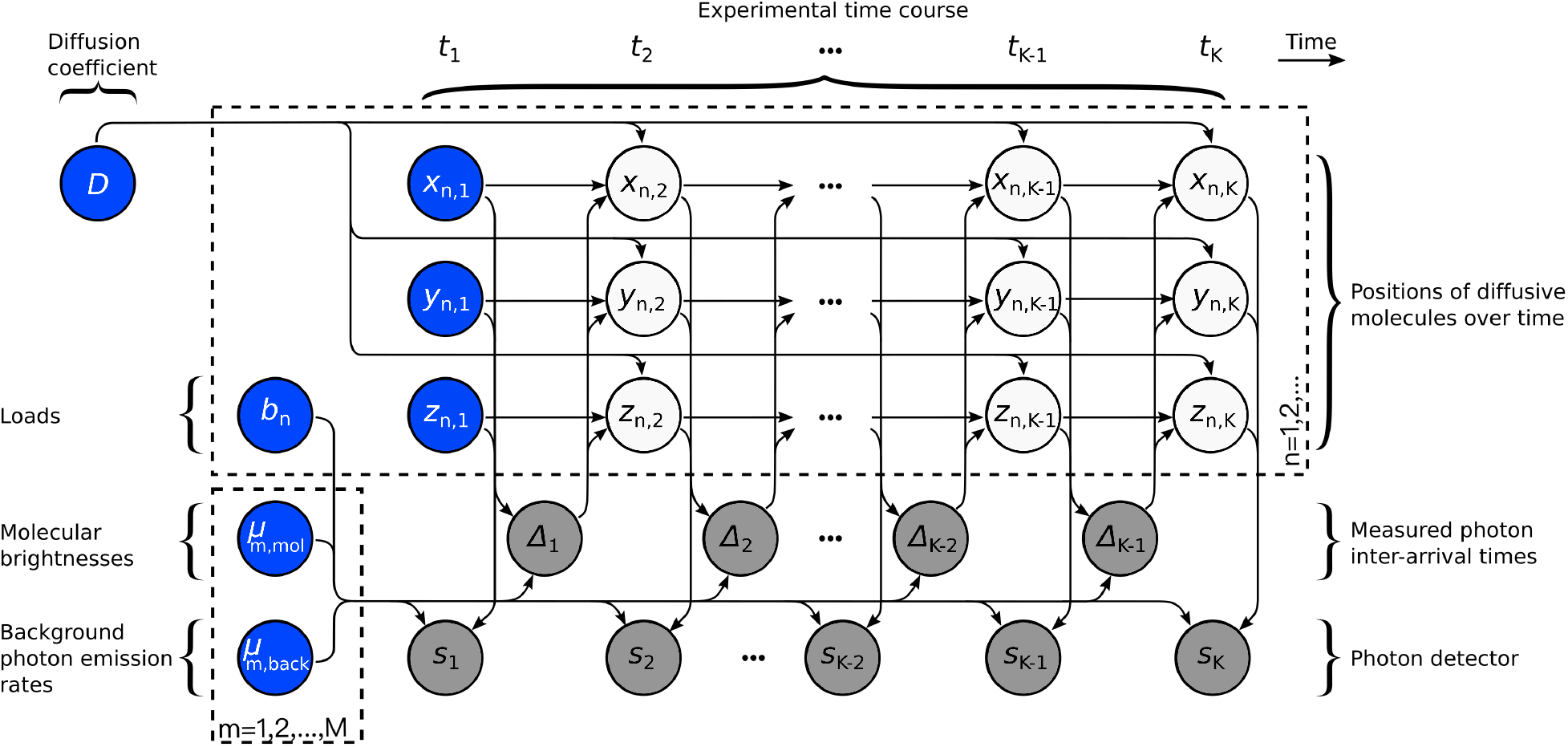
Graphical representation of the model for multiple molecules and many confocal volumes. Molecules diffuse over the course of the experiment. Here, (*x_n,k_, y_n,k_, z_n,k_*) denotes the location of molecule *n* at time *t_k_* and the photon arrivals over all confocal volumes are marked by *k* = 1, 2,…, *K*. During the experiment, *Δ_k_*, is the interarrival time which is equal to the time difference between successive photon detection times *t_k_* and *t*_*k*+1_. The *k*^th^ photon can be detected by any of the detectors *s_k_* = 1,…, *M*. In full generality, as each confocal volume may have different properties, we index the molecular brightness and the background photon emission rate of each as follows: 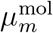 and 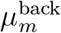 for *m* = 1,…, *M*. The diffusion coefficient *D* dictates the molecular dynamics and determines future molecular locations which, in turn, determine the location of the molecule within the illuminated volume, influences the photon emission rates and, ultimately, the photon arrival times. In order to learn the number of existing molecules we introduce auxiliary variables *b_n_* termed “loads”. Following machine learning convention,^42^ the circle surrounding measurement, 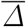 and 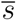, random variables are shaded in grey while model variables requiring prior probability distributions are shaded in blue.

As the full joint posterior over these variables does not assume an analytic form, we must develop a numerical scheme to sample from our posterior. For this reason, we generate samples of our posterior through a Gibbs sampling scheme.^38–40^ That is, we update each one of the variables sequentially by sampling from the marginal posterior conditioned on all other variables as well as the measurements which we describe below.

In greater detail, we start from single photon arrival times. These occur at times *t_k_* with *k* = 1,…, *K* and may be reported from any of *M* different detectors. For clarity, we combine the measurements from all detector channels into two observation traces: the interarrival times between successive photons, 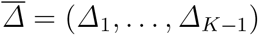; and the tags of the confocal volumes where each photon was collected, 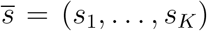. According to our convention, Δ_*k*_ = *t*_*k*+1_ – *t_k_* is the inter-arrival time between two detected photons and *s_k_* is the tag of the confocal volume which observed the photon at time *t_k_*.

Our goal is to use the data in 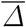 and 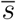 to construct a likelihood^40,41^ and, through it, a posterior probability distribution^40,41^ which yields estimates of (i) the number of molecules contributing photons to the data; (ii) the trajectories of these molecules; (iii) their diffusion coefficient and brightness; and, (iv) the background photon emission rate, which are represented by the variables introduced above.

In this section, we explain the way to achieve each one of these in detail. Further details and a computational implementation can be found in the SI Sec. S.5. A summary of the notation, abbreviations and mathematical definitions can be found in SI Tables S.3 and S.4. The graphical summary of the model explained below is shown on Fig. 6.

### Model description

Since we have two independent inputs, 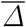 and 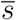, our likelihood is expressed by two independent parts: one expressing photon emission and one expressing photon detection. Respectively, these are:

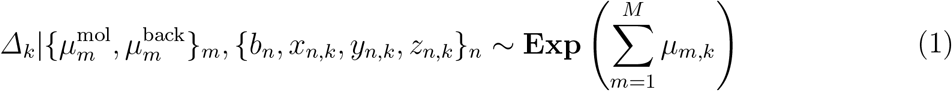

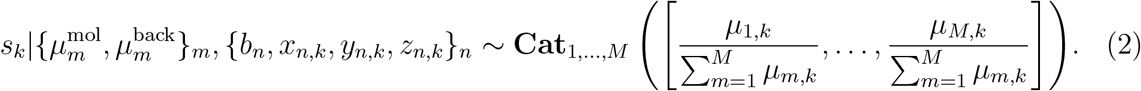

In the above, *μ_m,k_* is the rate of detected photons at the *m^th^* detector at the *k^th^* timepoint. Eq. (1) applies to *k* = 1,…, *K* – 1 and Eq. (2) applies to *k* = 1,…, *K*. The photon emission rates are given by

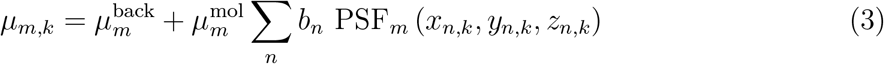

and the point spread function for each of the confocal volumes, PSF_m_ (*x,y, z*), indexed *m* is separately defined. We provide more detail in SI Sec. S3.2.

Here, for all the synthetic observation traces, we consider all confocal volumes as 3D Gaussians, but as we explain in the SI Sec. S.3.3, any functional form can be readily incorporated for the generation of analysis data

We mention that, since each confocal volume coincides with a different detector receiving photons from a different region of physical space, we represent different molecular brightnesses and background photon emission rates 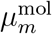 and 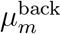 associated to each confocal volume *m*. We provide more detail in SI Sec. S3.2.

We assume a Brownian motion model^11–13^ (with unknown diffusion coefficient) connecting individual positions in the trajectories, 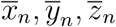, across time by

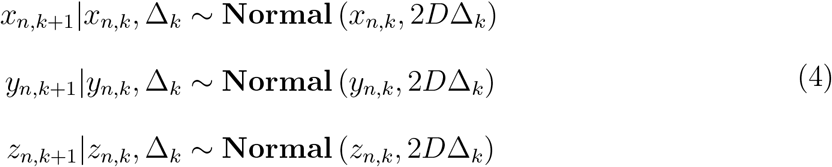

where, *k* = 1,…, *K* – 1.

### Model inference

The quantities that we wish to estimate are: the diffusion coefficient *D*, molecular brightnesses 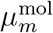 and background photon emission rates 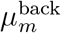 with respect to each of the confocal volumes *m*, the population of active molecules ∑_*n*_ *b_n_*, and the location of the molecule n through time 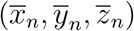. In this study we follow the Bayesian paradigm,^10,22,39^ and in particular the Bayesian nonparametric paradigm, to estimate variables of interest described above.

The variables *b_n_*, can take values 0 or 1 for each model molecule. Thus, each *b_n_* is a Bernoulli random variable. Here *b_n_* = 0 is associated with an inactive molecule *n* which does not contribute to the observations. If the *n^th^* molecule contributes photons at any point during the observation trace, then it is associated with *b_n_* = 1 and termed active. Within the purview of nonparametrics lies our ability to introduce an arbitrarily large number of molecules, each associated with its own *b_n_*, and determine which of these the data warrants as “active” versus “inactive”.

Within the Bayesian paradigm, whether parametric or nonparametric, we need to define priors over all unknowns (designated by blue circles in the graphical model of Fig. 6) and our choices over priors are detailed in the SI Sec. S.4.1.

Once the choices for the priors over random variables of (*D*, 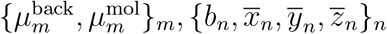) are made, we form a joint posterior probability 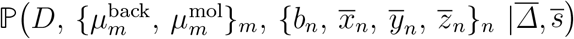 encompassing all unknown variables which we may wish to determine.

On account of the complexity of our posterior with respect to variables *D*, 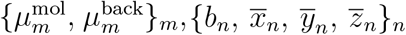, our posterior does not assume an analytic form. For this reason, we develop a computational scheme exploiting Markov chain Monte Carlo^38,39^ that can be used to generate pseudo-random samples from this posterior. Our MCMC exploits a Gibbs sampling scheme.^38–40^ Accordingly, posterior samples are generated by updating each one of the variables involved sequentially by sampling conditioned on all other variables and measurements 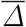. Conceptually, the steps involved in the generation of each posterior sample *D*, 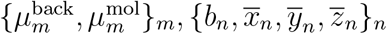 are:

1. Sampling the trajectories 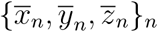 for *n* = 1,…, *N*
2. Sampling the diffusion coefficient *D*
3. Joint sampling the molecular brightnesses 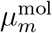 and background photon emission rates 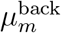 for *m* = 1,…, *M*
4. Sampling the molecular loads *b_n_* for *n* = 1,…, *N*.

To achieve step (1) without approximations on the forms of Eqs. (1) and (2), we use a Hamiltonian Monte Carlo (HMC) sampling scheme.^39,43-45^ This is by contrast to our previous work,^11–13^ where we exploited approximate Kalman filters^46–52^ to infer the molecular positions. Step (2) can be achieved analytically by virtue of the conjugacy of the prior to the marginal likelihood, steps (3) is achieved by a brute-force Metropolis scheme, and step (4) can be achieved by direct sampling. More details can be found in SI S.5.

A full detailed mathematical formulation and computational implementation of our proposed model, based on the multi-focus confocal microscope, is explained in the SI Sec. S.4 and S.5.

## Acknowledgement

S.P. thank NIH NIGMS (R01GM130745) for supporting Ioannis Sgouralis (early efforts in nonparametrics and tracking) and NIH NIGMS (R01GM134426) for supporting Sina Jazani (single photon efforts).

## Supporting Information Available

The following files are available free of charge.

- Filename: Supporting information
- Filename: Source code and GUI

