## Supporting Information for "A computational proposal for tracking multiple molecules in a multi-focus confocal setup"

Sina Jazani
*Department of Biophysics and Biophysical Chemistry*
*Johns Hopkins Medicine, Baltimore*
*Center for Biological Physics, Department of Physics*
*Arizona State University, Tempe*

Lance W.Q. Xu (徐伟青) and Douglas Shepherd
*Center for Biological Physics, Department of Physics*
*Arizona State University, Tempe*

Ioannis Sgouralis
*Department of Mathematics*
*University of Tennessee, Knoxville*
*Center for Biological Physics, Department of Physics*
*Arizona State University, Tempe*

Steve Pressé
*Center for Biological Physics,*
*Department of Physics and School of Molecular Sciences*
*Arizona State University, Tempe*

In these supplementary materials we discuss: (i) additional results that illustrate the potential of our method to estimate diffusion coefficient, molecular brightnesses and background photon emission rates. (ii) summary of point estimates results. (iii) details of the methods including descriptions of the motion model, point spread functions (PSFs), trajectory selection. (iv) a complete description of the inference framework developed that includes choices for the prior probability distributions. (v) a description of the computational implementation of the model. (vi) summary of notation and other conventions used throughout this study as well as detailed parameter choices for the simulations and analyses.

### Contents

S1. Additional results 3

S2. Summary of point estimates 11

S3. Detailed methods description 12
S3.1. Description of frame of reference 12
S3.2. Definition of molecular brightness 12
S3.3. Definition of point spread function models 12
S3.4. Description of the data simulation 13
S3.5. Description of the time trace preparation 14

S4. Detailed description of the inference framework 15
S4.1. Description of prior probability distributions 15
S4.1.1. Prior on the diffusion coefficient 15
S4.1.2. Priors on molecular brightness and background photon emission rates 15
S4.1.3. Priors on initial molecule locations 15
S4.1.4. Priors and hyperpriors for molecule loads 15
S4.2. Summary of our model 17

S5. Description of the computational scheme 17
S5.1. Overview of the sampling updates 18
S5.2. Sampling of active molecule trajectories 18
S5.2.1. Posterior transformation to a Hamiltonian 19
S5.2.2. Perform the Strang-splitting algorithm to solve Hamilton's equations 21
S5.2.3. Perform a Metropolis-Hastings test to accept or reject the proposed sample 24
S5.3. Sampling of inactive molecule trajectories 25
S5.4. Sampling the diffusion coefficient 25
S5.5. Sampling the molecule loads 26
S5.6. Joint sampling of molecular brightness and background photon emission rates 27
S5.7. Track the loads and the trajectories of molecules 28
S5.8. Maximum posteriori (MAP) sample 30

S6. Summary of notation, abbreviations, parameters and other quantities 33

### S1. Additional results

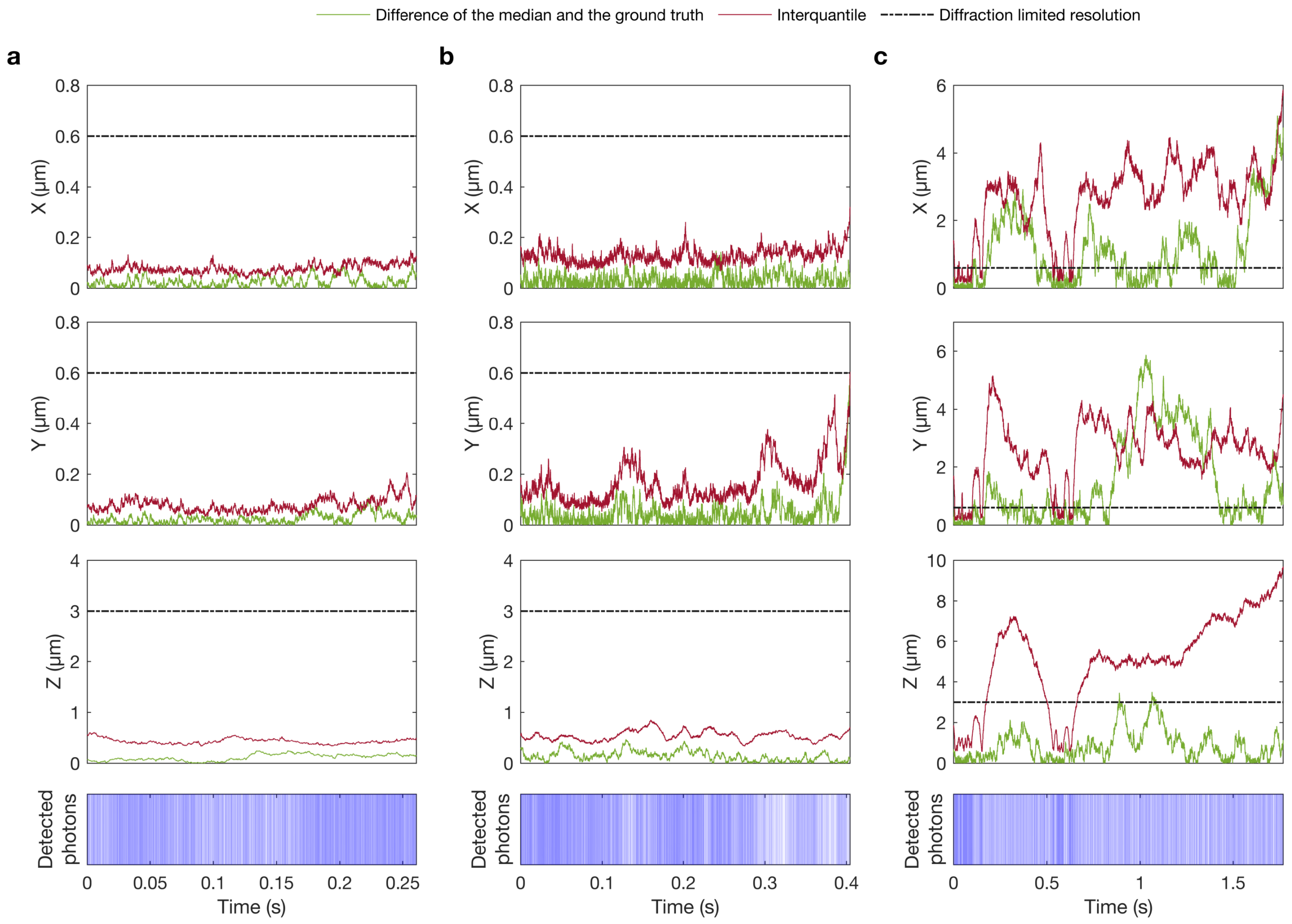


FIG. S1. **Accuracy and precision of the trajectory estimate. (a-c)** The difference of the median of the posterior and the ground truth as the accuracy (shown with green) and interquartile of the posterior as the precision (shown with maroon) for all coordinates associated with the synthetic single photon arrival time traces used in Fig. 2.

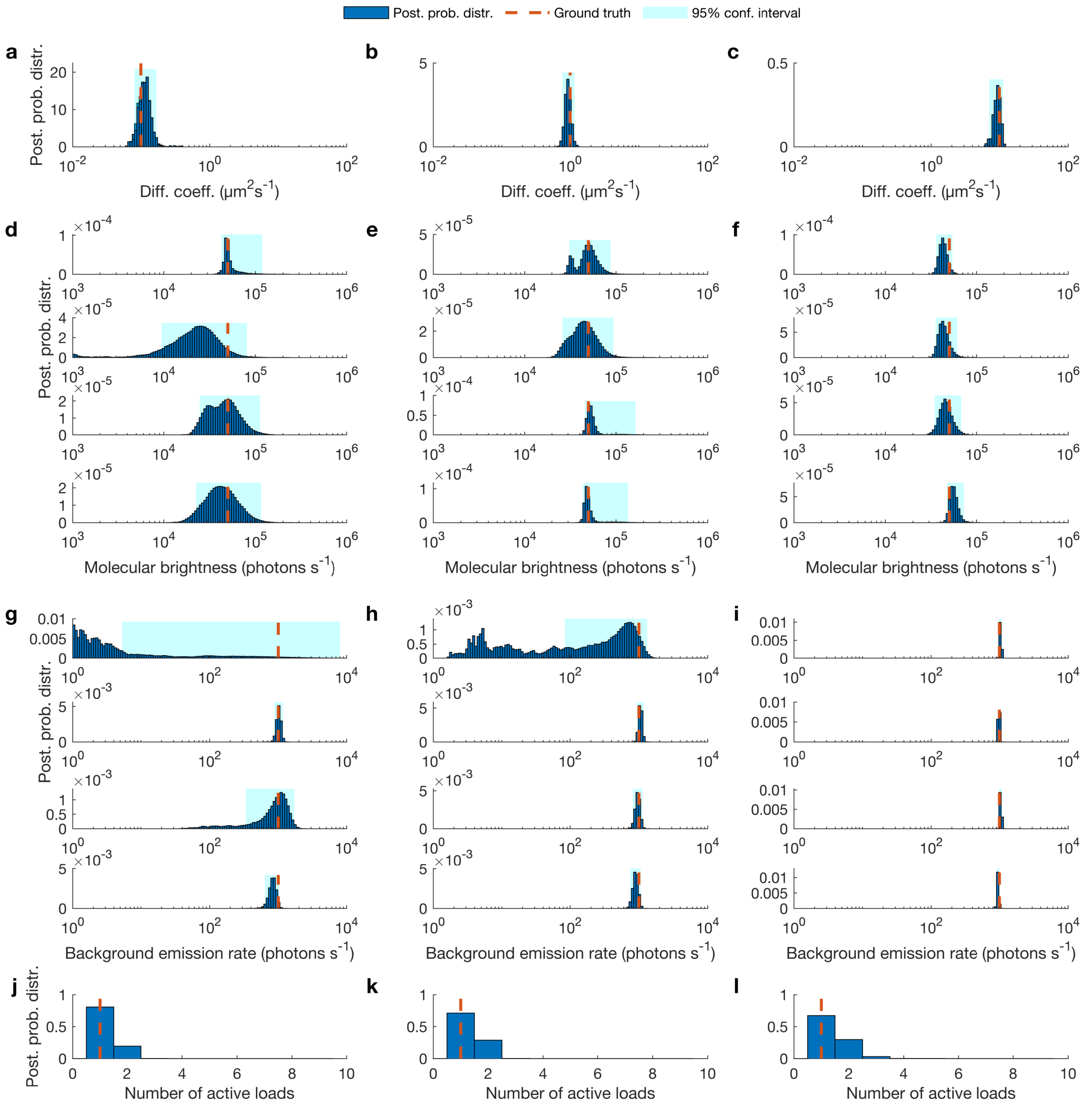


FIG. S2. **Estimated posterior probability distributions of the diffusion coefficients, molecular brightnesses and background photon emission rates. (a-c)** Posterior probability distributions of the diffusion coefficients associated with the synthetic single photon arrival time traces used in Fig. 2. **(d-f)** Posterior probability distributions of the molecular brightnesses associated with those same synthetic fluorescent intensity traces. **(g-i)** Posterior probability distributions of the background photon emission rates for those same traces. **(j-l)** Posterior probability distributions of the number of active molecules for those same traces.

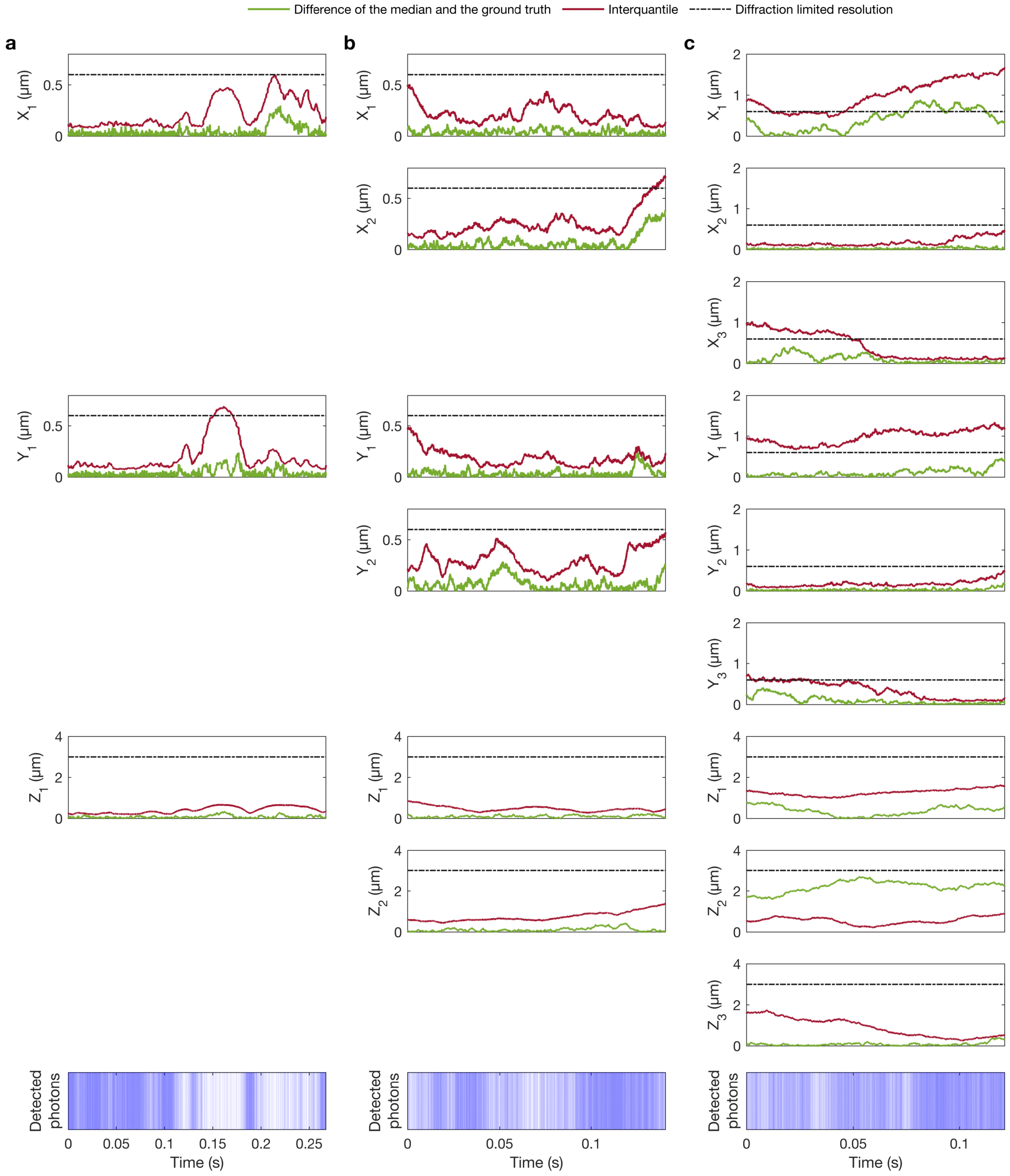


FIG. S3. **Accuracy and precision of the trajectory estimate. (a-c)** The difference of the median of the posterior and the ground truth as the accuracy (shown with green) and interquartile of the posterior as the precision (shown with maroon) for all coordinates associated with the synthetic single photon arrival time traces used in Fig. 3.

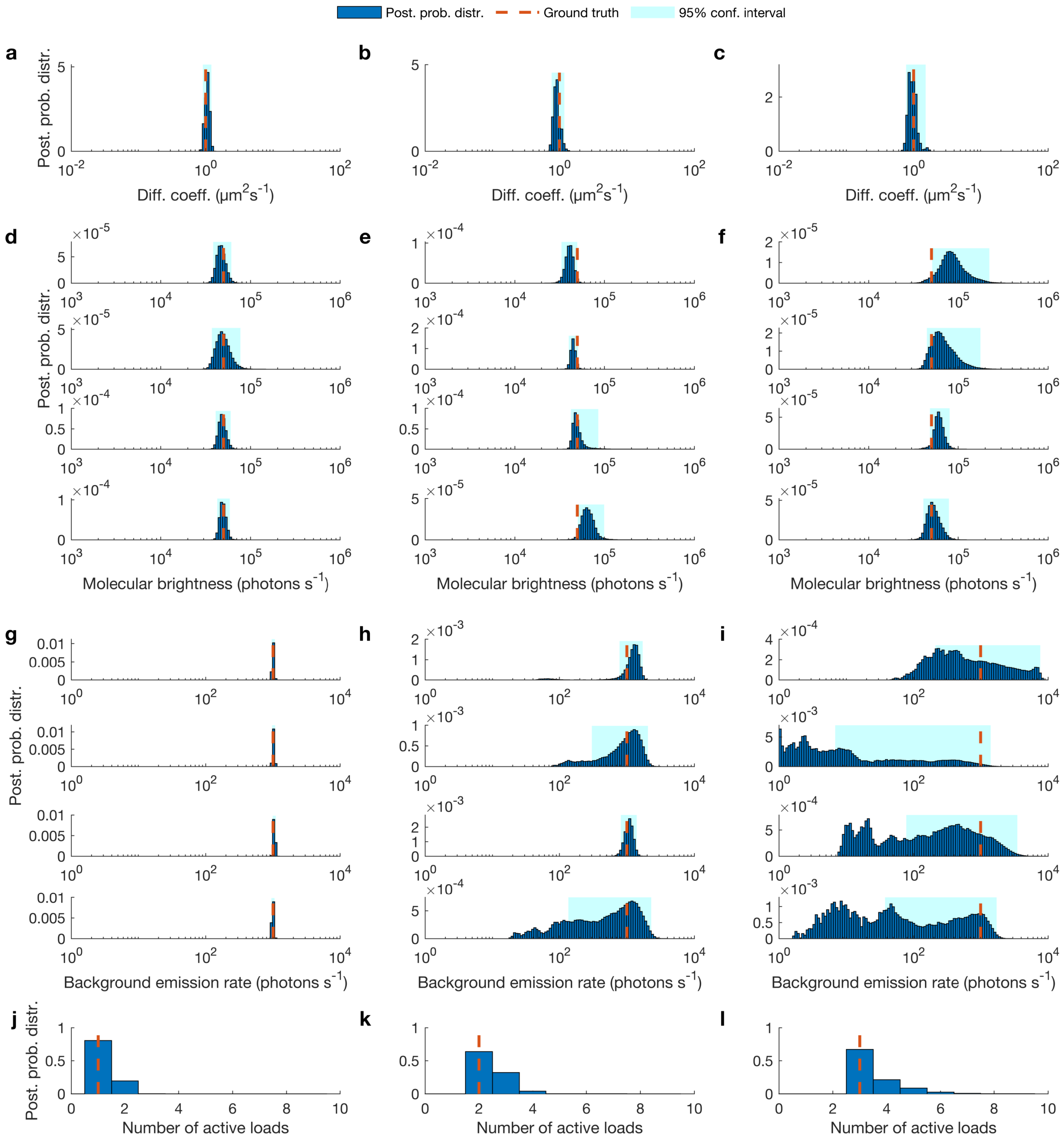


FIG. S4. **Estimated posterior probability distributions of the diffusion coefficients, molecular brightnesses and background photon emission rates.** **(a-c)** Posterior probability distributions of the diffusion coefficients related to synthetic fluorescent intensity trace used in Fig. 3. **(d-f)** Posterior probability distributions of the molecular brightnesses associated with those same synthetic fluorescent intensity traces. **(g-i)** Posterior probability distributions of the background photon emission rates for those same traces. **(j-l)** Posterior probability distributions of the number of active molecules for those same traces.

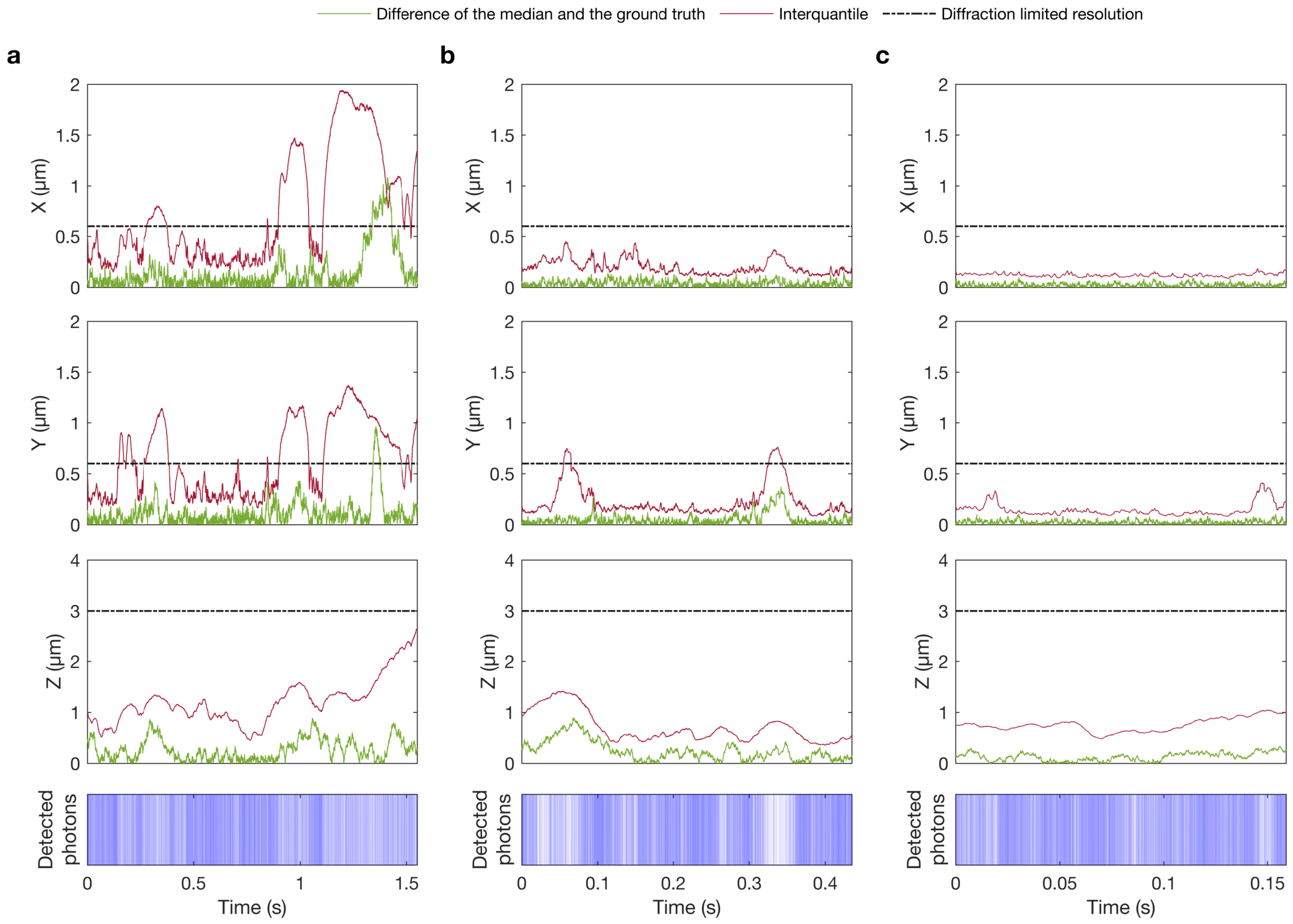


FIG. S5. **Accuracy and precision of the trajectory estimate. (a-c)** The difference of the median of the posterior and the ground truth as the accuracy (shown with green) and interquartile of the posterior as the precision (shown with maroon) for all coordinates associated with the synthetic single photon arrival time traces used in Fig. 4.

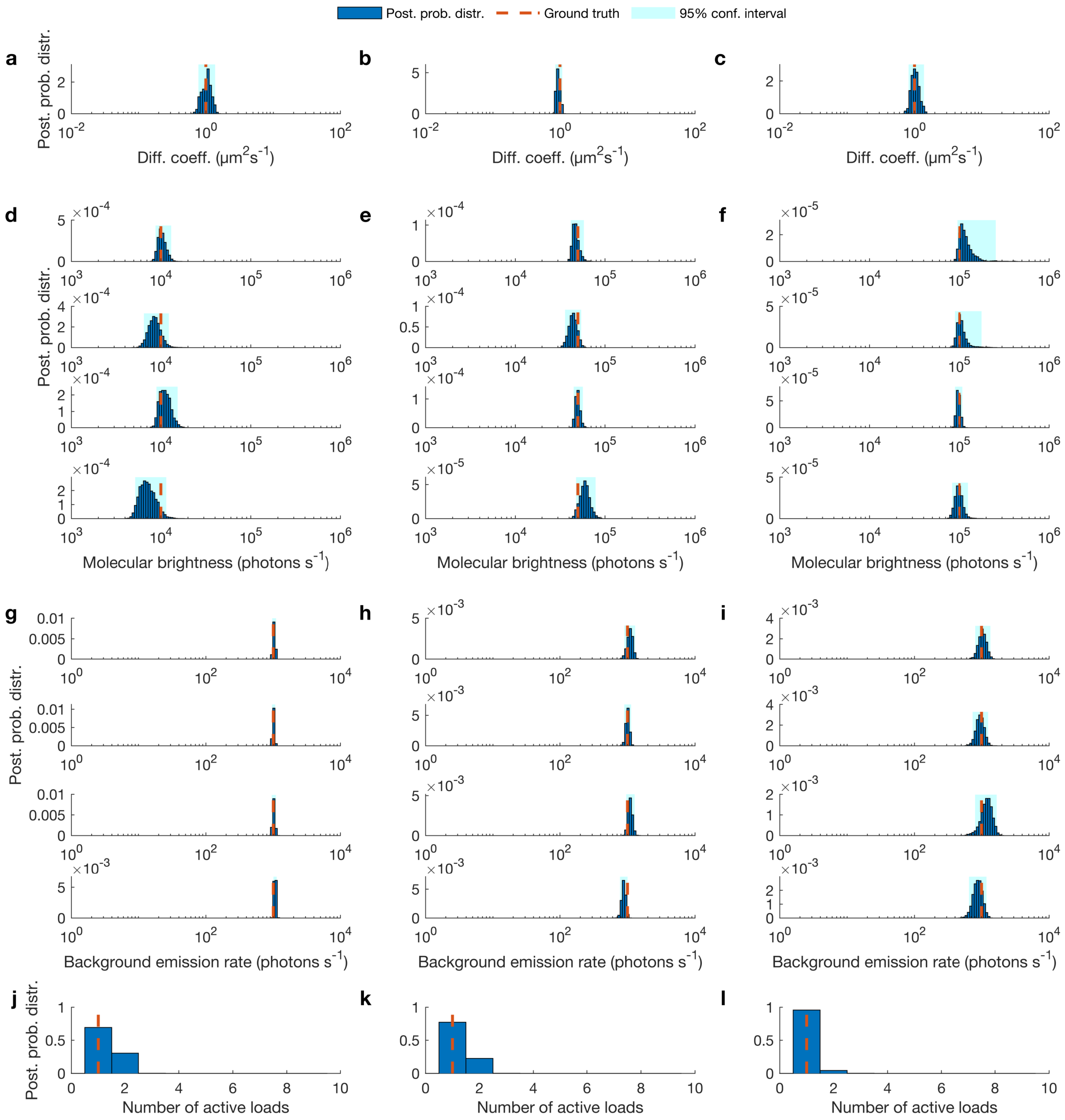


FIG. S6. **Estimated posterior probability distributions of the diffusion coefficients, molecular brightnesses and background photon emission rates.** **(a-c)** Posterior probability distributions of the diffusion coefficients related to synthetic fluorescent intensity trace used in Fig. 4. **(d-f)** Posterior probability distributions of the molecular brightnesses associated with those same synthetic fluorescent intensity traces. **(g-i)** Posterior probability distributions of the background photon emission rates for those same traces. **(j-l)** Posterior probability distributions of the number of active molecules for those same traces.

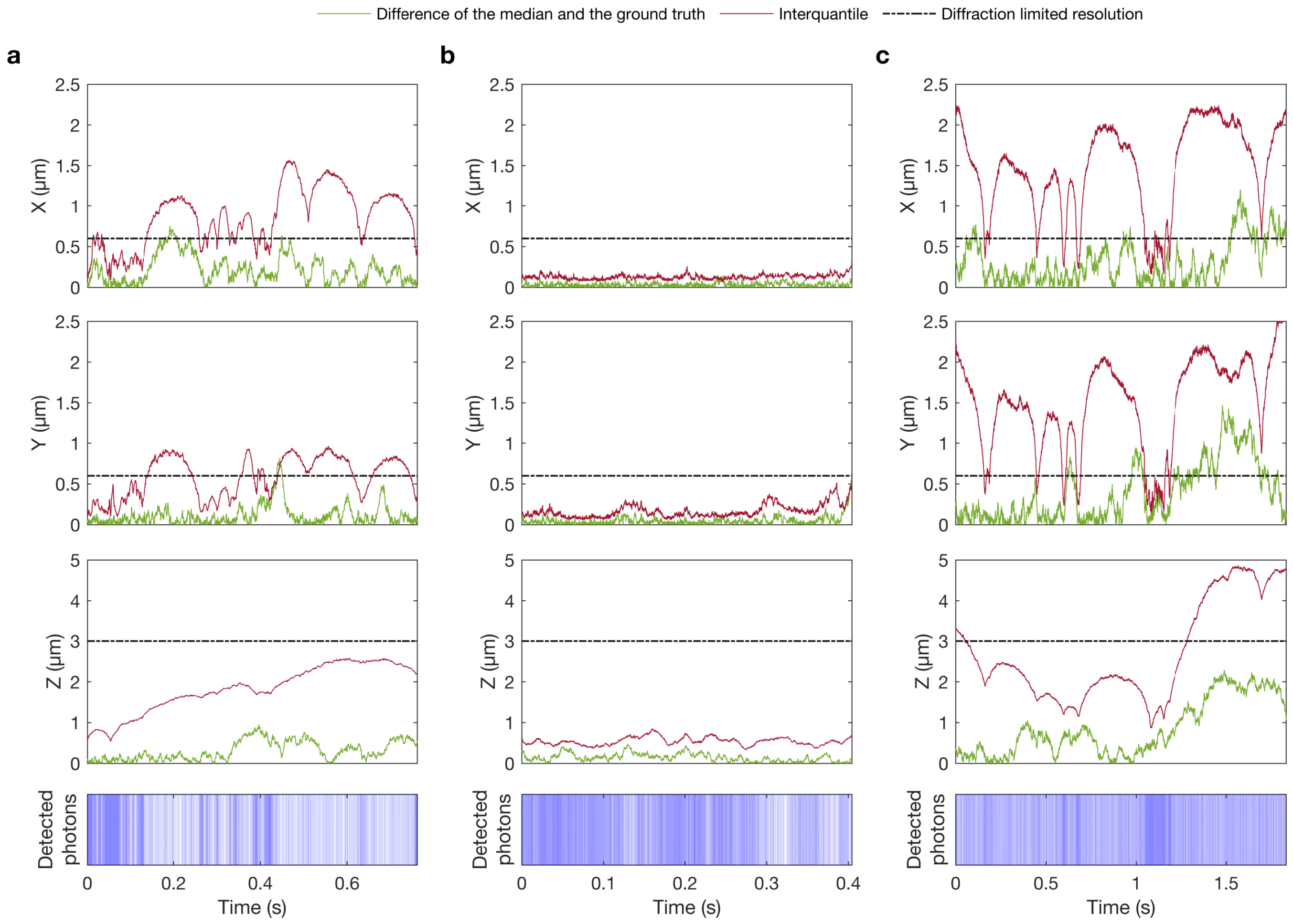


FIG. S7. **Accuracy and precision of the trajectory estimate. (a-c)** The difference of the median of the posterior and the ground truth as the accuracy (shown with green) and interquartile of the posterior as the precision (shown with maroon) for all coordinates associated with the synthetic single photon arrival time traces used in Fig. 5.

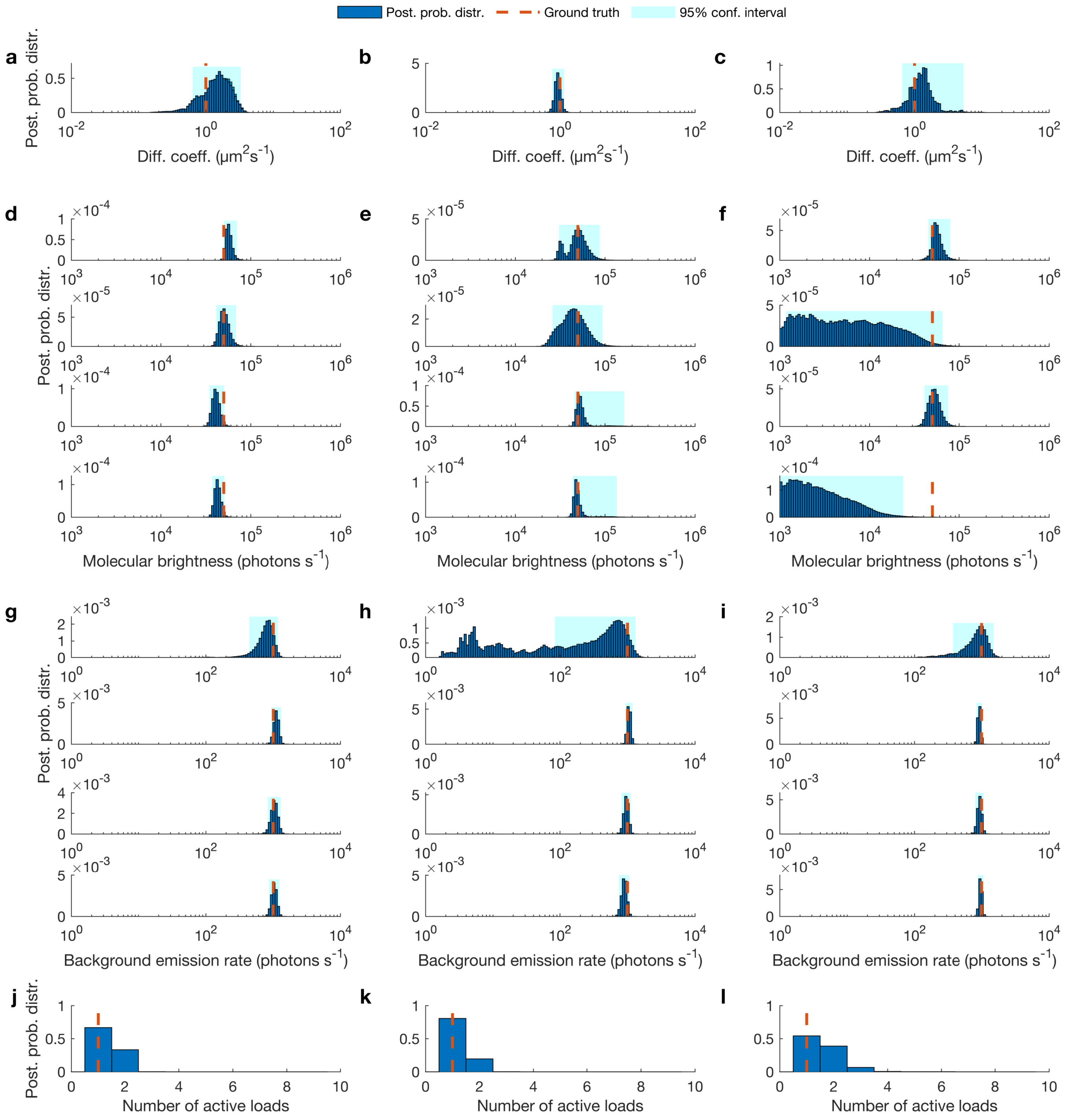


FIG. S8. **Estimated posterior probability distributions of the diffusion coefficients, molecular brightnesses and background photon emission rates.** **(a-c)** Posterior probability distributions of the diffusion coefficients related to synthetic fluorescent intensity trace used in Fig. 5. **(d-f)** Posterior probability distributions of the molecular brightnesses associated with those same synthetic fluorescent intensity traces. **(g-i)** Posterior probability distributions of the background photon emission rates for those same traces. **(j-l)** Posterior probability distributions of the number of active molecules for those same traces.

### S2. Summary of point estimates

TABLE S1. Here, we list characteristic values (point estimates) of the posterior probability distributions of the diffusion coefficients $D$, molecular brightnesses $\mu_m^{\text{mol}}$ and background photon emission rates $\mu_m^{\text{back}}$ of all examples shown in this study. Mean and std refer to mean value and standard deviation of the posterior (i.e., the square root of variance). Since, in this study we consider four confocal volumes $m = 1, 2, 3, 4$, there are four molecular brightnesses and background photon emission rates for each figure.

| | $D$ ($\mu$m$^2$s$^{-1}$) | | $\mu_m^{\text{mol}}$ (photons s$^{-1}$) | | $\mu_m^{\text{back}}$ ( photons s$^{-1}$) | |
|---|---|---|---|---|---|---|
| | mean | std | mean | std | mean | std |
| Fig. 2(a) | 0.13 | 0.07 | $\begin{bmatrix}5.65\\3.23\\5.50\\5.43\end{bmatrix} \times 10^4$ | $\begin{bmatrix}2.03\\1.84\\2.29\\2.40\end{bmatrix} \times 10^4$ | $\begin{bmatrix}2.10\\1.02\\1.08\\0.84\end{bmatrix} \times 10^3$ | $\begin{bmatrix}24.1\\0.75\\3.43\\0.99\end{bmatrix} \times 10^2$ |
| Fig. 2(b) | 0.96 | 0.13 | $\begin{bmatrix}5.31\\5.07\\6.43\\5.82\end{bmatrix} \times 10^4$ | $\begin{bmatrix}1.51\\1.81\\3.06\\2.52\end{bmatrix} \times 10^4$ | $\begin{bmatrix}0.72\\1.06\\0.97\\0.90\end{bmatrix} \times 10^3$ | $\begin{bmatrix}3.99\\0.63\\0.78\\0.81\end{bmatrix} \times 10^2$ |
| Fig. 2(c) | 9.37 | 2.12 | $\begin{bmatrix}4.61\\5.12\\4.65\\5.72\end{bmatrix} \times 10^4$ | $\begin{bmatrix}1.24\\2.52\\0.90\\0.68\end{bmatrix} \times 10^4$ | $\begin{bmatrix}1.04\\0.99\\1.04\\0.95\end{bmatrix} \times 10^3$ | $\begin{bmatrix}0.37\\0.25\\0.27\\0.24\end{bmatrix} \times 10^2$ |
| Fig. 3(a) | 1.07 | 0.08 | $\begin{bmatrix}4.78\\5.14\\5.35\\5.42\end{bmatrix} \times 10^4$ | $\begin{bmatrix}0.62\\1.04\\3.05\\3.08\end{bmatrix} \times 10^4$ | $\begin{bmatrix}1.01\\1.02\\1.04\\0.99\end{bmatrix} \times 10^3$ | $\begin{bmatrix}0.33\\0.28\\0.29\\0.33\end{bmatrix} \times 10^2$ |
| Fig. 3(b) | 0.94 | 0.11 | $\begin{bmatrix}4.14\\4.52\\5.23\\6.83\end{bmatrix} \times 10^4$ | $\begin{bmatrix}0.43\\0.28\\1.30\\1.43\end{bmatrix} \times 10^4$ | $\begin{bmatrix}1.29\\1.21\\1.10\\1.17\end{bmatrix} \times 10^3$ | $\begin{bmatrix}0.24\\0.45\\0.15\\0.57\end{bmatrix} \times 10^3$ |
| Fig. 3(c) | 1.03 | 0.28 | $\begin{bmatrix}1.03\\0.81\\0.62\\0.55\end{bmatrix} \times 10^5$ | $\begin{bmatrix}4.46\\3.58\\0.80\\1.04\end{bmatrix} \times 10^4$ | $\begin{bmatrix}3.51\\0.55\\1.33\\0.84\end{bmatrix} \times 10^3$ | $\begin{bmatrix}2.42\\0.40\\1.01\\0.47\end{bmatrix} \times 10^3$ |
| Fig. 4(a) | 1.06 | 0.16 | $\begin{bmatrix}1.05\\0.89\\1.14\\0.76\end{bmatrix} \times 10^4$ | $\begin{bmatrix}1.26\\1.55\\1.76\\2.17\end{bmatrix} \times 10^3$ | $\begin{bmatrix}1.03\\1.01\\1.02\\1.06\end{bmatrix} \times 10^3$ | $\begin{bmatrix}0.42\\0.30\\0.64\\0.31\end{bmatrix} \times 10^2$ |
| Fig. 4(b) | 0.92 | 1.12 | $\begin{bmatrix}4.79\\4.53\\4.94\\5.97\end{bmatrix} \times 10^4$ | $\begin{bmatrix}5.87\\6.77\\3.22\\10.2\end{bmatrix} \times 10^3$ | $\begin{bmatrix}1.10\\0.99\\1.09\\0.87\end{bmatrix} \times 10^3$ | $\begin{bmatrix}1.18\\1.08\\1.14\\0.82\end{bmatrix} \times 10^2$ |
| Fig. 4(c) | 1.03 | 0.14 | $\begin{bmatrix}1.27\\1.09\\0.99\\1.01\end{bmatrix} \times 10^5$ | $\begin{bmatrix}4.09\\2.23\\0.67\\1.12\end{bmatrix} \times 10^4$ | $\begin{bmatrix}1.06\\1.03\\1.22\\0.91\end{bmatrix} \times 10^3$ | $\begin{bmatrix}1.41\\7.97\\2.60\\1.42\end{bmatrix} \times 10^2$ |
| Fig. 5(a) | 1.92 | 0.77 | $\begin{bmatrix}5.76\\5.21\\4.16\\4.31\end{bmatrix} \times 10^4$ | $\begin{bmatrix}5.10\\6.93\\4.13\\3.54\end{bmatrix} \times 10^3$ | $\begin{bmatrix}0.83\\1.12\\1.05\\1.04\end{bmatrix} \times 10^3$ | $\begin{bmatrix}1.85\\0.95\\1.19\\0.94\end{bmatrix} \times 10^2$ |
| Fig. 5(b) | 0.96 | 0.14 | $\begin{bmatrix}5.31\\5.07\\6.43\\5.83\end{bmatrix} \times 10^4$ | $\begin{bmatrix}1.51\\1.81\\3.06\\2.52\end{bmatrix} \times 10^4$ | $\begin{bmatrix}0.72\\1.06\\0.97\\0.90\end{bmatrix} \times 10^3$ | $\begin{bmatrix}3.99\\0.63\\0.78\\0.81\end{bmatrix} \times 10^2$ |
| Fig. 5(c) | 3.03 | 6.45 | $\begin{bmatrix}5.76\\2.08\\5.41\\0.64\end{bmatrix} \times 10^4$ | $\begin{bmatrix}1.09\\1.72\\0.87\\0.72\end{bmatrix} \times 10^4$ | $\begin{bmatrix}1.18\\0.92\\0.95\\0.97\end{bmatrix} \times 10^3$ | $\begin{bmatrix}12.8\\0.45\\1.16\\0.47\end{bmatrix} \times 10^2$ |

### S3. Detailed methods description

#### S3.1. Description of frame of reference

The PSFs we consider in this study are created by the overlap of the detection profile with the excitation profile. It can be seen in Fig. S9(a-b) how $\mathrm{PSF}_m$, $m = 1, \ldots, M$ can be considered as the result of the detection profile and the excitation profile. As with regular FCS, the dimension of each $\mathrm{PSF}_m$ can be calibrated by considering a well known molecule with a known diffusion coefficient. [1–3] Also, as we illustrate in Fig. S9(c), to coordinate all PSFs, we define a global frame of reference where the point of origin is placed at (0,0,0) and each one of the PSFs are centered at distance $(\mathrm{C}_{m,x}, \mathrm{C}_{m,y}, \mathrm{C}_{m,z})$ from the point of origin.

#### S3.2. Definition of molecular brightness

Here, we use the term "molecular brightness" for the random variables $\mu_m^{\mathrm{mol}}$. This random variable the is the same as in our previous work on a single-focus confocal volume. [4] Specifically

$$\mu_m\,(x, y, z) = \mu_{\mathrm{exc}}\ \varphi_{\mathrm{d}}\ \varphi_{m,\mathrm{qe}}\ \varphi_{\mathrm{f}}\ \sigma\ \mathrm{EXC}(x, y, z)\ \mathrm{CEF}(x, y, z) \tag{S1}$$

where, $\mu_{\mathrm{exc}}$ is the maximum excitation intensity which occurs at the center of the excitation profile, $\varphi_{m,\mathrm{d}}$ is the efficiency of the photon collection at the center of the detection profile of detector $m$, $\varphi_{\mathrm{qe}}$ is the quantum efficiency of the detector $m$, $\varphi_{\mathrm{f}}$ is the quantum efficiency of the fluorophore (i.e. quantum yield), $\sigma$ is the absorption cross-section of the fluorophore, $\mathrm{EXC}\,(x, y, z)$ is the excitation profile and $\mathrm{CEF}_m\,(x, y, z)$ is the detection profile of detector $m$, i.e., collection efficiency function, which equals the fraction of the photons collected by the detector $m$ to the total photons emitted by a point source. [5]

To obtain Eq. (3), we cast Eq. (S1) in the simplified form

$$\mu_m(x, y, z) = \mu_m^{\mathrm{mol}}\,\mathrm{PSF}_m\,(x, y, z) \tag{S2}$$

where $\mu_m^{\mathrm{mol}} = \mu_{\mathrm{exc}}\,\varphi_{m,\mathrm{d}}\,\varphi_{m,\mathrm{qe}}\,\varphi_{\mathrm{f}}\,\sigma$, which we term molecular brightness at the center of the confocal volume $m$, [6] and $\mathrm{PSF}_m\,(x, y, z) = \mathrm{EXC}\,(x, y, z)\ \mathrm{CEF}_m\,(x, y, z)$, which we term the PSF.

#### S3.3. Definition of point spread function models

There are several PSF models one could consider. For example: the Airy function, [7–9] 3D-Gaussian (3DG), [10] 2D-Gaussian-Lorentzian (2DGL) [11–14] and 2D-Gaussian-Cylindrical (2DGC). [10] Our work is general and can accommodate any of these points spread functions (PSFs) or combinations thereof.

The definition of the 3DG PSF is

$$\mathrm{PSF}_{m,\mathrm{3DG}}(x, y, z) = \exp\left(-2\left(\left(\frac{x - \mathrm{C}_{m,x}}{\omega_{m,x}}\right)^2 + \left(\frac{y - \mathrm{C}_{m,y}}{\omega_{m,y}}\right)^2 + \left(\frac{z - \mathrm{C}_{m,z}}{\omega_{m,z}}\right)^2\right)\right) \tag{S3}$$

while that of the 2DGL PSF is

$$\mathrm{PSF}_{m,\mathrm{2DGL}}\,(x, y, z) = \frac{1}{1 + \left(\frac{z - \mathrm{C}_{m,z}}{\omega_{m,z}}\right)^2} \exp\left(-2\left(\frac{\left(\frac{x - \mathrm{C}_{m,x}}{\omega_{m,x}}\right)^2 + \left(\frac{y - \mathrm{C}_{m,y}}{\omega_{m,y}}\right)^2}{1 + \left(\frac{z - \mathrm{C}_{m,z}}{\omega_{m,z}}\right)^2}\right)\right). \tag{S4}$$

Finally, that of the 2DGC PSF is

$$\mathrm{PSF}_{m,\mathrm{2DGCZ}}\,(x, y, z) = \exp\left(-2\left(\left(\frac{x - \mathrm{C}_{m,x}}{\omega_{m,x}}\right)^2 + \left(\frac{y - \mathrm{C}_{m,y}}{\omega_{m,y}}\right)^2\right)\right) \tag{S5}$$

where $(\mathrm{C}_{m,x}, \mathrm{C}_{m,y}, \mathrm{C}_{m,z})$ is the coordinate of the center of the $\mathrm{PSF}_m$ from the point of origin and $(\omega_{m,x}, \omega_{m,y}, \omega_{m,z})$ are the semi-axes lateral and parallel to the optical axis of the confocal volume m. Here, the widths of the confocal volumes $(\omega_{m,x},\ \omega_{m,y},\ \omega_{m,z})$ can be directly estimated by calibration experiments with known diffusion coefficients; for example see Ref. [15].

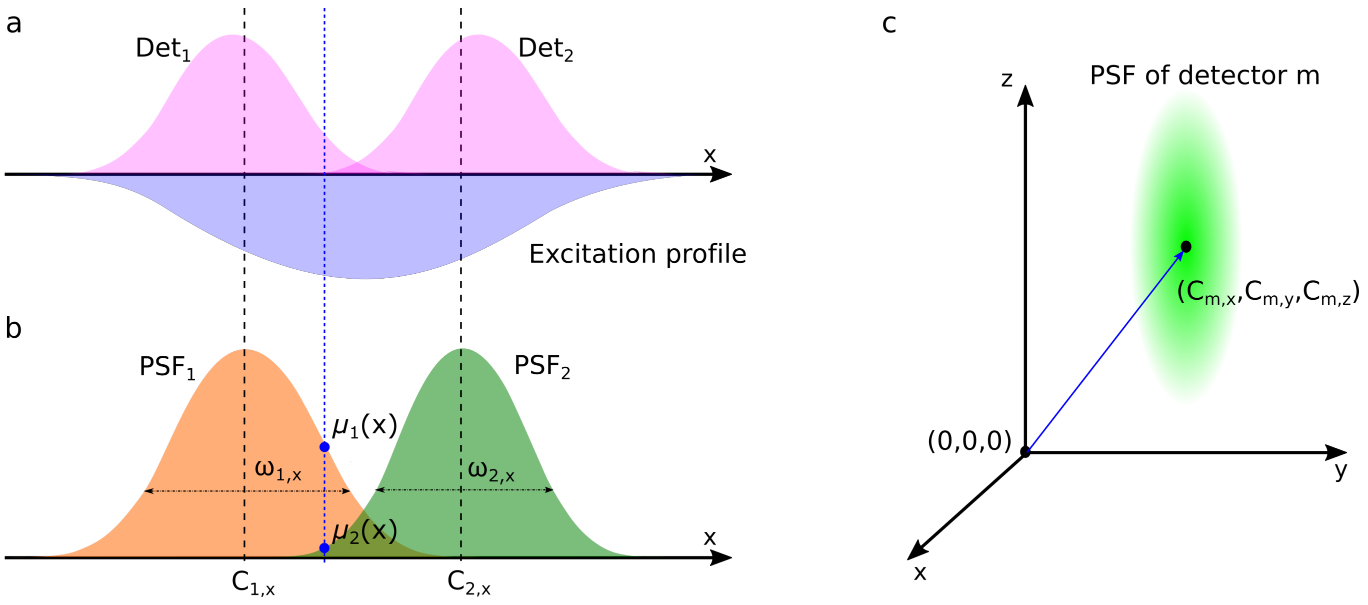


FIG. S9. **Illustration of a multi-focus confocal setup.** **(a)** The excitation profile is represented by the inverted blue Gaussian while the detection volumes ($\mathrm{Det}_1$ and $\mathrm{Det}_2$) are represented by the pink curves in 1D coordinate. **(b)** The overlap in the detection volume and illumination volumes produce two PSFs in 1D coordinate that we term "confocal volume". Each PSF has its own mean and standard deviation denoted by $\mathrm{C}_{1,x}$ and $\mathrm{C}_{2,x}$ as mean values and $\frac{\omega_{1,x}}{2}$ and $\frac{\omega_{2,x}}{2}$ as standard deviations . **(c)** The global frame of reference which shows the point of origin and position of PSFs in 3D.

#### S3.4. Description of the data simulation

To generate single photon arrival time traces mimicking a realistic multi-focus confocal setup, we simulate molecules moving [16, 17] through multi-illuminated 3D volumes. The number of diffusing molecules, $N$, is prescribed in each simulation. To maintain a relatively stable concentration of molecules near the confocal volume, we impose periodic rectangular boundaries. The boundaries are placed at $\pm L_x$ and $\pm L_y$ perpendicular to the focal plane and $\pm L_z$ perpendicular to the optical axis. We typically choose L large enough such that it be larger than the width of the combinations of PSFs

$$L_x \gg |\max\left(\overline{\mathrm{C}_x}\right) - \min\left(\overline{\mathrm{C}_x}\right)| + \max\left(\overline{\omega_x}\right) \tag{S6}$$

$$L_y \gg |\max\left(\overline{\mathrm{C}_y}\right) - \min\left(\overline{\mathrm{C}_y}\right)| + \max\left(\overline{\omega_y}\right) \tag{S7}$$

$$L_z \gg |\max\left(\overline{\mathrm{C}_z}\right) - \min\left(\overline{\mathrm{C}_z}\right)| + \max\left(\overline{\omega_z}\right) \tag{S8}$$

where $\left(\overline{C}_x, \overline{C}_y, \overline{C}_z\right)$ and $(\overline{\omega}_x, \overline{\omega}_y, \overline{\omega}_z)$ are the coordinate of the centers from point of origin and dimensions of confocal volumes. For simulation purposes, we assess the locations of the molecules $(x_{n,k},\ y_{n,k},\ z_{n,k})$, where $k = 1, \ldots, K$ label time levels and $n = 1, \ldots, N$ label molecules, at equidistant time intervals $t_1, t_2, \ldots, t_K$. The time interval $\delta t$, however, is smaller than the time interval between successive assessments $\varDelta_k = t_{k+1} - t_k$ and the total trace duration $T_{\text{total}} = t_K - t_0$.

Molecule locations at the first assessment $(x_{n,1},\ y_{n,1},\ z_{n,1})$ are sampled randomly from a uniform distribution with limits equal to the boundaries $\pm L_x$, $\pm L_y$ and $\pm L_z$ of our pre-specified volume. Subsequent locations are sampled according to the diffusion model which we considered it as a Brownian motion, [4, 18, 19] under a prescribed diffusion coefficient $D$.

Finally, we obtain individual photon inter-arrival times, $\varDelta_k$, from any of the confocal volumes and the detector, $s_k$ which can take value 1 to $M$, at which the $k^{\text{th}}$ photons is detected as follows

$$\varDelta_k \sim \mathbf{Exp}\left(\sum_{m=1}^{M} \mu_{m,k}\right) \tag{S9}$$

$$s_k \sim \mathbf{Cat}_{1,\ldots,\mathrm{M}}\left(\left[\frac{\mu_{1,k}}{\sum_{m=1}^{M} \mu_{m,k}}, \ldots, \frac{\mu_{M,k}}{\sum_{m=1}^{M} \mu_{m,k}}\right]\right) \tag{S10}$$

where the rate $\mu_{m,k}$ gathers single photon contributions from the background and the entire molecule population according to

$$\mu_{m,k} = \mu_m^{\text{back}} + \mu_m^{\text{mol}} \sum_{n=1}^{N} \text{PSF}_m \left(x_{n,k}, y_{n,k}, z_{n,k}\right). \tag{S11}$$

Here, both background photon emission rate and molecular brightness, $\mu_m^{\text{back}}$ and $\mu_m^{\text{mol}}$, are prescribed and $\text{PSF}_m$ stands for any of the PSFs introduced in Eqs. (S3)(S4)(S5). As an example, in Fig. S9, we illustrate two Gaussian shape PSFs overlapping with each others in 1D. Detailed parameter choices for all simulations performed are listed in Table S6.

#### S3.5. Description of the time trace preparation

In real experiments, we envision each of the $M$ photon detectors to be simultaneously active. We label the time of arrival of photons at the $m^{\text{th}}$ detector as $t_{m,k}$, where $k' = 1, 2, \ldots, K'_m$. Our data, the starting point for our analysis, consists of the detector label at which each successive photon is detected, $\overline{s} = (s_1, \ldots, s_K)$, and the photon inter-arrival times obtained by combining all traces, $\overline{\Delta} = (\Delta_1, \ldots, \Delta_{K-1})$ where $\Delta_k = t_{k+1} - t_k$, $k = 1, \ldots, K-1$.

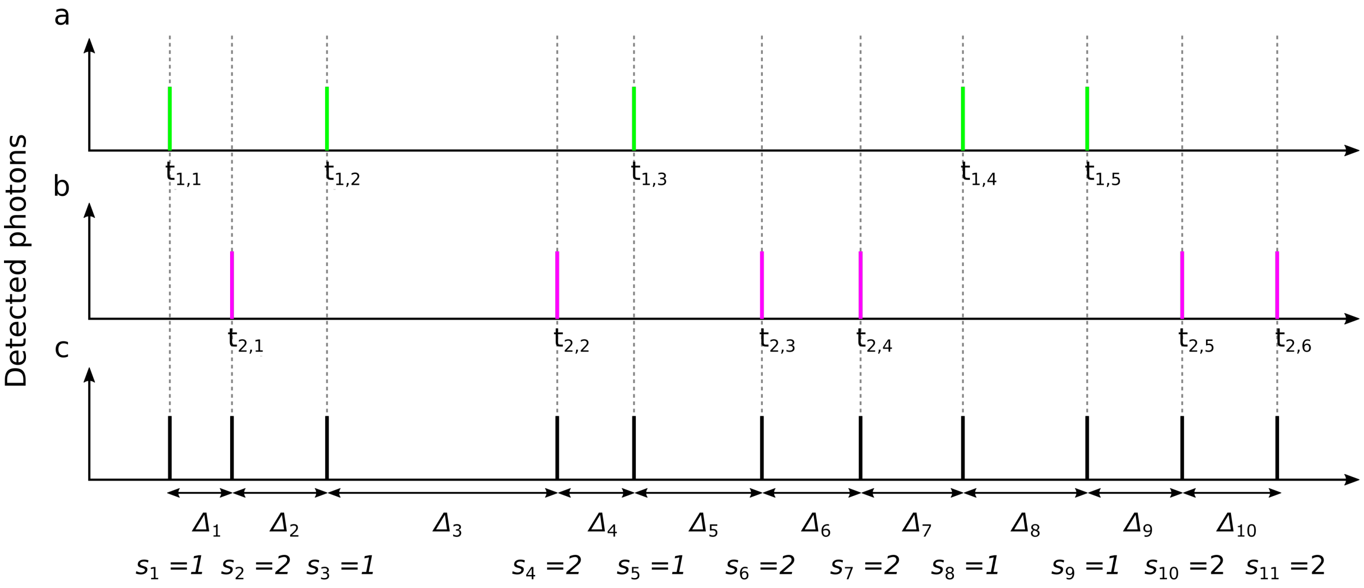


FIG. S10. **Illustration of time traces used in the analysis. (a-c)** Single photon time traces recorded by confocal volume one ($m$=1), two ($m$=2) and both ($m$=1,2), respectively. The inter-arrival times are denoted $\overline{\Delta}$ and $\overline{s}$ is a sequence of detector labels (with labels 1 through $M$ where $M = 2$ here for illustrative purposes) at which each photon was detected.

### S4. Detailed description of the inference framework

#### S4.1. Description of prior probability distributions

Model parameters in our framework requiring priors are: the diffusion coefficient $D$; the molecular brightness and background photon emission rates $\mu_m^{\text{mol}}$ and $\mu_m^{\text{back}}$; the initial molecule locations $x_{n,1}$,$y_{n,1}$,$z_{\text{n},1}$; and load $b_n$. As we will see, the load plays the role of an indicator variable which is either zero or one whether the molecule is or is not contributing photons. Our choices are described below.

##### S4.1.1. Prior on the diffusion coefficient

To ensure that $D$ sampled in our formulation attains only positive values, we place a inverse-Gamma prior

$$D \sim \textbf{InvGamma}\left(\alpha_D, \beta_D\right). \tag{S12}$$

Besides ensuring a positive $D$, this prior is conjugate to the motion model, captured by Eq. 4, which facilitates the computations (see below).

##### S4.1.2. Priors on molecular brightness and background photon emission rates

To ensure that $\mu_m^{\text{mol}}$ and $\mu_m^{\text{back}}$ sampled only attain positive values, we place Gamma priors on both

$$\begin{aligned} \mu_m^{\text{mol}} &\sim \textbf{Gamma}\left(\alpha_{\text{mol}}, \beta_{\text{mol}}\right) \\ \mu_m^{\text{back}} &\sim \textbf{Gamma}\left(\alpha_{\text{back}}, \beta_{\text{back}}\right). \end{aligned} \tag{S13}$$

Due to the specific dependencies of the likelihood (that we will discuss shortly) on the photon emission rates, conjugate priors cannot be achieved for $\mu_m^{\text{mol}}$ and $\mu_m^{\text{back}}$. So, the above choice offers no computational advantage (see below) and could be readily replaced with other distributions.

##### S4.1.3. Priors on initial molecule locations

Since, molecules can be anywhere in space, we place priors on the initial locations. To facilitate the computations (see below), we place independent symmetric normal distributions, on each Cartesian coordinate of the model molecule

$$x_{n,1} \sim \textbf{Normal}\left(\mu_{x_0}, \sigma_{x_0}^2\right) \tag{S14}$$

$$y_{n,1} \sim \textbf{Normal}\left(\mu_{y_0}, \sigma_{y_0}^2\right) \tag{S15}$$

$$z_{n,1} \sim \textbf{Normal}\left(\mu_{z_0}, \sigma_{z_0}^2\right). \tag{S16}$$

where ($\sigma_{x_0}$, $\sigma_{y_0}$, $\sigma_{\text{z}_0}$) denote the standard deviations (set to large values as compared to the size of the confocal volumes), and ($\mu_{x_0}$, $\mu_{y_0}$, $\mu_{z_0}$) denotes the mean which we choose to be centered at the origin.

##### S4.1.4. Priors and hyperpriors for molecule loads

In previous work, we explored the Beta-Bernoulli process used to determine how many molecules are contributing photons. [4, 18, 19] Briefly, we define a large population of molecules N which include both active and inactive molecules. These molecules are collectively indexed by $n = 1, 2, \ldots, N$. Estimating how many molecules are actually warranted by the data under analysis is equivalent to estimating how many of those N molecules are active, i.e., $b_n$=1 and emitting detected photons, while the remaining inactive ones, i.e., $b_n$=0, have no impact whatsoever and are instantiated only for computational purposes.

To ensure that each load $b_n$ takes only values 0 or 1, we place a Bernoulli prior of weight $q_n$ on $b_n$ and a conjugate Beta prior on $q_n$

$$q_n \sim \textbf{Beta}\left(\frac{\gamma_b}{N}, \frac{N-1}{N}\right) \tag{S17}$$
$$b_n|q_n \sim \textbf{Bernoulli}\left(q_n\right) \tag{S18}$$

where, n=1, ..., N. Under these choices, and in the limit that $N \to \infty$; that is, when the assumed molecule population is allowed to be large, this prior/hyperprior converge to the Beta-Bernoulli process, [20, 21] a novel mathematical tool that avoids having to pre-specify the number of active molecules by hand (as would be required within the traditional, parametric, Bayesian paradigm). Thanks to these new tools, even for $N \gg 1$, the posterior sharpens at the correct value of active molecules irrespective of how large we make $N$ initially. In other words, provided $N$ is large enough, our choice of $N$ is insignificant; while its precise value has only computational implications.

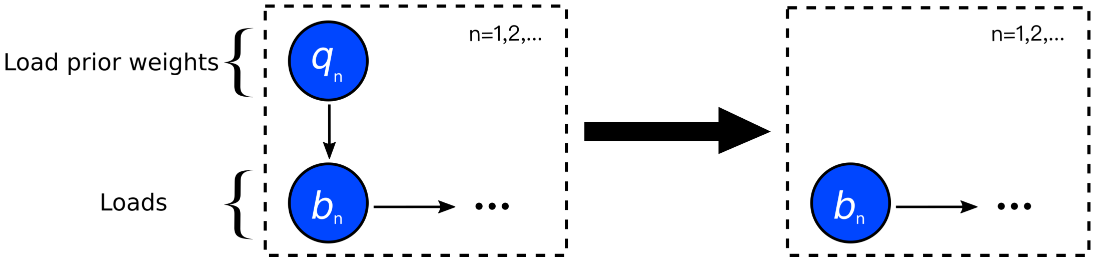


FIG. S11. **Graphical summary of the general framework developed for Beta-Bernoulli process.** A population of model molecules, labeled by $n = 1, 2, \ldots,$ load on each molecule is shown by $b_n$ and the weight on each load is by $q_n$.

As a way of simplification, and to avoid learning the $q_n$, we may marginalize over these as follows

$$\begin{aligned}
\mathbb{P}\left(b_n\right) &= \int_0^1 \mathbb{P}\left(b_n\right)\mathbb{P}\left(q_n\right)dq_\mathrm{n} = \int_0^1 q_n^{b_n}\left(1-q_\mathrm{n}\right)^{1-b_n} \frac{\Gamma\left(\frac{\gamma_b}{\mathrm{N}}+\frac{N-1}{\mathrm{N}}\right)}{\Gamma\left(\frac{\gamma_b}{N}\right)\Gamma\left(\frac{N-1}{N}\right)} q_n^{\frac{\gamma_b}{N}-1}\left(1-q_n\right)^{\frac{N-1}{N}-1} dq_n \\
&= \frac{\Gamma\left(\frac{\gamma_b}{N}+\frac{N-1}{N}\right)}{\Gamma\left(\frac{\gamma_b}{N}\right)\Gamma\left(\frac{N-1}{N}\right)} \int_0^1 q_n^{\frac{\gamma_b}{N}+b_n-1}\left(1-q_n\right)^{\frac{N-1}{N}-b_n+1-1} dq_\mathrm{n} \\
&= \frac{\Gamma\left(\frac{\gamma_b}{N}+\frac{N-1}{N}\right)}{\Gamma\left(\frac{\gamma_b}{N}\right)\Gamma\left(\frac{N-1}{N}\right)} \frac{\Gamma\left(\frac{\gamma_b}{N}+b_n\right)\Gamma\left(\frac{N-1}{N}-b_n+1\right)}{\Gamma\left(\frac{\gamma_b}{N}+\frac{N-1}{N}+1\right)} \underbrace{\int_0^1 \textbf{Beta}\left(q_n; \frac{\gamma_b}{N}+b_n, \frac{N-1}{N}-b_n+1\right)dq_n}_{1} \\
&= \frac{1}{\frac{\gamma_b+(N-1)}{N}} \frac{\Gamma\left(\frac{\gamma_b}{N}+b_n\right)\Gamma\left(\frac{N-1}{\mathrm{N}}-b_\mathrm{n}+1\right)}{\Gamma\left(\frac{\gamma_b}{N}\right)\Gamma\left(\frac{N-1}{N}\right)}.
\end{aligned} \tag{S19}$$

As $b_n$ can only attain values of 0 or 1, we arrive at a renormalized Bernoulli probability distribution over $b_n$

$$\begin{cases} \mathbb{P}\left(b_n=0\right) = \frac{1}{1+\frac{\gamma_b}{N-1}} \\ \mathbb{P}\left(b_n=1\right) = \frac{1}{1+\frac{N-1}{\gamma_b}} \end{cases} \Longrightarrow b_n \sim \textbf{Bernoulli}\left(\frac{1}{1+\frac{N-1}{\gamma_b}}\right). \tag{S20}$$

##### S4.2. Summary of our model

For concreteness, below we summarize the entire set of equations used in our framework, including a complete list of priors and hyperpriors

$$D \sim \mathbf{InvGamma}, (\alpha_D, \beta_D) \tag{S21}$$

$$\mu_m^{\text{mol}} \sim \mathbf{Gamma}\left(\alpha_{\text{mol}}, \beta_{\text{mol}}\right) \tag{S22}$$
$$\mu_m^{\text{back}} \sim \mathbf{Gamma}\left(\alpha_{\text{back}}, \beta_{\text{back}}\right) \tag{S23}$$

$$b_n \sim \mathbf{Bernoulli}\left(\frac{1}{1+\frac{N-1}{\gamma_b}}\right) \tag{S24}$$

$$x_{n,1} \sim \mathbf{Normal}\left(x_0, \sigma_{x_0}^2\right) \tag{S25}$$
$$y_{n,1} \sim \mathbf{Normal}\left(y_0, \sigma_{y_0}^2\right) \tag{S26}$$
$$z_{n,1} \sim \mathbf{Normal}\left(z_0, \sigma_{z_0}^2\right) \tag{S27}$$

$$x_{n,k+1}|x_{n,k}, D \sim \mathbf{Normal}\left(x_{n,k}, 2D\Delta_k\right), \qquad k=1,\dots,K-1 \tag{S28}$$
$$y_{n,k+1}|y_{n,k}, D \sim \mathbf{Normal}\left(y_{n,k}, 2D\Delta_k\right), \qquad k=1,\dots,K-1 \tag{S29}$$
$$z_{n,k+1}|z_{n,k}, D \sim \mathbf{Normal}\left(z_{n,k}, 2D\Delta_k\right), \qquad k=1,\dots,K-1 \tag{S30}$$

$$\Delta_k|\{\mu_m^{\text{mol}}, \mu_m^{\text{back}}\}_m, \{b_n, x_{n,k}, y_{n,k}, z_{n,k}\}_n \sim \mathbf{Exp}\left(\sum_{m=1}^{M} \mu_{m,k}\right), \qquad k=1,\dots,K-1 \tag{S31}$$

$$s_k|\{\mu_m^{\text{mol}}, \mu_m^{\text{back}}\}_m, \{b_n, x_{n,k}, y_{n,k}, z_{n,k}\}_n \sim \mathbf{Cat}_{1,\dots,M}\left(\frac{\mu_{1,k}}{\sum_{m=1}^{M}\mu_{m,k}}, \dots, \frac{\mu_{M,k}}{\sum_{m=1}^{M}\mu_{m,k}}\right), \qquad k=1,\dots,K \tag{S32}$$

$$\mu_{m,k} = \left(\mu_m^{\text{back}} + \mu_m^{\text{mol}} \sum_n b_n \ \text{PSF}_m\left(x_{n,k}, y_{n,k}, z_{n,k}\right)\right). \tag{S33}$$

#### S5. Description of the computational scheme

The posterior over all unknowns that we wish to infer is $\mathbb{P}\left(D, \{\mu_m^{\text{mol}}, \mu_m^{\text{back}}\}_m, \{b_n, \overline{x}_n, \overline{y}_n, \overline{z}_n\}_n | \overline{\Delta}, \overline{s}\right)$, where molecular trajectories and intensities (measurements) are gathered in

$$\overline{x}_n = (x_{\text{n},1}, x_{n,2}, \dots, x_{n,K}) \tag{S34}$$
$$\overline{y}_n = (y_{\text{n},1}, y_{n,2}, \dots, y_{n,K}) \tag{S35}$$
$$\overline{z}_n = (z_{\text{n},1}, z_{n,2}, \dots, z_{n,K}) \tag{S36}$$
$$\overline{\Delta} = (\Delta_1, \Delta_2, \dots, \Delta_{K-1}) \tag{S37}$$
$$\overline{s} = (s_1, s_2, \dots, s_K). \tag{S38}$$

This posterior corresponds to the graphical model shown in Fig. 6. Due to the nonlinear dependency over molecular positions introduced in the PSF and the non-parametric prior on $b_n$, analytic evaluation or direct sampling of this posterior is impossible. For this reason, we develop a specialized Markov chain Monte Carlo (MCMC) scheme that can be used to generate pseudo-random samples from this posterior. [22–26] This scheme is explained in detail below.

The implementation of the proposed model as the source code and GUI, see Fig. S12, are available through the Supporting Materials.

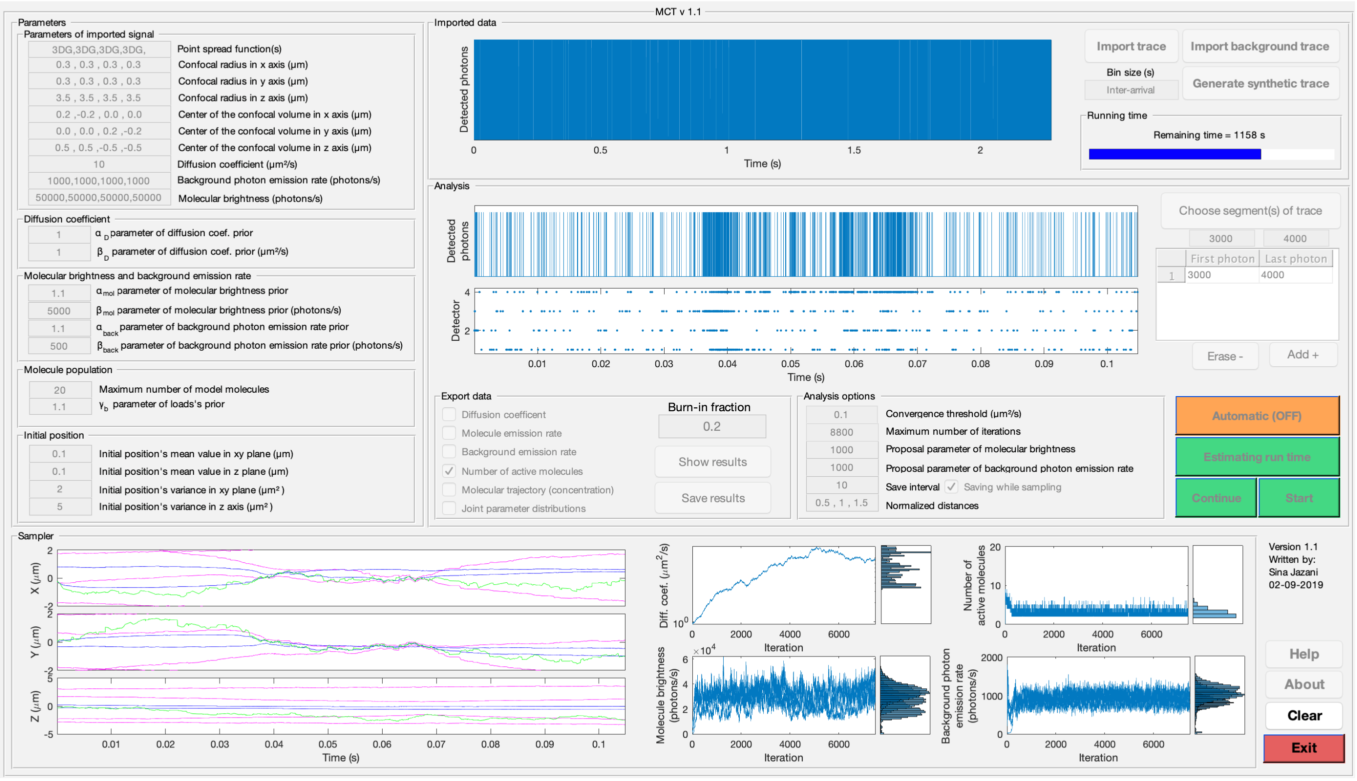


FIG. S12. **A working implementation of the framework described in this study is available through the Supporting Materials.** Along with this implementation, we provide a graphical user interface (GUI) that can be used to analyze intensity traces from confocal microscopy.

#### S5.1. Overview of the sampling updates

The MCMC we used exploits a Gibbs sampling scheme [22–24] by sampling each one of the random variables sequentially from their conditional probabilities on other random variables and the measurements $\overline{\Delta}$ and $\overline{s}$. Conceptually, the steps involved in the generation of each posterior sample $\left(D, \{\mu_m^{\text{mol}}, \mu_m^{\text{back}}\}_m, \{b_n, \overline{x}_n, \overline{y}_n, \overline{z}_n\}_n\right)$ are:

(1) For each n of the *active* molecules

  (i) Update trajectory $\overline{x}_n$ of active molecule $n$

  (ii) Update trajectory $\overline{y}_n$ of active molecule $n$

  (iii) Update trajectory $\overline{z}_n$ of active molecule $n$

(2) Update jointly the trajectories $\overline{x}_n, \overline{y}_n, \overline{z}_n$ for all $n$ of the *inactive* molecules

(3) Update the diffusion coefficient $D$

(4) Update jointly the loads $b_n$ for all model molecules

(5) Update jointly the molecular brightness and background photon emission rates $\mu_m^{\text{mol}}$ and $\mu_m^{\text{back}}$, respectively

These steps are described in detail below.

#### S5.2. Sampling of active molecule trajectories

We sample the trajectory of active molecules $\overline{x}_n$ from the corresponding conditional probability distribution $\mathbb{P}\left(\overline{x}_n | D, \{\mu_m^{\text{mol}}, \mu_m^{\text{back}}\}_m, \{b_{n'}, \overline{y}_{n'}, \overline{z}_{n'}\}_{n'}, \{\overline{x}_{n'}\}_{n' \neq n}, \overline{\Delta}, \overline{s}\right)$. We, directly sample the trajectories using Hamiltonian Monte Carlo (HMC) [23, 27, 28] which we expand below.

As far as we know, this strategy has rarely been used in the Natural Sciences and yet is critical in avoiding approximations in sampling the molecular trajectories. In our previous work [4, 18, 19] we used Kalman filters for

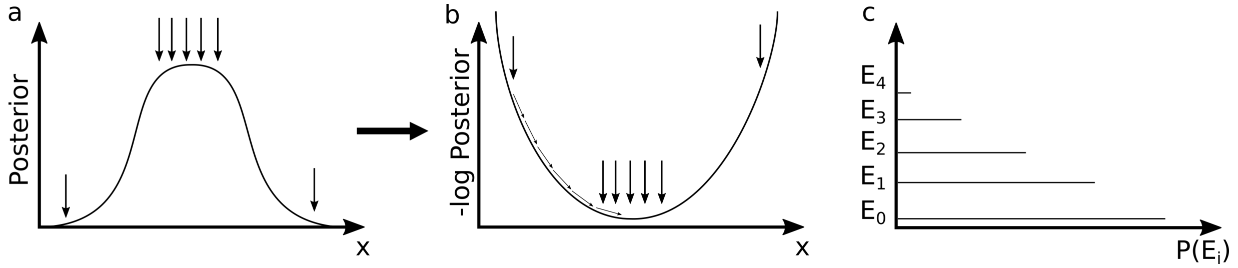


FIG. S13. **A cartoon representation of HMC. (a)** Samples from the posterior for molecular location are often more likely drawn near the posterior's mode. As a result, in **(b)**, samples are drawn more often near the minimum or minima of the negative logarithm of the posterior. **(c)** In HMC, we ascribe an interpretation of the negative logarithm of the posterior. We think of it as a potential and think of HMC as a means of locating potential minima. Each equiprobable region of the posterior is thought of as an isoenergetic surface of our potential.

this task due to its computational efficiency, and to do this, we needed to approximate the likelihood which results in new point statistics that appear as a transformation of the data. To do this approximation, we imposed some assumptions and as the result, the posterior sampled was an approximate posterior. Now, by applying HMC, we can directly target any posterior, without concern as to the complex dependency of the likelihood on the parameters we wish to infer.

In HMC, we have three main steps:

(1) Posterior transformation to a Hamiltonian

(2) Perform Strang-splitting [29, 30] to solve the Hamiltonian equations and propose locations of molecules

(3) Perform a Metropolis-Hastings to accept or reject the proposed sample.

The benefits of posterior transformation to a Hamiltonian is that, we can estimate the random variables which in this case are the positions of the molecule by solving the Hamiltonian equations.

##### S5.2.1. Posterior transformation to a Hamiltonian

The idea underlying HMC is summarized in Fig. S13. Without approximation, we re-write the posterior probability distribution as $\mathbb{P}\left(\overline{x}_n|\overline{\Delta},\overline{s},\dots\right)\propto\exp\left(-U\left(\overline{x}_n\right)\right)$ with the potential energy defined as $U\left(\overline{x}_n\right)=-\log\mathbb{P}\left(\overline{x}_n|\overline{\Delta},\overline{s},\dots\right)$.
The target probability distribution to sample from is

$$\begin{aligned}\mathbb{P}\big(\overline{x}_n|D,\{\mu_m^{\text{mol}},\mu_m^{\text{back}}\}_m,&\{b_{n'},\overline{y}_{n'},\overline{z}_{n'}\}_{n'},\{\overline{x}_{n'}\}_{n'\neq n},\overline{\Delta},\overline{s}\big)\propto\\ &\mathbb{P}\left(\overline{s}|\overline{x}_n,D,\{\mu_m^{\text{mol}},\mu_m^{\text{back}}\}_m,\{b_{n'},\overline{y}_{n'},\overline{z}_{n'}\}_{n'},\{\overline{x}_{n'}\}_{n'\neq n}\right)\\ &\times\mathbb{P}\left(\overline{\Delta}|\overline{x}_n,D,\{\mu_m^{\text{mol}},\mu_m^{\text{back}}\}_m,\{b_{n'},\overline{y}_{n'},\overline{z}_{n'}\}_{\text{n}'},\{\overline{x}_{n'}\}_{n'\neq n}\right)\\ &\times\mathbb{P}\left(\overline{x}_n|D,\overline{\Delta}\right)\end{aligned}\tag{S39}$$

where the positions of the active molecule $\overline{x}_n$ are the variables and both $\overline{\Delta}$ and $\overline{s}$ are observations. In HMC, the logarithm of the above conditional coincides with what is termed the "HMC potential"

$$\begin{aligned}U(\overline{x}_n)&=-\log\mathbb{P}(\overline{x}_n|\overline{\Delta},\overline{s},\dots)\\ &=-\log\mathbb{P}(\overline{s}|\overline{x}_n,\dots)-\log\mathbb{P}(\overline{\Delta}|\overline{x}_n,\dots)-\log\mathbb{P}(\overline{x}_n|\overline{\Delta},D)+\text{Constant}\\ &=V(\overline{x}_n)+L(\overline{x}_n)+\text{Constant}.\end{aligned}\tag{S40}$$

For computational reasons, we have split the potential into two where $V\left(\overline{x}_n\right)=-\log\left(\mathbb{P}\left(\overline{\Delta}|\overline{x}_n,\dots\right)\mathbb{P}\left(\overline{s}|\overline{x}_n,\dots\right)\right)$

contains the likelihood portion of the posterior and $L\left(\overline{x}_n\right) = -\log \mathbb{P}\left(\overline{x}_n | D, \overline{\varDelta},\right)$ contains the prior.

$$V(\overline{x}_n) = -\log\left(\left[\prod_{k=1}^{K-1} \mathbb{P}\left(\varDelta_k | x_{n,k}, \dots\right) \mathbb{P}\left(s_k | x_{n,k}, \dots\right)\right] \mathbb{P}\left(s_K | x_{n,K}, \dots\right)\right) \tag{S41}$$

$$= -\log\left(\left[\prod_{k=1}^{K-1} \mathrm{EXP}\left(\varDelta_k; \sum_{m=1}^{M} \mu_{m,k}\right) \frac{\mu_{s_k,k}}{\sum_{m=1}^{M} \mu_{m,k}}\right] \frac{\mu_{s_K,K}}{\sum_{m=1}^{M} \mu_{m,K}}\right) \tag{S42}$$

$$= -\log\left(\left[\prod_{k=1}^{K-1} \mu_{s_k,k} \exp\left(-\varDelta_k \sum_{m=1}^{M} \mu_{m,k}\right)\right] \frac{\mu_{s_K,K}}{\sum_{m=1}^{M} \mu_{m,K}}\right) \tag{S43}$$

$$= \left[\sum_{k=1}^{K-1} -\log\left(\mu_{s_k,k}\right) + \varDelta_k \sum_{m=1}^{M} \mu_{m,k}\right] - \log\left(\mu_{s_K,K}\right) + \log\left(\sum_{m=1}^{M} \mu_{m,K}\right). \tag{S44}$$

Also, as the result of expanding the partial potential $L\left(\overline{x}_n\right) = -\log \mathbb{P}\left(\overline{x}_n | \overline{\varDelta}, D\right)$ in Eq. (S39) we have

$$\mathbb{P}(\overline{x}_n | D, \overline{\varDelta}) = \mathbb{P}(x_{n,1}) \mathbb{P}(x_{n,2} | x_{n,1}, \varDelta_1, D) \cdots \mathbb{P}(x_{n,K} | x_{n,K-1}, \varDelta_{K-1}, D). \tag{S45}$$

Since $\mathbb{P}(x_{\mathrm{n},1})$ above is given by Eq. (S14) and the subsequent conditional probabilities are dictated by the motion model, Eq. (S28), we arrive at

$$\mathbb{P}(\overline{x}_n | \overline{\varDelta}, D) = \mathbf{Normal}\left(x_{n,1}; x_0, \sigma_{x_0}^2\right) \mathbf{Normal}\left(x_{n,2}; x_{n,1}, 2D\varDelta_1\right) \cdots \mathbf{Normal}\left(x_{n,K}; x_{n,K-1}, 2D\varDelta_{K-1}\right) \tag{S46}$$

where $x_0$, $\sigma_{x_0}^2$ are the mean value and the variance of the initial position's prior in Eqs. (S25), (S26), (S27). The negative logarithm of the above yields

$$\begin{aligned}
L(\overline{x}_n) &= -\log\ \mathbb{P}(\overline{x}_n) \\
&= -\log\ \mathbf{Normal}\left(x_{n,1}; \mu_{x_0}, \sigma_{x_0}^2\right) - \log\ \mathbf{Normal}\left(x_{n,2}; x_{n,1}, 2D\varDelta_1\right) - \log\ \mathbf{Normal}\left(x_{n,3}; x_{n,2}, 2D\varDelta_2\right) \\
&\quad \vdots \\
&\quad -\log\ \mathbf{Normal}\left(x_{n,K-1}; x_{n,K-2}, 2D\varDelta_{K-2}\right) - \log\ \mathbf{Normal}\left(x_{n,K}; x_{n,K-1}, 2D\varDelta_{K-1}\right) \\
&= +\frac{1}{2\sigma_x^2}(x_{n,1} - \mu)^2 \\
&\quad + \frac{1}{4D\varDelta_1}(x_{n,2} - x_{n,1})^2 + \frac{1}{4D\varDelta_2}(x_{n,3} - x_{n,2})^2 \\
&\quad \vdots \\
&\quad + \frac{1}{4D\varDelta_{K-2}}(x_{n,K-1} - x_{n,K-2})^2 + \frac{1}{4D\varDelta_{K-1}}(x_{n,K} - x_{n,K-1})^2 + \text{Constant.}
\end{aligned} \tag{S47}$$

Now that we have the potential $U(\overline{x}_n) = V(\overline{x}_n) + L(\overline{x}_n)$, we define a kinetic energy required for the HMC sampler. To do this, we introduce a new set of auxiliary random variables, $\overline{p}_n$, as well as a mass matrix $\mathbf{M}$. As usual, the kinetic energy is given by

$$T(\overline{p}_n) = \frac{\overline{p}_n^T \mathbf{M}^{-1} \overline{p}_n}{2}. \tag{S48}$$

The mass matrix $\mathbf{M}$ has to be positive definite (i.e. has exclusively positive real eigenvalues). Since, any choice that satisfies this requirement works, we choose the simplest choice which is the diagonal matrix [27]

$$\mathbf{M} = \begin{bmatrix} m_1 & 0 & \dots & 0 \\ 0 & m_2 & \dots & 0 \\ \vdots & \vdots & \ddots & \vdots \\ 0 & 0 & \dots & m_K \end{bmatrix}. \tag{S49}$$

The resulting full Hamiltonian (ignoring the constant term), the key quantity of HMC, is separable and of the form

$$\begin{aligned} H(\overline{x}_n, \overline{p}_n) &= T(\overline{p}_n) + U(\overline{x}_n) \\ &= T(\overline{p}_n) + V(\overline{x}_n) + L(\overline{x}_n). \end{aligned} \tag{S50}$$

##### S5.2.2. Perform the Strang-splitting algorithm to solve Hamilton's equations

In order to find $\overline{x}_n$ and $\overline{p}_n$, we need solve the Hamiltonian's equations [27, 28]

$$\frac{d\overline{x}_n}{dt} = +H_{\overline{p}_n}(\overline{x}_n, \overline{p}_n) \tag{S51}$$

$$\frac{d\overline{p}_n}{dt} = -H_{\overline{x}_n}(\overline{x}_n, \overline{p}_n). \tag{S52}$$

where $H_{\overline{p}_n}$ and $H_{\overline{x}_n}$ are gradients of the Hamiltonian with respect to the subscripted quantity. If we could solve Eqs. (S51), (S52) without approximation, we could directly sample the whole trajectory of the molecule $n$, $\overline{x}_n$. However, in our case, there is no analytic solution and we need to use some numerical method to solve them.

To solve the Hamilton's equations Eqs. (S51), (S52), we use Strang-splitting [29, 30] which is a symplectic integrator preserving the energy of the mechanical system (and thus the target probability distribution from which we want to sample) and preserving the phase volume. Since, Strang-splitting, is a numerical method to solve differential equations, we might have some error in the final answer. So, to correct such error, at the end we evaluate the answer by a Metropolis-Hastings algorithm to avoid any error caused by the Strang-splitting.

To use the Strang-splitting, we split the Hamiltonian in Eq. (S53) into two Hamiltonians:

$$H(\overline{x}_n, \overline{p}_n) = H^1(\overline{x}_n, \overline{p}_n) + H^2(\overline{x}_n, \overline{p}_n) \tag{S53}$$

where

$$H^1(\overline{x}_n, \overline{p}_n) = V(\overline{x}_n) \tag{S54}$$

$$H^2(\overline{x}_n, \overline{p}_n) = T(p) + L(\overline{x}_n). \tag{S55}$$

To integrate the dynamics, in Strang-splitting, instead of considering the full step, we consider many small steps with a step size of $h$ and integrate each of the partial Hamiltonians $H^1(\overline{x}_n, \overline{p}_n)$ and $H^2(\overline{x}_n, \overline{p}_n)$ successively. As the result, Strang-splitting is based on single steps with the following fractional steps of

(1) Advance half-step using $H^1(\overline{x}_n, \overline{p}_n)$
(2) Advance whole-step using $H^2(\overline{x}_n, \overline{p}_n)$
(3) Advance half-step using $H^1(\overline{x}_n, \overline{p}_n)$.

Taken together, these three steps are equal to a full step for each Hamiltonian. Below, we describe each of these steps in detail.

**(1) Advance half-step using $H^1(\overline{x}_n, \overline{p}_n)$**

In this step we have a half-update using the $H^1(\overline{x}_n, \overline{p}_n) = V(\overline{x}_n)$ and integrate the dynamics as follows

$$\frac{d\overline{x}_n}{dt} = +H^1_{\overline{p}_n}(\overline{x}_n, \overline{p}_n) \tag{S56}$$

$$\frac{d\overline{p}_n}{dt} = -H^1_{\overline{x}_n}(\overline{x}_n, \overline{p}_n) \tag{S57}$$

which simplifies to

$$\frac{d\overline{x}_n}{dt} = 0 \tag{S58}$$

$$\frac{d\overline{p}_n}{dt} = -V_{\overline{x}_n}(\overline{x}_n). \tag{S59}$$

To solve Eqs. (S58), (S59), we use Stormer-Verlet, [31] which is second order and symplectic

$$\frac{\overline{p}_n^{\text{mid}} - \overline{p}_n^{old}}{h/4} = -H^1_{\overline{x}_n}(\overline{p}_n^{\text{mid}}, \overline{x}_n^{\text{old}}) \tag{S60}$$

$$\frac{\overline{x}_n^{\text{new}} - \overline{x}_n^{old}}{h/4} = -H^1_{\overline{p}_n}(\overline{p}_n^{\text{mid}}, \overline{x}_n^{\text{old}}) + H^1_{\overline{p}_n}(\overline{p}_n^{mid}, \overline{x}_n^{new}) \tag{S61}$$

$$\frac{\overline{p}_n^{\text{new}} - \overline{p}_n^{mid}}{h/4} = -H^1_{\overline{x}_n}(\overline{p}_n^{\text{mid}}, \overline{x}_n^{\text{new}}) \tag{S62}$$

which immediately simplifies to

$$\frac{\overline{p}_n^{\text{mid}} - \overline{p}_n^{\text{old}}}{h/4} = -V_{\overline{x}_n}(\overline{x}_n^{\text{old}}) \tag{S63}$$

$$\frac{\overline{x}_n^{\text{new}} - \overline{x}_n^{\text{old}}}{h/4} = 0 \tag{S64}$$

$$\frac{\overline{p}_n^{\text{new}} - \overline{p}_n^{\text{mid}}}{h/4} = -V_{\overline{x}_n}(\overline{x}_n^{\text{new}}) \tag{S65}$$

such that

$$\overline{x}_n^{\text{new}} = \overline{x}_n^{\text{old}} \tag{S66}$$

$$\overline{p}_{\text{n}}^{\text{new}} = \overline{p}_{\text{n}}^{\text{old}} - \frac{h}{2} V_{\overline{x}_{\text{n}}}\left(\overline{x}_{\text{n}}^{\text{old}}\right). \tag{S67}$$

The gradient of the partial potential $V(\overline{x}_n)$ can be written as

$$V_{\overline{x}_n}(\overline{x}_n) = \frac{\partial V_{\overline{x}_n}(\overline{x}_n)}{\partial \overline{x}_n} = \frac{\partial}{\partial \overline{x}_n}\left(\left[\sum_{k=1}^{K-1} -\log\left(\mu_{s_k,k}\right) + \Delta_k \sum_{m=1}^{M} \mu_{m,k}\right] - \log\left(\mu_{s_K,K}\right) + \log\left(\sum_{m=1}^{M} \mu_{m,K}\right)\right) \tag{S68}$$

$$= \begin{bmatrix} -\frac{\frac{\partial \mu_{s_1,1}}{\partial x_{n,1}}}{\mu_{s_1,1}} + \Delta_1 \sum_{m=1}^{M} \frac{\partial \mu_{m,1}}{\partial x_{n,1}} \\ -\frac{\frac{\partial \mu_{s_2,2}}{\partial x_{n,2}}}{\mu_{s_2,2}} + \Delta_2 \sum_{m=1}^{M} \frac{\partial \mu_{m,2}}{\partial x_{n,2}} \\ -\frac{\frac{\partial \mu_{s_3,3}}{\partial x_{n,3}}}{\mu_{s_3,3}} + \Delta_3 \sum_{m=1}^{M} \frac{\partial \mu_{m,3}}{\partial x_{n,3}} \\ \vdots \\ -\frac{\frac{\partial \mu_{s_{K-1},K-1}}{\partial x_{n,K-1}}}{\mu_{s_{K-1},K-1}} + \Delta_{K-1} \sum_{m=1}^{M} \frac{\partial \mu_{m,K-1}}{\partial x_{n,K-1}} \\ -\frac{\frac{\partial \mu_{s_K,K}}{\partial x_{n,K}}}{\mu_{s_K,K}} + \frac{\sum_{m=1}^{M} \frac{\partial \mu_{m,K}}{\partial x_{n,K}}}{\sum_{m=1}^{M} \mu_{m,K}} \end{bmatrix}. \tag{S69}$$

**(2) Advance whole-step using $H^2(\overline{x}_n, \overline{p}_n)$**

In this step we have a whole-update using the $H^2(\overline{x}_n, \overline{p}_n) = T(\overline{p}_n) + L(\overline{x}_n)$ and integrate the dynamics as follows

$$\frac{d\overline{x}_n}{dt} = +H^2_{\overline{p}_n}(\overline{x}_n, \overline{p}_n) \tag{S70}$$

$$\frac{d\overline{p}_n}{dt} = -H^2_{\overline{x}_n}(\overline{x}_n, \overline{p}_n) \tag{S71}$$

which simplifies to

$$\frac{d\overline{x}_n}{dt} = M^{-1}\overline{p}_n \tag{S72}$$

$$\frac{d\overline{p}_n}{dt} = -L_{\overline{x}_n}(\overline{x}_n). \tag{S73}$$

The gradient of the partial potential $L(\overline{x}_n)$ is

$$
L_{\overline{x}_n}(\overline{x}_n) = \begin{bmatrix} L_{x_{n,1}}(\overline{x}_n) \\ L_{x_{n,2}}(\overline{x}_n) \\ L_{x_{n,3}}(\overline{x}_n) \\ \vdots \\ L_{x_{n,K-1}}(\overline{x}_n) \\ L_{x_{n,K}}(\overline{x}_n) \end{bmatrix} = \begin{bmatrix} \frac{(x_{n,1}-\mu_{x_0})}{\sigma_{x_0}^2} + \frac{(x_{n,1}-x_{n,2})}{2D\,(t_2-t_1)} \\ \frac{(x_{n,2}-x_{n,1})}{2D\,(t_2-t_1)} + \frac{(x_{n,2}-x_{n,3})}{2D\,(t_3-t_2)} \\ \frac{(x_{n,3}-x_{n,2})}{2D\,(t_3-t_2)} + \frac{(x_{n,3}-x_{n,4})}{2D\,(t_4-t_3)} \\ \vdots \\ \frac{(x_{n,K-1}-x_{n,K-2})}{2D\,(t_{K-1}-t_{K-2})} + \frac{(x_{n,K-1}-x_{n,K})}{2D\,(t_K-t_{K-1})} \\ \frac{(x_{n,K}-x_{n,\mathrm{K}-1})}{2D\,(t_K-t_{K-1})} \end{bmatrix} = \begin{bmatrix} \frac{(x_{n,1}-\mu_{x_0})}{\sigma_{x_0}^2} + \frac{(x_{n,1}-x_{n,2})}{2D\varDelta_1} \\ \frac{(x_{n,2}-x_{n,1})}{2D\varDelta_1} + \frac{(x_{n,2}-x_{n,3})}{2D\varDelta_2} \\ \frac{(x_{n,3}-x_{n,2})}{2D\varDelta_2} + \frac{(x_{n,3}-x_{n,4})}{2D\varDelta_3} \\ \vdots \\ \frac{(x_{n,K-1}-x_{n,K-2})}{2D\varDelta_{K-2}} + \frac{(x_{n,K-1}-xn,K)}{2D\varDelta_{K-1}} \\ \frac{(x_{n,K}-xn,K-1)}{2D\varDelta_{K-1}} \end{bmatrix}
$$

$$
= -\frac{1}{2D}\begin{bmatrix} -(\frac{2D}{\sigma_{x_0}^2}+\frac{1}{\varDelta_1}) & \frac{1}{\varDelta_1} & 0 & \dots & 0 & 0 \\ \frac{1}{\varDelta_1} & -(\frac{1}{\varDelta_1}+\frac{1}{\varDelta_2}) & \frac{1}{\varDelta_2} & \dots & 0 & 0 \\ 0 & \frac{1}{\varDelta_2} & -(\frac{1}{\varDelta_2}+\frac{1}{\varDelta_3}) & \dots & 0 & 0 \\ \vdots & \vdots & \vdots & \ddots & \vdots & \vdots \\ 0 & 0 & 0 & \dots & -(\frac{1}{\varDelta_{K-2}}+\frac{1}{\varDelta_{K-1}}) & \frac{1}{\varDelta_{K-1}} \\ 0 & 0 & 0 & \dots & \frac{1}{\varDelta_{K-1}} & -\frac{1}{\varDelta_{K-1}} \end{bmatrix}\begin{bmatrix} x_{n,1} \\ x_{n,2} \\ x_{n,3} \\ \vdots \\ x_{n,K-1} \\ x_{n,K} \end{bmatrix} - \frac{1}{\sigma_{x_0}^2}\begin{bmatrix} x_0 \\ 0 \\ 0 \\ \vdots \\ 0 \\ 0 \end{bmatrix}
$$

$$
= -\frac{1}{2D}\mathbf{A}\overline{x}_n - \frac{1}{\sigma_{x_0}^2}\overline{\nu}. \tag{S74}
$$

Now, inserting the gradient, shown in Eq. (S74), into Eqs. (S72) and (S73), we have

$$
\begin{aligned} \frac{d\overline{x}_n}{dt} &= \mathbf{M}^{-1}\overline{p}_n \\ \frac{d\overline{p}_n}{dt} &= \frac{1}{2D}\mathbf{A}\overline{x}_n + \frac{1}{\sigma_{x_0}^2}\overline{\nu}. \end{aligned} \tag{S75}
$$

To be able to solve the above equations efficiently, we apply the implicit midpoint method, [31] which is second order and symplectic. By applying the implicit midpoint, we recover

$$
\begin{aligned} \frac{\overline{x}_n^{\mathrm{new}} - \overline{x}_n^{\mathrm{old}}}{h} &= \mathbf{M}^{-1}\frac{\overline{p}_n^{\mathrm{old}} + \overline{p}_n^{\mathrm{new}}}{2} \\ \frac{\overline{p}_n^{\mathrm{new}} - \overline{p}_n^{\mathrm{old}}}{h} &= \frac{1}{2D}\mathbf{A}\frac{\overline{x}_n^{\mathrm{old}} + \overline{x}_n^{\mathrm{new}}}{2} + \frac{1}{\sigma_{x_0}^2}\overline{\nu}. \end{aligned} \tag{S76}
$$

Next, we first solve $\overline{p}_n^{\mathrm{new}}$ from the first equation and plug it into the second equation

$$
\left(\frac{2\mathbf{M}}{h} - \frac{h}{4D}\mathbf{A}\right)\overline{x}_n^{\mathrm{new}} = \left(\frac{2\mathbf{M}}{h} + \frac{h}{4D}\mathbf{A}\right)\overline{x}_n^{\mathrm{old}} + \frac{h}{\sigma_{x_0}^2}\overline{\nu} + 2\overline{p}_n^{\mathrm{old}} \tag{S77}
$$

$$
\overline{p}_n^{\mathrm{new}} = \frac{2\mathbf{M}}{h}\left(\overline{x}_n^{\mathrm{new}} - \overline{x}_n^{\mathrm{old}}\right) - \overline{p}_n^{\mathrm{old}}. \tag{S78}
$$

In order to simplify Eqns. (S77), (S78), we now introduce these metrices

$$
\mathbf{G} = \frac{2}{h}\mathbf{M} \tag{S79}
$$

$$
\mathbf{F}_1 = \mathbf{G} - \frac{h}{4D}\mathbf{A} \tag{S80}
$$

$$
\mathbf{F}_2 = \mathbf{G} + \frac{h}{4D}\mathbf{A} \tag{S81}
$$

and recast Eqns. (S77), (S78) in the form

$$\mathbf{F}_1\overline{x}_n^{\text{new}} = \mathbf{F}_2\overline{x}_n^{\text{old}} + \frac{h}{\sigma_x^2}\overline{\nu} + 2p_n^{\text{old}} \tag{S82}$$

$$p_n^{\text{new}} = \mathbf{G}\left(\overline{x}_n^{\text{new}} - \overline{x}_n^{\text{old}}\right) - p_n^{\text{old}}. \tag{S83}$$

Next, our goal is to solve Eqs. (S82), (S83) to find the $\overline{x}_n^{\text{new}}$ and the $p_n^{\text{new}}$. To solve these equations, we use tri-diagonal solver also known as Thomas algorithm [32] (this is due to the tri-diagonality of $\mathrm{L}(\overline{x}_n)$ which leads to tri-diagonal $\mathbf{F}_1$ and $\mathbf{F}_2$). Re-writing Eq. (S82) we have

$$\mathbf{F}_1\overline{x}_n^{\text{new}} = d \tag{S84}$$

where $d = \mathbf{F}_2\overline{x}_n^{\text{old}} + \frac{h}{\sigma_{x_0}^2}\overline{\nu} + 2\overline{p}_n^{\text{old}}$. With all matrices made explicit, we have

$$\begin{bmatrix} \phi_1 & \psi_1 & 0 & 0 & \dots & 0 & 0 & 0 \\ \psi_1 & \phi_2 & \psi_2 & 0 & \dots & 0 & 0 & 0 \\ 0 & \psi_2 & \phi_3 & \psi_3 & \dots & 0 & 0 & 0 \\ \vdots & \vdots & \vdots & \vdots & \ddots & \vdots & \vdots & \vdots \\ 0 & 0 & 0 & 0 & \dots & \psi_{K-2} & \phi_{K-1} & \psi_{K-1} \\ 0 & 0 & 0 & 0 & \dots & 0 & \psi_{K-1} & \phi_K \end{bmatrix} \begin{bmatrix} x_{n,1}^{\text{new}} \\ x_{n,2}^{\text{new}} \\ x_{n,3}^{\text{new}} \\ \vdots \\ x_{n,K-1}^{\text{new}} \\ x_{n,K}^{\text{new}} \end{bmatrix} = \begin{bmatrix} d_1 \\ d_2 \\ d_3 \\ \vdots \\ d_{K-1} \\ d_K \end{bmatrix} \tag{S85}$$

where, $\overline{\psi}$ and $\overline{\phi}$ are

$$\psi_k = -\frac{h}{4D\varDelta_k} \quad , \quad k = 1, \dots, K$$

$$\phi_k = \begin{cases} \frac{2m_k}{h} + \frac{h}{4D}\left(\frac{2D}{\sigma_{x_0}^2} + \frac{1}{\varDelta_k}\right) & , \quad k = 1 \\ \frac{2m_k}{h} + \frac{h}{4D}\left(\frac{1}{\varDelta_k} + \frac{1}{\varDelta_{k-1}}\right) & , \quad k = 2, \dots, K-1 \\ \frac{2m_k}{h} + \frac{h}{4D}\frac{1}{\varDelta_{k-1}} & , \quad k = K. \end{cases} \tag{S86}$$

To implement the Thomas algorithm to solve Eq. (S85), we must first "march forward" as follows

$$C'_{\mathrm{k}} = \begin{cases} \frac{\psi_k}{\phi_k} & , \quad k = 1 \\ \frac{\psi_k}{\phi_k - \psi_{k-1}C'_{k-1}} & , \quad k = 2, \dots, K-1 \end{cases} \tag{S87}$$

$$d'_k = \begin{cases} \frac{d_k}{\phi_k} & , \quad k = 1 \\ \frac{d_k - \psi_{k-1}d'_{k-1}}{\phi_k - \psi_{k-1}C'_{k-1}} & , \quad k = 2, \dots, K \end{cases} \tag{S88}$$

and by marching backward we have

$$\begin{aligned} x_{n,K}^{\text{new}} &= d'_K \\ x_{n,k}^{\text{new}} &= d'_k - C'_k x_{n,k+1}^{\text{new}} \quad , \quad k = K-1, \dots, 1. \end{aligned} \tag{S89}$$

##### S5.2.3. Perform a Metropolis-Hastings test to accept or reject the proposed sample

Solutions of the Hamiltonian equations are deterministic. However, due to the error caused using of Strang-splitting to solve these equations, we now need to evaluate the proposed trajectory by comparing posteriors over the positions determined with the old posteriors in the Metropolis-algorithm and accept or reject positions $\overline{x}_n^{\text{new}}$. Since, we already, calculated the logarithm of the posterior, we compare the logarithm of the posterior ratio which is equal to the difference of Hamiltonians defined by Eq. (S50).

#### S5.3. Sampling of inactive molecule trajectories

After updating the trajectories of the active molecules, we update the trajectories of the inactive ones. For this, we sample from the corresponding conditionals $\mathbb{P}(\{\overline{x}_n, \overline{y}_n, \overline{z}_n\}_{n:b_n=0}|D, \{\mu^{\text{mol}}_{\underline{m}}, \mu^{\text{back}}_m\}_m, \{b_n\}_n, \overline{\Delta}, \overline{s})$. Since the locations of inactive molecules are not associated with the observations in $\overline{\Delta}$ and $\overline{s}$, these conditionals simplify to $\mathbb{P}(\{\overline{x}_n, \overline{y}_n, \overline{z}_n\}_{\text{n}:b_n=0}|D, \{b_n\}_n, \overline{\Delta})$ which can be readily simulated jointly in the same manner as standard 3D Brownian motion.

So, the conditional probability distribution $\mathbb{P}(\{\overline{x}_n, \overline{y}_n, \overline{z}_n\}_{n:b_n=0}|D, \{b_n\}_n, \overline{\Delta})$ can be written as

$$\begin{aligned}\mathbb{P}(\{\overline{x}_n, \overline{y}_n, \overline{z}_n\}_{n:b_n=0}|D, \{b_n\}_n, \overline{\Delta}) &= \mathbb{P}\left(\overline{x}_{n,b_n=0}|D, \{b_n\}_n, \overline{\Delta}\right) \mathbb{P}\left(\overline{y}_{n,b_n=0}|D, \{b_n\}_n, \overline{\Delta}\right) \mathbb{P}\left(\overline{z}_{n,b_n=0}|D, \{b_n\}_n, \overline{\Delta}\right) \\ &= \mathbb{P}\left(x_{n,1}\right) \prod_{k=1}^{K-1} \mathbb{P}\left(x_{\text{n,k+1}}|x_{n,k}, D, \Delta_k\right) \\ &\times \mathbb{P}\left(y_{n,1}\right) \prod_{k=1}^{K-1} \mathbb{P}\left(y_{\text{n,k+1}}|y_{n,k}, D, \Delta_k\right) \\ &\times \mathbb{P}\left(z_{n,1}\right) \prod_{k=1}^{K-1} \mathbb{P}\left(z_{\text{n,k+1}}|z_{n,k}, D, \Delta_k\right).\end{aligned} \tag{S90}$$

Since coordinates of $(x, y, z)$ are independent from each others, we can sample them separately. For the first positions of inactive molecules, we sample them form the prior.

$$\begin{aligned}x_{n,1,b_n=0} &\sim \textbf{Normal}\left(x_0, \sigma^2_{x_0}\right) \\ y_{n,1,b_n=0} &\sim \textbf{Normal}\left(y_0, \sigma^2_{y_0}\right) \\ z_{n,1,b_n=0} &\sim \textbf{Normal}\left(z_0, \sigma^2_{z_0}\right)\end{aligned} \tag{S91}$$

and for the rest of the trajectory, we march forward and sample them form

$$\begin{aligned}x_{n,k+1,b_n=0} &\sim \textbf{Normal}\left(x_{n,k,b_n=0}, D, \Delta_k\right) \\ y_{n,k+1,b_n=0} &\sim \textbf{Normal}\left(y_{n,k,b_n=0}, D, \Delta_k\right) \quad , \quad k = 1, \ldots, K-1 \\ z_{n,k+1,b_n=0} &\sim \textbf{Normal}\left(z_{n,k,b_n=0}, D, \Delta_k\right).\end{aligned} \tag{S92}$$

#### S5.4. Sampling the diffusion coefficient

By having the updated locations of molecules, we sample the diffusion coefficient $D$ from the corresponding conditional probability distribution of $\mathbb{P}\left(D|\{\mu^{\text{mol}}_m, \mu^{\text{back}}_m\}_m, \{b_n, \overline{x}_n, \overline{y}_n, \overline{z}_n\}_n, \overline{\Delta}, \overline{s}\right)$, which due to independence of the diffusion coefficient from the emission rates and the labels on the confocal volume, simplifies to $\mathbb{P}\left(D|\{\overline{x}_n, \overline{y}_n, \overline{z}_n\}_n, \overline{\Delta}\right)$; such variable dependencies are also shown graphically in Fig. 6. Now, using the prior, Eq. (S12), and motion model, Eqs. (S28), in 1D, we arrive at the marginal posterior

$$\begin{aligned}\mathbb{P}\left(D|\{\overline{x}_n\}_n, \overline{\Delta}\right) &\propto \prod_{n=1}^{N} \prod_{k=1}^{K-1} \textbf{Normal}\left(x_{n,k+1}; x_{n,k}, 2D\Delta_k\right) \textbf{InvGamma}\left(D; \alpha_D, \beta_D\right) \\ &= \frac{\beta_D^{\alpha_D}}{\Gamma\left(\alpha_D\right)\left(4\pi\right)^{\frac{N(k-1)}{2}}} D^{-(\alpha_D + \frac{N(k-1)}{2})-1} \exp\left(-\frac{\beta_D + \frac{1}{4}\sum_{n=1}^{N}\sum_{k=1}^{K-1}\frac{\left(x_{n,k+1}-x_{n,k}\right)^2}{\Delta_\text{k}}}{D}\right) \\ &\propto \textbf{InvGamma}\left(D; \alpha'_D, \beta'_D\right)\end{aligned} \tag{S93}$$

where $\alpha'_D$ and $\beta'_D$ are given by

$$\alpha'_D = \alpha_D + \frac{N(K-1)}{2}, \quad \beta'_D = \beta_D + \frac{1}{4}\sum_{n=1}^{N}\sum_{k=1}^{K-1}\frac{\left(x_{n,k+1}-x_{n,k}\right)^2}{\Delta_k}. \tag{S94}$$

In 3D, Eq. (S93) can be re-written by using Eqs. (S28), (S29) and (S29) and the parameters of $\alpha'_D$ and $\beta'_D$ have new form of

$$\alpha'_D = \alpha_D + \frac{3N(K-1)}{2}, \quad \beta'_D = \beta_D + \frac{1}{4}\sum_{n=1}^{N}\sum_{k=1}^{K-1}\left(\frac{(x_{n,k+1}-x_{n,k})^2 + (y_{n,k+1}-y_{n,k})^2 + (z_{n,k+1}-z_{n,k})^2}{\mathit{\Delta}_\mathrm{k}}\right). \quad \text{(S95)}$$

#### S5.5. Sampling the molecule loads

In the next step, where we sample the loads of molecules $\{b_n\}_n$, we sample from $\mathbb{P}(\{b_n\}_n|D, \{\mu_m^{\mathrm{mol}}, \mu_m^{\mathrm{back}}\}_m, \{\overline{x}_n, \overline{y}_\mathrm{n}, \overline{z}_n\}_n, \overline{\mathit{\Delta}}, \overline{s})$ which simplifies to $\mathbb{P}\left(\{b_n\}_\mathrm{n}|\{\mu_m^{\mathrm{mol}}, \mu_m^{\mathrm{back}}\}_m, \{\overline{x}_n, \overline{y}_n, \overline{z}_n\}_n, \overline{\mathit{\Delta}}, \overline{s}\right)$; variable dependencies are also shown graphically in Fig. 6. Based on these dependencies, the marginal posterior can be written as follows

$$\begin{aligned}\mathbb{P}\left(\{b_n\}_n|\{\mu_m^{\mathrm{mol}}, \mu_m^{\mathrm{back}}\}_m, \{\overline{x}_n, \overline{y}_n, \overline{z}_n\}_n, \overline{\mathit{\Delta}}, \overline{s}\right) \propto\ & \mathbb{P}\left(\overline{\mathit{\Delta}}|\{\mu_m^{\mathrm{mol}}, \mu_m^{\mathrm{back}}\}_m, \{b_n, \overline{x}_n, \overline{y}_n, \overline{z}_n\}_n\right) \\ & \times \mathbb{P}\left(\overline{s}|\{\mu_m^{\mathrm{mol}}, \mu_m^{\mathrm{back}}\}_m, \{b_n, \overline{x}_n, \overline{y}_n, \overline{z}_n\}_n\right) \\ & \times \mathbb{P}\left(\{b_n\}_n\right).\end{aligned} \quad \text{(S96)}$$

We could, in principle, sample each load successively, that is from

$$\begin{aligned}\mathbb{P}\left(b_{n'}|\{\mu_m^{\mathrm{mol}}, \mu_m^{\mathrm{back}}\}_m, \{\overline{x}_n, \overline{y}_n, \overline{z}_n\}_n, \{b_n\}_{n\neq n'}, \overline{\mathit{\Delta}}, \overline{s}\right) \propto\ & \mathbb{P}\left(\overline{\mathit{\Delta}}|\{\mu_m^{\mathrm{mol}}, \mu_m^{\mathrm{back}}\}_m, \{b_n, \overline{x}_n, \overline{y}_n, \overline{z}_n\}_n\right) \\ & \times \mathbb{P}\left(\overline{s}|\{\mu_m^{\mathrm{mol}}, \mu_m^{\mathrm{back}}\}_m, \{b_n, \overline{x}_n, \overline{y}_n, \overline{z}_n\}_n\right) \\ & \times \mathbb{P}\left(b_{n'}\right).\end{aligned} \quad \text{(S97)}$$

In practice, we have found that this gives rise to poor mixing of our MCMC chain.

The best mixing would be achieved if we could sample all loads simultaneously. This can be done by calculating the posteriors of all configurations of loads

$$\begin{aligned} B_1 &= [0,\ 0, \dots, 0] \quad , \quad P_1 = \mathbb{P}\left(B_1|\{\mu_m^{\mathrm{mol}}, \mu_m^{\mathrm{back}}\}_m, \dots\right) \\ B_2 &= [1,\ 0, \dots, 0] \quad , \quad P_2 = \mathbb{P}\left(B_2|\{\mu_m^{\mathrm{mol}}, \mu_m^{\mathrm{back}}\}_m, \dots\right) \\ B_3 &= [0,\ 1, \dots, 0] \quad , \quad P_3 = \mathbb{P}\left(B_3|\{\mu_m^{\mathrm{mol}}, \mu_m^{\mathrm{back}}\}_m, \dots\right) \\ &\vdots \\ B_{2^N} &= [1,\ 1, \dots, 1] \quad , \quad P_{2^N} = \mathbb{P}\left(B_{2^N}|\{\mu_m^{\mathrm{mol}}, \mu_m^{\mathrm{back}}\}_m, \dots\right). \end{aligned} \quad \text{(S98)}$$

and construct the categorical distribution and sample the configuration of loads

$$\{b_n\}_n|\{\mu_m^{\mathrm{mol}}, \mu_m^{\mathrm{back}}\}_m, \{\overline{x}_n, \overline{y}_n, \overline{z}_n\}_n, \overline{\Delta}, \overline{s} \sim \mathbf{Cat}_{[B_1, B_2, B_3, \dots, B_{2^N}]}\left(P_1, P_2, P_3, \dots, P_{2^N}\right). \quad \text{(S99)}$$

The problem is that this categorical distribution has $2^N$ arguments, i.e., each load can be 0 or 1. The computational cost associated to calculating these probabilities is prohibitively high.

For this reason, we compromise. We pick a fixed number of loads at random (from a uniform discrete distribution with $N$ outcomes). We update these simultaneously and repeat for the remainder of the loads until all loads have been updated.

Concretely, we define a random sets of loads $\{b_{n'}\}_{n'}$ where $n' = 1 \dots, N'$ and apply direct sampling to these. The posterior over this smaller set of loads that we update simultaneously is $\mathbb{P}\left(\{b_{n'}\}_{n'}|\{\mu_m^{\mathrm{mol}}, \mu_m^{\mathrm{back}}\}_m, \{\overline{x}_n, \overline{y}_n, \overline{z}_n, b_{n,n\neq n'}\}_n, \overline{\Delta}, \overline{s}\right)$. So, the conditional probability distribution can be written as

$$\begin{aligned}\mathbb{P}\left(\{b_{n'}\}_{n'}|\{\mu_m^{\text{mol}},\mu_m^{\text{back}}\}_m,\{\overline{x}_n,\overline{y}_n,\overline{z}_n,b_{n,n\neq n'}\}_n,\overline{\Delta},\overline{s}\right) &\propto \mathbb{P}\left(\overline{\Delta}|\{\mu_m^{\text{mol}},\mu_m^{\text{back}}\}_m,\{b_{n,n\neq n'},\overline{x}_n,\overline{y}_n,\overline{z}_n\}_n,\{b_{n'}\}_{n'}\right)\\ &\times \mathbb{P}\left(\overline{s}|\{\mu_m^{\text{mol}},\mu_m^{\text{back}}\}_m,\{b_{n,n\neq n'},\overline{x}_n,\overline{y}_n,\overline{z}_n\}_n,\{b_{n'}\}_{n'}\right)\\ &\times \mathbb{P}\left(\{b_{n'}\}_{n'}\right)\\ = \left[\prod_{k=1}^{K-1}\mathbf{Exp}\left(\Delta_k;\sum_{m=1}^{M}\mu_{m,k}\right)\right]\left[\prod_{k=1}^{K}\mathbf{Cat}\left(s_k;\frac{\mu_{1,k}}{\sum_{m=1}^{M}\mu_{m,k}},\dots,\frac{\mu_{M,k}}{\sum_{m=1}^{M}\mu_{m,k}}\right)\right]&\left[\prod_{n'=1}^{N'}\mathbf{Bernoulli}\left(b_{n'};\frac{1}{1+\frac{N-1}{\gamma_b}}\right)\right]\\ = \left(\frac{\mu_{s_K,K}}{\sum_{m=1}^{M}\mu_{m,K}}\right)\left[\prod_{k=1}^{K-1}\mu_{s_k,k}\exp\left(-\Delta_k\sum_{m=1}^{M}\mu_{m,k}\right)\right]&\left[\prod_{n'=1}^{N'}\mathbf{Bernoulli}\left(b_{n'};\frac{1}{1+\frac{N-1}{\gamma_b}}\right)\right]\end{aligned} \tag{S100}$$

where, $\mu_{m,k}=\mu_m^{\text{back}}+\mu_m^{\text{mol}}\sum_{n=1}^{N}b_n\ \text{PSF}_m\left(x_{n,k},y_{n,k},z_{n,k}\right)$.

Again, to simplify the computational calculations, we calculate the logarithmic of this conditional probability distribution as

$$\begin{aligned}\log\mathbb{P}\Big(\{b_{n'}\}_{n'}|\{\mu_m^{\text{mol}},\mu_m^{\text{back}}\}_m,\{\overline{x}_n,\overline{y}_n,\overline{z}_n,b_{n,n\neq n'}\}_n,\overline{\Delta},\overline{s}\Big) &= \log\left(\frac{\mu_{s_K,K}}{\sum_{m=1}^{M}\mu_{m,K}}\right)+\left[\sum_{k=1}^{K-1}\log\left(\mu_{s_k,k}\right)-\Delta_k\sum_{m=1}^{M}\mu_{m,k}\right]\\ &+N'\log\left(1-\frac{1}{1+\frac{N-1}{\gamma_b}}\right)-\left(\sum_{n'=1}^{N'}b_{n'}\right)\log\left(\frac{N-1}{\gamma_b}\right)\\ &+\text{Constant}.\end{aligned} \tag{S101}$$

#### S5.6. Joint sampling of molecular brightness and background photon emission rates

Finally, after we update locations of molecules as well as loads, we update the molecular brightnesses and background photon emission rates related to all of confocal volumes $\{\mu_m^{\text{mol}}\}_m$ and $\{\mu_m^{\text{back}}\}_m$ by sampling from the corresponding conditional $\mathbb{P}\left(\{\mu_m^{\text{mol}},\mu_m^{\text{back}}\}_m|D,\{b_n,\overline{x}_n,\overline{y}_n,\overline{z}_n\}_n,\overline{\Delta},\overline{s}\right)$, which simplifies to $\mathbb{P}\left(\{\mu_m^{\text{mol}},\mu_m^{\text{back}}\}_m|\{b_n,\overline{x}_n,\overline{y}_n,\overline{z}_n\}_n,\overline{\Delta},\overline{s}\right)$ and $m=1,\dots,M$ where the $M$ is the total number confocal volumes.

$$\begin{aligned}\mathbb{P}\left(\{\mu_m^{\text{mol}},\mu_m^{\text{back}}\}_m|\{b_n,\overline{x}_n,\overline{y}_n,\overline{z}_n\}_n,\overline{\Delta},\overline{s}\right) &\propto \mathbb{P}\left(\overline{\Delta}|\{\mu_m^{\text{mol}},\mu^{\text{back}}\}_m,\{b_n,\overline{x}_n,\overline{y}_n,\overline{z}_n\}_n\right)\\ &\times \mathbb{P}\left(\overline{s}|\{\mu_m^{\text{mol}},\mu^{\text{back}}\}_m,\{b_n,\overline{x}_n,\overline{y}_n,\overline{z}_n\}_n\right)\\ &\times \mathbb{P}\left(\{\mu_m^{\text{mol}}\}_m\right)\mathbb{P}\left(\{\mu_m^{\text{back}}\}_m\right)\quad,\quad m=1,\dots,M.\end{aligned} \tag{S102}$$

We carry over this sampling using a Metropolis-Hastings update where proposals for $\mu_m^{\text{mol}}$ and $\mu_m^{\text{back}}$ are computed according to

$$\begin{aligned}\left(\mu_m^{\text{mol}}\right)^{\text{prop}} &\sim \mathbf{Gamma}\left(\alpha_{\text{mol}}^{\text{prop}},\frac{\left(\mu_m^{\text{mol}}\right)^{\text{old}}}{\alpha_{\text{mol}}^{\text{prop}}}\right)\quad,\quad m=1,\dots,M\\ \left(\mu_m^{\text{back}}\right)^{\text{prop}} &\sim \mathbf{Gamma}\left(\alpha_{\text{back}}^{\text{prop}},\frac{\left(\mu_m^{\text{back}}\right)^{\text{old}}}{\alpha_{\text{back}}^{\text{prop}}}\right)\quad,\quad m=1,\dots,M\end{aligned} \tag{S103}$$

where $\left(\mu_m^{\text{mol}}\right)^{\text{old}}$ and $\left(\mu_m^{\text{back}}\right)^{\text{old}}$ denote the existing samples. So, by defining the proposal probability distributions we can calculate the ratio

$$
\begin{aligned}
r = {} & \frac{\mathbb{P}\left(\overline{\Delta}|\{(\mu_m^{\mathrm{mol}})^{\mathrm{prop}},(\mu_m^{\mathrm{back}})^{\mathrm{prop}}\}_m,\{b_n,\overline{x}_n,\overline{y}_n,\overline{z}_n\}_n\right)\mathbb{P}\left(\overline{s}|\{(\mu_m^{\mathrm{mol}})^{\mathrm{prop}},(\mu_m^{\mathrm{back}})^{\mathrm{prop}}\}_m,\{b_n,\overline{x}_n,\overline{y}_n,\overline{z}_n\}_n\right)}{\mathbb{P}\left(\overline{\Delta}|\{(\mu_m^{\mathrm{mol}})^{\mathrm{old}},(\mu_m^{\mathrm{back}})^{\mathrm{old}}\}_m,\{b_n,\overline{x}_n,\overline{y}_n,\overline{z}_n\}_n\right)\mathbb{P}\left(\overline{s}|\{(\mu_m^{\mathrm{mol}})^{\mathrm{old}},(\mu_m^{\mathrm{back}})^{\mathrm{old}}\}_m,\{b_n,\overline{x}_n,\overline{y}_n,\overline{z}_n\}_n\right)} \\
& \times \prod_{m=1}^{M}\frac{\mathbb{P}\left((\mu_m^{\mathrm{mol}})^{\mathrm{prop}}\right)}{\mathbb{P}\left((\mu_m^{\mathrm{mol}})^{\mathrm{old}}\right)}\frac{\mathbb{P}\left((\mu_m^{\mathrm{mol}})^{\mathrm{old}}\,|\,(\mu_m^{\mathrm{mol}})^{\mathrm{prop}}\right)}{\mathbb{P}\left((\mu_m^{\mathrm{mol}})^{\mathrm{prop}}\,|\,(\mu_m^{\mathrm{mol}})^{\mathrm{old}}\right)}\frac{\mathbb{P}\left((\mu_m^{\mathrm{back}})^{\mathrm{prop}}\right)}{\mathbb{P}\left((\mu_m^{\mathrm{back}})^{\mathrm{old}}\right)}\frac{\mathbb{P}\left((\mu_m^{\mathrm{back}})^{\mathrm{old}}\,|\,(\mu_m^{\mathrm{back}})^{\mathrm{prop}}\right)}{\mathbb{P}\left((\mu_m^{\mathrm{back}})^{\mathrm{prop}}\,|\,(\mu_m^{\mathrm{back}})^{\mathrm{old}}\right)} \\
= {} & \prod_{k=1}^{K-1}\left[\exp\left(\Delta_k\left[\sum_{m=1}^{M}\left((\mu_m^{\mathrm{back}})^{\mathrm{old}}-(\mu_m^{\mathrm{back}})^{\mathrm{prop}}\right)+\left((\mu_m^{\mathrm{mol}})^{\mathrm{old}}-(\mu_m^{\mathrm{mol}})^{\mathrm{prop}}\right)\sum_{n=1}^{N}b_n\mathrm{PSF}_m\left(x_{n,k},y_{n,k},z_{n,k}\right)\right]\right)\right. \\
& \times \frac{(\mu_{s_k}^{\mathrm{back}})^{\mathrm{prop}}+(\mu_{s_k}^{\mathrm{mol}})^{\mathrm{prop}}\sum_{n=1}^{N}b_n\mathrm{PSF}_{s_k}\left(x_{n,k},y_{n,k},z_{n,k}\right)}{(\mu_{s_k}^{\mathrm{back}})^{\mathrm{old}}+(\mu_{s_k}^{\mathrm{mol}})^{\mathrm{old}}\sum_{\mathrm{n}=1}^{N}b_n\mathrm{PSF}_{s_k}\left(x_{n,k},y_{n,k},z_{\mathrm{n,k}}\right)} \\
& \times \prod_{m=1}^{M}\left[\left(\frac{(\mu_m^{\mathrm{mol}})^{\mathrm{old}}}{(\mu_m^{\mathrm{mol}})^{\mathrm{prop}}}\right)^{2\alpha_{\mathrm{mol}}^{\mathrm{prop}}-\alpha_{\mathrm{mol}}}\exp\left(\alpha_{\mathrm{mol}}^{\mathrm{prop}}\left(\frac{(\mu_m^{\mathrm{mol}})^{\mathrm{prop}}}{(\mu_m^{\mathrm{mol}})^{\mathrm{old}}}-\frac{(\mu_m^{\mathrm{mol}})^{\mathrm{old}}}{(\mu_m^{\mathrm{mol}})^{\mathrm{prop}}}\right)+\frac{(\mu_m^{\mathrm{mol}})^{\mathrm{old}}-(\mu_m^{\mathrm{mol}})^{\mathrm{prop}}}{\beta_{\mathrm{mol}}}\right)\right. \\
& \left.\times\left(\frac{(\mu_m^{\mathrm{back}})^{\mathrm{old}}}{(\mu_m^{\mathrm{back}})^{\mathrm{prop}}}\right)^{2\alpha_{\mathrm{back}}^{\mathrm{prop}}-\alpha_{\mathrm{back}}}\exp\left(\alpha_{\mathrm{back}}^{\mathrm{prop}}\left(\frac{(\mu_m^{\mathrm{back}})^{\mathrm{prop}}}{(\mu_m^{\mathrm{back}})^{\mathrm{old}}}-\frac{(\mu_m^{\mathrm{back}})^{\mathrm{old}}}{(\mu_m^{\mathrm{back}})^{\mathrm{prop}}}\right)+\frac{(\mu_m^{\mathrm{back}})^{\mathrm{old}}-(\mu_m^{\mathrm{back}})^{\mathrm{prop}}}{\beta_{\mathrm{back}}}\right)\right].
\end{aligned}
\tag{S104}
$$

As it is convenient in computation, we compute the log of above ratio which reads

$$
\begin{aligned}
\log r = {} & \sum_{k=1}^{K-1}\left[\Delta_k\left(\sum_{m=1}^{M}\left((\mu_m^{\mathrm{back}})^{\mathrm{old}}-(\mu_m^{\mathrm{back}})^{\mathrm{prop}}\right)+\left((\mu_m^{\mathrm{mol}})^{\mathrm{old}}-(\mu_m^{\mathrm{mol}})^{\mathrm{prop}}\right)\sum_{n=1}^{N}b_n\mathrm{PSF}_m\left(x_{n,k},y_{n,\mathrm{k}},z_{n,k}\right)\right)\right. \\
& \left.+\log\left(\frac{(\mu_{s_k}^{\mathrm{back}})^{\mathrm{prop}}+(\mu_{s_k}^{\mathrm{mol}})^{\mathrm{prop}}\sum_{\mathrm{n}=1}^{N}b_n\mathrm{PSF}_{s_k}\left(x_{n,k},y_{n,k},z_{n,k}\right)}{(\mu_{s_k}^{\mathrm{back}})^{\mathrm{old}}+(\mu_{s_k}^{\mathrm{mol}})^{\mathrm{old}}\sum_{n=1}^{N}b_n\mathrm{PSF}_{s_k}\left(x_{n,k},y_{n,k},z_{n,k}\right)}\right)\right] \\
& +\sum_{m=1}^{M}\left[(2\alpha_{\mathrm{mol}}^{\mathrm{prop}}-\alpha_{\mathrm{mol}})\log\left(\frac{(\mu_m^{\mathrm{mol}})^{\mathrm{old}}}{(\mu_m^{\mathrm{mol}})^{\mathrm{prop}}}\right)+\alpha_{\mathrm{mol}}^{\mathrm{prop}}\left(\frac{(\mu_m^{\mathrm{mol}})^{\mathrm{prop}}}{(\mu_m^{\mathrm{mol}})^{\mathrm{old}}}-\frac{(\mu_m^{\mathrm{mol}})^{\mathrm{old}}}{(\mu_m^{\mathrm{mol}})^{\mathrm{prop}}}\right)+\frac{(\mu_m^{\mathrm{mol}})^{\mathrm{old}}-(\mu_m^{\mathrm{mol}})^{\mathrm{prop}}}{\beta_{\mathrm{mol}}}\right. \\
& \left.+(2\alpha_{\mathrm{back}}^{\mathrm{prop}}-\alpha_{\mathrm{back}})\log\left(\frac{(\mu_m^{\mathrm{back}})^{\mathrm{old}}}{(\mu_m^{\mathrm{back}})^{\mathrm{prop}}}\right)+\alpha_{\mathrm{back}}^{\mathrm{prop}}\left(\frac{(\mu_m^{\mathrm{back}})^{\mathrm{prop}}}{(\mu_m^{\mathrm{back}})^{\mathrm{old}}}-\frac{(\mu_m^{\mathrm{back}})^{\mathrm{old}}}{(\mu_m^{\mathrm{back}})^{\mathrm{prop}}}\right)+\frac{(\mu_m^{\mathrm{back}})^{\mathrm{old}}-(\mu_m^{\mathrm{back}})^{\mathrm{prop}}}{\beta_{\mathrm{back}}}\right].
\end{aligned}
\tag{S105}
$$

#### S5.7. Track the loads and the trajectories of molecules

After sampling all of the random variables in one iteration of the Gibbs sampling scheme, we need to find a way to sort the sampled values for our random variables. The most important variables in this study are the trajectories of the active molecules and the loads which represent the active molecules $\{\overline{x}_n,\overline{y}_n,\overline{z}_n\}_{n,b_n=1}$ . As we show in Fig. S14(a), due to the exchangeability of the sampler, the values of variables (molecular trajectories and the loads) will swap and yield identical posteriors. This is a generic characteristic of Bayesian nonparametric approaches. [33–35] For this reason, we need to find a way to relate the sampled trajectories and loads of each molecule at any iteration of the Gibbs sampler (iterations are shown with index $i$).

To be able to distinguish the trajectories and loads, we calculate the distances of each trajectory from the others. At the same time, we need to consider the effect of the PSFs, molecular brightnesses and the loads. To do this, we consider the molecular emission rate of each individual model molecule at any given time $\eta_{n,k}$,

$$
\eta_{n,k}=b_n\sum_{m=1}^{M}\mu_m^{\mathrm{mol}}\mathrm{PSF}_m\left(x_{n,k},y_{n,k},z_{n,k}\right)\qquad,\qquad k=1,\ldots,K-1. \tag{S106}
$$

For each molecule, $\overline{\eta}_n=(\eta_{n,1},\ldots,\eta_{n,K})$ represents the total emission rate of that molecule detected over all confocal volumes any time we detect a photon at time $t_k$.

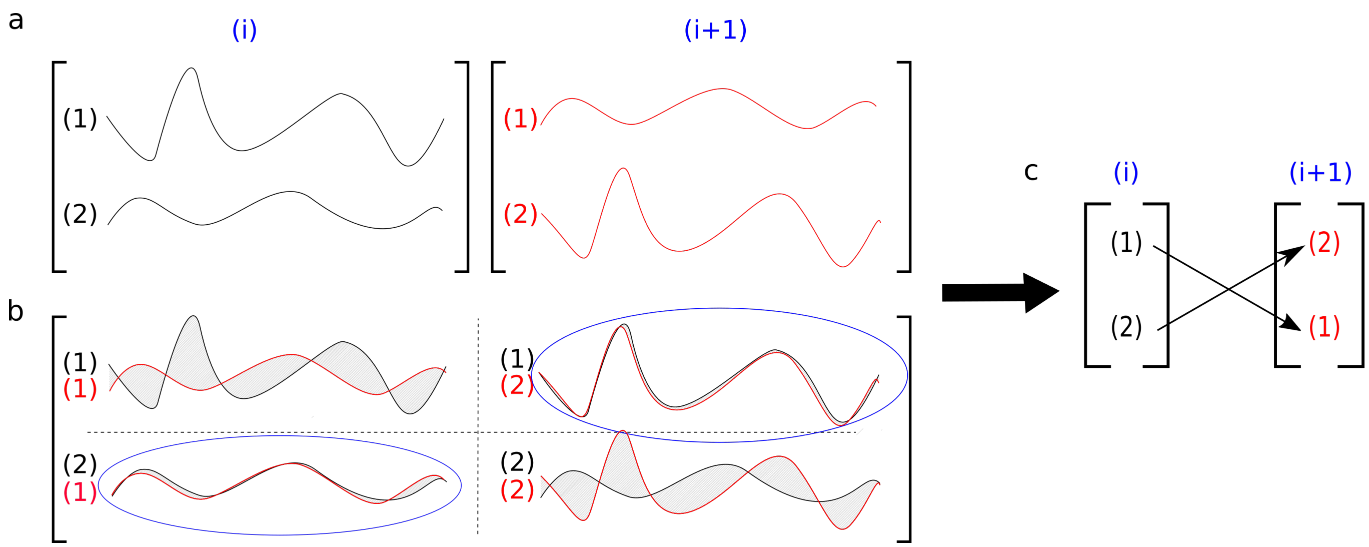


FIG. S14. **Hungarian algorithm**. **(a)** A simple example of the Hungarian algorithm when we have only two molecular trajectories sampled at iteration $i$ and $i+1$ of the Gibbs sampler. At each iteration of the Gibbs sampler, we sample all of the random variables but here we only illustrate the molecular trajectories. In the first iteration, the first trajectory can be assigned to molecule 1 and the second to molecule 2 and vice versa for the second iteration. We emphasize this problem is fundamental (and not a limitation of our approach) as the posterior is invariant with respect to label swapping. **(b)** We compare trajectories to re-assign molecular identity labels. This can be achieved by minimizing the area between the trajectories. The area here serves as a post-processing cost-function. **(c)** The Hungarian algorithm results in re-assigned labels for trajectories.

Using $\overline{\eta}_n$, we now construct the cost matrix (the matrix of distances between trajectories incorporating information on molecular emission rate) demanded of the Hungarian algorithm [36]

$$\text{Cost matrix} = \begin{bmatrix} d_{1,1} & d_{1,2} & \dots & d_{1,N} \\ d_{1,1} & d_{1,2} & \dots & d_{1,N} \\ \vdots & \vdots & \ddots & \vdots \\ d_{N,1} & d_{N,2} & \dots & d_{N,N} \end{bmatrix} \tag{S107}$$

where each element of this matrix is equal to

$$d_{n,n'} = \sum_{k=1}^{K-1} |\eta_{n,k} - \eta_{n',k}| \,. \tag{S108}$$

The goal is then to associate elements of the cost matrix in Eq. (S107) (each row corresponds to a trajectory and its load of the fixed values (pivot) and each column corresponds to a trajectory and its load at iteration $i$) that the sum of them is minimized using the Hungarian algorithm [36].

We are flexible to choose any pivot. Here, we choose the trajectories and loads of the MAP computed sample as our pivot.

#### S5.8. Maximum posteriori (MAP) sample

For each iteration of our MCMC chain, we have one sample from the posterior probability distribution. We can process these samples to find the MAP sample which is used as the pivot in the Sec. S.5.7. To compute the MAP sample, we calculate the joint posterior over all random variables

$$
\begin{aligned}
&\mathbb{P}\left(D, \{\mu_m^{\text{mol}}, \mu_m^{\text{back}}\}_m, \{b_n, \overline{x}_n, \overline{y}_n, \overline{z}_n\}_n | \overline{\mathit{\Delta}}, \overline{s}\right) \\
&\quad \propto \left[\prod_{k=1}^{K-1} \mathbb{P}\left(\mathit{\Delta}_k | \{b_n, x_{n,k}, y_{n,k}, z_{n,k}\}_n, \{\mu_m^{\text{mol}}, \mu_m^{\text{back}}\}_m\right) \mathbb{P}\left(s_k | \{b_n, x_{n,k}, y_{n,k}, z_{n,k}\}_n, \{\mu_m^{\text{mol}}, \mu_m^{\text{back}}\}_m\right)\right] \\
&\quad \times \left[\prod_{n=1}^{N}\left[\prod_{k=1}^{K-1} \mathbb{P}\left(x_{n,k+1} | x_{n,k}, D, \mathit{\Delta}_k\right) \mathbb{P}\left(y_{n,k+1} | y_{n,k}, D, \mathit{\Delta}_k\right) \mathbb{P}\left(z_{n,k+1} | z_{n,k}, D, \mathit{\Delta}_k\right)\right] \mathbb{P}\left(x_{n,1}\right) \mathbb{P}\left(y_{n,1}\right) \mathbb{P}\left(z_{n,1}\right) \mathbb{P}\left(\{b_n\}_n\right)\right] \\
&\quad \times \mathbb{P}\left(D\right) \prod_{m=1}^{M} \mathbb{P}\left(\mu_m^{\text{mol}}\right) \mathbb{P}\left(\mu_m^{\text{back}}\right) \\
&\quad = \prod_{k=1}^{K-1}\left(\mu_{s_k}^{\text{back}} + \mu_{s_k}^{\text{mol}} \sum_{n=1}^{N} b_n \text{PSF}_{s_k}\left(x_{n,k}, y_{n,k}, z_{n,k}\right)\right) \exp\left(-\mathit{\Delta}_k\left[\sum_{m=1}^{M} \mu_m^{\text{back}} + \mu_m^{\text{mol}} \sum_{n=1}^{N} b_n \text{PSF}_m\left(x_{n,k}, y_{n,k}, z_{n,k}\right)\right]\right) \\
&\quad \times \left[\prod_{n=1}^{N}\left[\prod_{k=1}^{K-1} \frac{1}{\left(4\pi D \mathit{\Delta}_k\right)^{\frac{3}{2}}} \exp\left(-\frac{\left(x_{n,k+1}-x_{n,k}\right)^2 + \left(y_{n,k+1}-y_{n,k}\right)^2 + \left(z_{n,k+1}-z_{n,k}\right)^2}{4D\mathit{\Delta}_k}\right)\right]\right. \\
&\quad \left. \times \frac{1}{\sqrt{8\pi^3 \sigma_{x_0}^2 \sigma_{y_0}^2 \sigma_{z_0}^2}} \exp\left(-\frac{(x_{n,1}-x_0)^2}{2\sigma_{x_0}^2} + \frac{(y_{n,1}-y_0)^2}{2\sigma_0^2} + \frac{(z_{n,1}-z_0)^2}{2\sigma_{z_0}^2}\right)\left(\frac{1}{1+\frac{N-1}{\gamma_b}}\right)^{b_n}\left(1-\frac{1}{1+\frac{N-1}{\gamma_b}}\right)^{1-b_n}\right] \\
&\quad \times \mathbf{InvGamma}(D; \alpha_D, \beta_D) \prod_{m=1}^{M} \mathbf{Gamma}(\mu_m^{\text{mol}}; \alpha_{\text{mol}}, \beta_{\text{mol}}) \mathbf{Gamma}(\mu_m^{\text{back}}; \alpha_{\text{back}}, \beta_{back})
\end{aligned}
\tag{S109}
$$

where $\mu_{m,k} = \mu_m^{\text{back}} + \mu_m^{\text{mol}} \sum_n b_n \mathbf{PSF}_m(x_{n,k}, y_{n,k}, z_{n,k})$. Of course, to avoid numerical underflow, we compute directly the logarithm $\log \mathbb{P}\left(D, \{\mu_m^{\text{mol}}, \mu_m^{\text{back}}\}_m, \{b_n, \overline{x}_n, \overline{y}_n, \overline{z}_n\}_n | \overline{\mathit{\Delta}}, \overline{s}\right)$.

TABLE S2. Summary of notation.

| Description | Variable | Units |
|---|---|---|
| Diffusion coefficient | $D$ | $\mu$m$^2$s$^{-1}$ |
| $\alpha$ parameter of the diffusion coefficient prior | $\alpha_D$ | - |
| $\beta$ parameter of the diffusion coefficient prior | $\beta_D$ | $\mu$m$^2$s$^{-1}$ |
| Total time trace duration | $T_{\text{total}}$ | s |
| Molecular brightness at the center of the confocal volume m | $\mu_m^{\text{mol}}$ | photons s$^{-1}$ |
| $\alpha$ parameter of the molecular brightness's prior | $\alpha_{\text{mol}}$ | - |
| $\beta$ parameter of the molecular brightness's prior | $\beta_{\text{mol}}$ | photons s$^{-1}$ |
| Proposal parameter of the molecule photon emission rate | $\alpha_{\text{mol}}^{\text{prop}}$ | - |
| Combined photon emission rates of all molecules at time $t_k$ of the confocal volume m | $\mu_{m,k}$ | photons s$^{-1}$ |
| Background photon emission rate of confocal volume m | $\mu_m^{\text{back}}$ | photons s$^{-1}$ |
| $\alpha$ parameter of the background photon emission rate's prior | $\alpha_{\text{back}}$ | - |
| $\beta$ parameter of the background photon emission rate's prior | $\beta_{\text{back}}$ | photons s$^{-1}$ |
| Proposal parameter of the background photon emission rate | $\alpha_{\text{back}}^{\text{prop}}$ | - |
| Minor and major semi-axes of confocal PSF (x axis) of confocal volume m | $\omega_{m,x}$, $\omega_{m,y}$, $\omega_{m,z}$ | $\mu$m |
| Location of molecule n at time $t_k$ in x, y and z coordinates | $x_{n,k}$, $y_{n,k}$, $z_{n,k}$ | $\mu$m |
| Recorded photon inter-arrival by all of the detector | $\Delta_k$ | s |
| Label on the detected photon at time $t_k$ | $s_k$ | - |
| Load variable for molecule $n$ | $b_n$ | - |
| Prior weight for $b_n$ | $q_n$ | - |
| Parameter of hyperprior $q_n$ | $\gamma_b$ | - |
| Upper bound for the number of model molecules | $N$ | - |
| Number of confocal volumes | $\mathbf{M}$ | - |
| Mean values of initial molecule position's prior in the x, y and z axes | $x_0$, $y_0$, $z_0$ | $\mu$m |
| Variances of the initial molecule position's prior in the x, y and z axes | $\sigma_{x_0}^2$, $\sigma_{y_0}^2$, $\sigma_{z_0}^2$ | $\mu$m |
| Periodic boundaries in the x, y and z axes (focal plane) | $L_x$, $L_y$, $L_z$ | $\mu$m |

TABLE S3. List of abbreviations.

| Phrase | Abbreviation |
|---|---|
| Fluorescence confocal microscopy | FCM |
| Fluorescence correlation spectroscopy | FCS |
| Region of interest | ROI |
| Hamiltonian Monte Carlo | HMC |
| Point spread function | PSF |
| Three dimensional Gaussian | 3DG |
| Two dimensional Gaussian-Lorentzian | 2DGL |
| Two dimensional Gaussian-Cylindrical | 2DGC |
| Markov chain Monte Carlo | MCMC |
| Graphical user interface | GUI |
| Excitation profile | EXC |

TABLE S4. Probability distributions used and their densities. Here, the corresponding random variables are denoted by $x$.

| Distribution | Notation | Probability density function | Mean value | Variance/Covariance |
|---|---|---|---|---|
| Normal | $\mathbf{Normal}(\mu, \sigma^2)$ | $\frac{1}{\sqrt{2\pi\sigma^2}} e^{-\frac{(x-\mu)^2}{2\sigma^2}}$ | $\mu$ | $\sigma^2$ |
| Exponential | $\mathbf{Exp}(\mu)$ | $\mu e^{-\mu x}$ | $\mu^{-1}$ | $\mu^{-2}$ |
| Gamma | $\mathbf{Gamma}(\alpha, \beta)$ | $\frac{1}{\Gamma(\alpha)\beta^\alpha} x^{\alpha-1} e^{-\frac{x}{\beta}}$ | $\alpha\beta$ | $\alpha\beta^2$ |
| Inverse Gamma | $\mathbf{InvGamma}(\alpha, \beta)$ | $\frac{\beta^\alpha}{\Gamma(\alpha)} x^{-\alpha-1} e^{-\frac{\beta}{x}}$ | $\frac{\beta}{\alpha-1}$ | $\frac{\beta^2}{(\alpha-1)^2(\alpha-2)}$ |
| Beta | $\mathbf{Beta}(\alpha, \beta)$ | $\frac{\Gamma(\alpha+\beta)}{\Gamma(\alpha)\Gamma(\beta)} x^{\alpha-1}(1-x)^{\beta-1}$ | $\frac{\alpha}{\alpha+\beta}$ | $\frac{\alpha\beta}{(\alpha+\beta)^2(\alpha+\beta+1)}$ |
| Bernoulli | $\mathbf{Bernoulli}(q)$ | $(q-1)\delta_0(x) + q\delta_1(x)$ | $q$ | $q(1-q)$ |

### S6. Summary of notation, abbreviations, parameters and other quantities

TABLE S5. Parameter values used in the generation of synthetic traces. Choices are listed according to figures. Since we consider four confocal volumes $m = 1, 2, 3, 4$, we have four PSFs, molecular brightnesses and background photon emission rates for any of the respective figures. Additional values are listed in Table. S6

| | $L_{xy}$ | $L_z$ | $\text{PSF}_m$ | $[\text{C}_{m,x}\ , \text{C}_{m,y}\ , \text{C}_{m,z}]$ | $[\omega_{m,x}\ , \omega_{m,y}\ , \omega_{m,z}]$ | $N$ | $D$ | $\mu_m^{\text{mol}}$ | $\mu_m^{\text{back}}$ | $\text{T}_{\text{total}}$ |
|---|---|---|---|---|---|---|---|---|---|---|
| Units | $\mu$m | $\mu$m | - | $\mu$m | $\mu$m | - | $\mu\text{m}^2\text{s}^{-1}$ | pht $\text{s}^{-1}$ | pht $\text{s}^{-1}$ | s |
| Fig. 2(a) | 6 | 8 | $\begin{bmatrix}\text{3DG}\\\text{3DG}\\\text{3DG}\\\text{3DG}\end{bmatrix}$ | $\begin{bmatrix}-0.3 & 0.0 & -0.5\\0.3 & 0.0 & -0.5\\0.0 & -0.3 & 0.5\\0.0 & 0.3 & 0.5\end{bmatrix}$ | $\begin{bmatrix}0.3 & 0.3 & 1.5\\0.3 & 0.3 & 1.5\\0.3 & 0.3 & 1.5\\0.3 & 0.3 & 1.5\end{bmatrix}$ | 1 | 0.1 | $\begin{bmatrix}5\\5\\5\\5\end{bmatrix}\times 10^4$ | $\begin{bmatrix}1\\1\\1\\1\end{bmatrix}\times 10^3$ | 0.26 |
| Fig. 2(b) | 6 | 8 | $\begin{bmatrix}\text{3DG}\\\text{3DG}\\\text{3DG}\\\text{3DG}\end{bmatrix}$ | $\begin{bmatrix}-0.3 & 0.0 & -0.5\\0.3 & 0.0 & -0.5\\0.0 & -0.3 & 0.5\\0.0 & 0.3 & 0.5\end{bmatrix}$ | $\begin{bmatrix}0.3 & 0.3 & 1.5\\0.3 & 0.3 & 1.5\\0.3 & 0.3 & 1.5\\0.3 & 0.3 & 1.5\end{bmatrix}$ | 1 | 1 | $\begin{bmatrix}5\\5\\5\\5\end{bmatrix}\times 10^4$ | $\begin{bmatrix}1\\1\\1\\1\end{bmatrix}\times 10^3$ | 0.41 |
| Fig. 2(c) | 6 | 8 | $\begin{bmatrix}\text{3DG}\\\text{3DG}\\\text{3DG}\\\text{3DG}\end{bmatrix}$ | $\begin{bmatrix}-0.3 & 0.0 & -0.5\\0.3 & 0.0 & -0.5\\0.0 & -0.3 & 0.5\\0.0 & 0.3 & 0.5\end{bmatrix}$ | $\begin{bmatrix}0.3 & 0.3 & 1.5\\0.3 & 0.3 & 1.5\\0.3 & 0.3 & 1.5\\0.3 & 0.3 & 1.5\end{bmatrix}$ | 1 | 10 | $\begin{bmatrix}5\\5\\5\\5\end{bmatrix}\times 10^4$ | $\begin{bmatrix}1\\1\\1\\1\end{bmatrix}\times 10^3$ | 1.77 |
| Fig. 3(a) | 6 | 8 | $\begin{bmatrix}\text{3DG}\\\text{3DG}\\\text{3DG}\\\text{3DG}\end{bmatrix}$ | $\begin{bmatrix}-0.3 & 0.0 & -0.5\\0.3 & 0.0 & -0.5\\0.0 & -0.3 & 0.5\\0.0 & 0.3 & 0.5\end{bmatrix}$ | $\begin{bmatrix}0.3 & 0.3 & 1.5\\0.3 & 0.3 & 1.5\\0.3 & 0.3 & 1.5\\0.3 & 0.3 & 1.5\end{bmatrix}$ | 1 | 1 | $\begin{bmatrix}5\\5\\5\\5\end{bmatrix}\times 10^4$ | $\begin{bmatrix}1\\1\\1\\1\end{bmatrix}\times 10^3$ | 0.27 |
| Fig. 3(b) | 6 | 8 | $\begin{bmatrix}\text{3DG}\\\text{3DG}\\\text{3DG}\\\text{3DG}\end{bmatrix}$ | $\begin{bmatrix}-0.3 & 0.0 & -0.5\\0.3 & 0.0 & -0.5\\0.0 & -0.3 & 0.5\\0.0 & 0.3 & 0.5\end{bmatrix}$ | $\begin{bmatrix}0.3 & 0.3 & 1.5\\0.3 & 0.3 & 1.5\\0.3 & 0.3 & 1.5\\0.3 & 0.3 & 1.5\end{bmatrix}$ | 2 | 1 | $\begin{bmatrix}5\\5\\5\\5\end{bmatrix}\times 10^4$ | $\begin{bmatrix}1\\1\\1\\1\end{bmatrix}\times 10^3$ | 0.14 |
| Fig. 3(c) | 6 | 8 | $\begin{bmatrix}\text{3DG}\\\text{3DG}\\\text{3DG}\\\text{3DG}\end{bmatrix}$ | $\begin{bmatrix}-0.3 & 0.0 & -0.5\\0.3 & 0.0 & -0.5\\0.0 & -0.3 & 0.5\\0.0 & 0.3 & 0.5\end{bmatrix}$ | $\begin{bmatrix}0.3 & 0.3 & 1.5\\0.3 & 0.3 & 1.5\\0.3 & 0.3 & 1.5\\0.3 & 0.3 & 1.5\end{bmatrix}$ | 3 | 1 | $\begin{bmatrix}5\\5\\5\\5\end{bmatrix}\times 10^4$ | $\begin{bmatrix}1\\1\\1\\1\end{bmatrix}\times 10^3$ | 0.12 |
| Fig. 4(a) | 6 | 8 | $\begin{bmatrix}\text{3DG}\\\text{3DG}\\\text{3DG}\\\text{3DG}\end{bmatrix}$ | $\begin{bmatrix}-0.3 & 0.0 & -0.5\\0.3 & 0.0 & -0.5\\0.0 & -0.3 & 0.5\\0.0 & 0.3 & 0.5\end{bmatrix}$ | $\begin{bmatrix}0.3 & 0.3 & 1.5\\0.3 & 0.3 & 1.5\\0.3 & 0.3 & 1.5\\0.3 & 0.3 & 1.5\end{bmatrix}$ | 1 | 1 | $\begin{bmatrix}1\\1\\1\\1\end{bmatrix}\times 10^4$ | $\begin{bmatrix}1\\1\\1\\1\end{bmatrix}\times 10^3$ | 1.33 |
| Fig. 4(b) | 6 | 8 | $\begin{bmatrix}\text{3DG}\\\text{3DG}\\\text{3DG}\\\text{3DG}\end{bmatrix}$ | $\begin{bmatrix}-0.3 & 0.0 & -0.5\\0.3 & 0.0 & -0.5\\0.0 & -0.3 & 0.5\\0.0 & 0.3 & 0.5\end{bmatrix}$ | $\begin{bmatrix}0.3 & 0.3 & 1.5\\0.3 & 0.3 & 1.5\\0.3 & 0.3 & 1.5\\0.3 & 0.3 & 1.5\end{bmatrix}$ | 1 | 1 | $\begin{bmatrix}5\\5\\5\\5\end{bmatrix}\times 10^4$ | $\begin{bmatrix}1\\1\\1\\1\end{bmatrix}\times 10^3$ | 0.23 |
| Fig. 4(c) | 6 | 8 | $\begin{bmatrix}\text{3DG}\\\text{3DG}\\\text{3DG}\\\text{3DG}\end{bmatrix}$ | $\begin{bmatrix}-0.3 & 0.0 & -0.5\\0.3 & 0.0 & -0.5\\0.0 & -0.3 & 0.5\\0.0 & 0.3 & 0.5\end{bmatrix}$ | $\begin{bmatrix}0.3 & 0.3 & 1.5\\0.3 & 0.3 & 1.5\\0.3 & 0.3 & 1.5\\0.3 & 0.3 & 1.5\end{bmatrix}$ | 1 | 1 | $\begin{bmatrix}1\\1\\1\\1\end{bmatrix}\times 10^5$ | $\begin{bmatrix}1\\1\\1\\1\end{bmatrix}\times 10^3$ | 0.06 |
| Fig. 5(a) | 6 | 8 | $\begin{bmatrix}\text{3DG}\\\text{3DG}\\\text{3DG}\\\text{3DG}\end{bmatrix}$ | $\begin{bmatrix}-0.01 & 0.0 & -0.01\\0.01 & 0.0 & -0.01\\0.0 & -0.01 & 0.01\\0.0 & 0.01 & 0.01\end{bmatrix}$ | $\begin{bmatrix}0.3 & 0.3 & 1.5\\0.3 & 0.3 & 1.5\\0.3 & 0.3 & 1.5\\0.3 & 0.3 & 1.5\end{bmatrix}$ | 1 | 1 | $\begin{bmatrix}5\\5\\5\\5\end{bmatrix}\times 10^4$ | $\begin{bmatrix}1\\1\\1\\1\end{bmatrix}\times 10^3$ | 0.76 |
| Fig. 5(b) | 6 | 8 | $\begin{bmatrix}\text{3DG}\\\text{3DG}\\\text{3DG}\\\text{3DG}\end{bmatrix}$ | $\begin{bmatrix}-0.3 & 0.0 & -0.5\\0.3 & 0.0 & -0.5\\0.0 & -0.3 & 0.5\\0.0 & 0.3 & 0.5\end{bmatrix}$ | $\begin{bmatrix}0.3 & 0.3 & 1.5\\0.3 & 0.3 & 1.5\\0.3 & 0.3 & 1.5\\0.3 & 0.3 & 1.5\end{bmatrix}$ | 1 | 1 | $\begin{bmatrix}5\\5\\5\\5\end{bmatrix}\times 10^4$ | $\begin{bmatrix}1\\1\\1\\1\end{bmatrix}\times 10^3$ | 0.75 |
| Fig. 5(c) | 6 | 8 | $\begin{bmatrix}\text{3DG}\\\text{3DG}\\\text{3DG}\\\text{3DG}\end{bmatrix}$ | $\begin{bmatrix}-1.0 & 0.0 & -1.5\\1.0 & 0.0 & -1.5\\0.0 & -1.0 & 1.5\\0.0 & 1.0 & 1.5\end{bmatrix}$ | $\begin{bmatrix}0.3 & 0.3 & 1.5\\0.3 & 0.3 & 1.5\\0.3 & 0.3 & 1.5\\0.3 & 0.3 & 1.5\end{bmatrix}$ | 1 | 1 | $\begin{bmatrix}5\\5\\5\\5\end{bmatrix}\times 10^4$ | $\begin{bmatrix}1\\1\\1\\1\end{bmatrix}\times 10^3$ | 1.83 |

TABLE S6. Parameter values used in the analyses of the traces. Choices are listed according to figures. Since, in this study we consider four confocal volumes $m = 1, 2, 3, 4$, we have four PSFs, molecular brightnesses and background photon emission rates for any of the respective figures. Additional values are listed in Table. S6

| | $\mathrm{PSF}_m$ | $[\mathrm{C}_{m,x}, \mathrm{C}_{m,y}, \mathrm{C}_{m,z}]$ | $[\omega_{m,x}, \omega_{m,y}, \omega_{m,z}]$ | N | $\alpha_D$ | $\beta_D$ | $\alpha_{\mathrm{mol}}$ | $\beta_{\mathrm{mol}}$ | $\alpha_{\mathrm{mol}}^{\mathrm{prop}}$ | $\alpha_{\mathrm{back}}$ | $\beta_{\mathrm{back}}$ | $\alpha_{\mathrm{back}}^{\mathrm{prop}}$ | $\gamma_b$ | $\begin{bmatrix} x_0 \\ y_0 \\ z_0 \end{bmatrix}$ | $\begin{bmatrix} \sigma_{x_0}^2 \\ \sigma_{y_0}^2 \\ \sigma_{z_0}^2 \end{bmatrix}$ |
|---|---|---|---|---|---|---|---|---|---|---|---|---|---|---|---|
| Units | - | $\mu$m | $\mu$m | - | - | $\mu\mathrm{m}^2\mathrm{s}^{-1}$ | - | pht $\mathrm{s}^{-1}$ | - | - | pht $\mathrm{s}^{-1}$ | - | - | $\mu$m | $\mu\mathrm{m}^2$ |
| Fig. 2(a) | $\begin{bmatrix} \text{3DG} \\ \text{3DG} \\ \text{3DG} \\ \text{3DG} \end{bmatrix}$ | $\begin{bmatrix} -0.3 & 0.0 & -0.5 \\ 0.3 & 0.0 & -0.5 \\ 0.0 & -0.3 & 0.5 \\ 0.0 & 0.3 & 0.5 \end{bmatrix}$ | $\begin{bmatrix} 0.3 & 0.3 & 1.5 \\ 0.3 & 0.3 & 1.5 \\ 0.3 & 0.3 & 1.5 \\ 0.3 & 0.3 & 1.5 \end{bmatrix}$ | 20 | 1 | 1 | 1 | $10^4$ | $10^3$ | 1 | $10^3$ | $10^2$ | 1 | $\begin{bmatrix} 0 \\ 0 \\ 0 \end{bmatrix}$ | $\begin{bmatrix} 4 \\ 4 \\ 4 \end{bmatrix}$ |
| Fig. 2(b) | $\begin{bmatrix} \text{3DG} \\ \text{3DG} \\ \text{3DG} \\ \text{3DG} \end{bmatrix}$ | $\begin{bmatrix} -0.3 & 0.0 & -0.5 \\ 0.3 & 0.0 & -0.5 \\ 0.0 & -0.3 & 0.5 \\ 0.0 & 0.3 & 0.5 \end{bmatrix}$ | $\begin{bmatrix} 0.3 & 0.3 & 1.5 \\ 0.3 & 0.3 & 1.5 \\ 0.3 & 0.3 & 1.5 \\ 0.3 & 0.3 & 1.5 \end{bmatrix}$ | 20 | 1 | 1 | 1 | $10^4$ | $10^3$ | 1 | $10^3$ | $10^2$ | 1 | $\begin{bmatrix} 0 \\ 0 \\ 0 \end{bmatrix}$ | $\begin{bmatrix} 4 \\ 4 \\ 4 \end{bmatrix}$ |
| Fig. 2(c) | $\begin{bmatrix} \text{3DG} \\ \text{3DG} \\ \text{3DG} \\ \text{3DG} \end{bmatrix}$ | $\begin{bmatrix} -0.3 & 0.0 & -0.5 \\ 0.3 & 0.0 & -0.5 \\ 0.0 & -0.3 & 0.5 \\ 0.0 & 0.3 & 0.5 \end{bmatrix}$ | $\begin{bmatrix} 0.3 & 0.3 & 1.5 \\ 0.3 & 0.3 & 1.5 \\ 0.3 & 0.3 & 1.5 \\ 0.3 & 0.3 & 1.5 \end{bmatrix}$ | 20 | 1 | 1 | 1 | $10^4$ | $10^3$ | 1 | $10^3$ | $10^2$ | 1 | $\begin{bmatrix} 0 \\ 0 \\ 0 \end{bmatrix}$ | $\begin{bmatrix} 4 \\ 4 \\ 4 \end{bmatrix}$ |
| Fig. 3(a) | $\begin{bmatrix} \text{3DG} \\ \text{3DG} \\ \text{3DG} \\ \text{3DG} \end{bmatrix}$ | $\begin{bmatrix} -0.3 & 0.0 & -0.5 \\ 0.3 & 0.0 & -0.5 \\ 0.0 & -0.3 & 0.5 \\ 0.0 & 0.3 & 0.5 \end{bmatrix}$ | $\begin{bmatrix} 0.3 & 0.3 & 1.5 \\ 0.3 & 0.3 & 1.5 \\ 0.3 & 0.3 & 1.5 \\ 0.3 & 0.3 & 1.5 \end{bmatrix}$ | 20 | 1 | 1 | 1 | $10^4$ | $10^3$ | 1 | $10^3$ | $10^2$ | 1 | $\begin{bmatrix} 0 \\ 0 \\ 0 \end{bmatrix}$ | $\begin{bmatrix} 4 \\ 4 \\ 4 \end{bmatrix}$ |
| Fig. 3(b) | $\begin{bmatrix} \text{3DG} \\ \text{3DG} \\ \text{3DG} \\ \text{3DG} \end{bmatrix}$ | $\begin{bmatrix} -0.3 & 0.0 & -0.5 \\ 0.3 & 0.0 & -0.5 \\ 0.0 & -0.3 & 0.5 \\ 0.0 & 0.3 & 0.5 \end{bmatrix}$ | $\begin{bmatrix} 0.3 & 0.3 & 1.5 \\ 0.3 & 0.3 & 1.5 \\ 0.3 & 0.3 & 1.5 \\ 0.3 & 0.3 & 1.5 \end{bmatrix}$ | 20 | 1 | 1 | 1 | $10^4$ | $10^3$ | 1 | $10^3$ | $10^2$ | 1 | $\begin{bmatrix} 0 \\ 0 \\ 0 \end{bmatrix}$ | $\begin{bmatrix} 4 \\ 4 \\ 4 \end{bmatrix}$ |
| Fig. 3(c) | $\begin{bmatrix} \text{3DG} \\ \text{3DG} \\ \text{3DG} \\ \text{3DG} \end{bmatrix}$ | $\begin{bmatrix} -0.3 & 0.0 & -0.5 \\ 0.3 & 0.0 & -0.5 \\ 0.0 & -0.3 & 0.5 \\ 0.0 & 0.3 & 0.5 \end{bmatrix}$ | $\begin{bmatrix} 0.3 & 0.3 & 1.5 \\ 0.3 & 0.3 & 1.5 \\ 0.3 & 0.3 & 1.5 \\ 0.3 & 0.3 & 1.5 \end{bmatrix}$ | 20 | 1 | 1 | 1 | $10^4$ | $10^3$ | 1 | $10^3$ | $10^2$ | 1 | $\begin{bmatrix} 0 \\ 0 \\ 0 \end{bmatrix}$ | $\begin{bmatrix} 4 \\ 4 \\ 4 \end{bmatrix}$ |
| Fig. 4(a) | $\begin{bmatrix} \text{3DG} \\ \text{3DG} \\ \text{3DG} \\ \text{3DG} \end{bmatrix}$ | $\begin{bmatrix} -0.3 & 0.0 & -0.5 \\ 0.3 & 0.0 & -0.5 \\ 0.0 & -0.3 & 0.5 \\ 0.0 & 0.3 & 0.5 \end{bmatrix}$ | $\begin{bmatrix} 0.3 & 0.3 & 1.5 \\ 0.3 & 0.3 & 1.5 \\ 0.3 & 0.3 & 1.5 \\ 0.3 & 0.3 & 1.5 \end{bmatrix}$ | 20 | 1 | 1 | 1 | $10^4$ | $10^3$ | 1 | $10^3$ | $10^2$ | 1 | $\begin{bmatrix} 0 \\ 0 \\ 0 \end{bmatrix}$ | $\begin{bmatrix} 4 \\ 4 \\ 4 \end{bmatrix}$ |
| Fig. 4(b) | $\begin{bmatrix} \text{3DG} \\ \text{3DG} \\ \text{3DG} \\ \text{3DG} \end{bmatrix}$ | $\begin{bmatrix} -0.3 & 0.0 & -0.5 \\ 0.3 & 0.0 & -0.5 \\ 0.0 & -0.3 & 0.5 \\ 0.0 & 0.3 & 0.5 \end{bmatrix}$ | $\begin{bmatrix} 0.3 & 0.3 & 1.5 \\ 0.3 & 0.3 & 1.5 \\ 0.3 & 0.3 & 1.5 \\ 0.3 & 0.3 & 1.5 \end{bmatrix}$ | 20 | 1 | 1 | 1 | $10^4$ | $10^3$ | 1 | $10^3$ | $10^2$ | 1 | $\begin{bmatrix} 0 \\ 0 \\ 0 \end{bmatrix}$ | $\begin{bmatrix} 4 \\ 4 \\ 4 \end{bmatrix}$ |
| Fig. 4(c) | $\begin{bmatrix} \text{3DG} \\ \text{3DG} \\ \text{3DG} \\ \text{3DG} \end{bmatrix}$ | $\begin{bmatrix} -0.3 & 0.0 & -0.5 \\ 0.3 & 0.0 & -0.5 \\ 0.0 & -0.3 & 0.5 \\ 0.0 & 0.3 & 0.5 \end{bmatrix}$ | $\begin{bmatrix} 0.3 & 0.3 & 1.5 \\ 0.3 & 0.3 & 1.5 \\ 0.3 & 0.3 & 1.5 \\ 0.3 & 0.3 & 1.5 \end{bmatrix}$ | 20 | 1 | 1 | 1 | $10^4$ | $10^3$ | 1 | $10^3$ | $10^2$ | 1 | $\begin{bmatrix} 0 \\ 0 \\ 0 \end{bmatrix}$ | $\begin{bmatrix} 4 \\ 4 \\ 4 \end{bmatrix}$ |
| Fig. 5(a) | $\begin{bmatrix} \text{3DG} \\ \text{3DG} \\ \text{3DG} \\ \text{3DG} \end{bmatrix}$ | $\begin{bmatrix} -0.01 & 0.0 & -0.01 \\ 0.01 & 0.0 & -0.01 \\ 0.0 & -0.01 & 0.01 \\ 0.0 & 0.01 & 0.01 \end{bmatrix}$ | $\begin{bmatrix} 0.3 & 0.3 & 1.5 \\ 0.3 & 0.3 & 1.5 \\ 0.3 & 0.3 & 1.5 \\ 0.3 & 0.3 & 1.5 \end{bmatrix}$ | 20 | 1 | 1 | 1 | $10^4$ | $10^3$ | 1 | $10^3$ | $10^2$ | 1 | $\begin{bmatrix} 0 \\ 0 \\ 0 \end{bmatrix}$ | $\begin{bmatrix} 4 \\ 4 \\ 4 \end{bmatrix}$ |
| Fig. 5(b) | $\begin{bmatrix} \text{3DG} \\ \text{3DG} \\ \text{3DG} \\ \text{3DG} \end{bmatrix}$ | $\begin{bmatrix} -0.3 & 0.0 & -0.5 \\ 0.3 & 0.0 & -0.5 \\ 0.0 & -0.3 & 0.5 \\ 0.0 & 0.3 & 0.5 \end{bmatrix}$ | $\begin{bmatrix} 0.3 & 0.3 & 1.5 \\ 0.3 & 0.3 & 1.5 \\ 0.3 & 0.3 & 1.5 \\ 0.3 & 0.3 & 1.5 \end{bmatrix}$ | 20 | 1 | 1 | 1 | $10^4$ | $10^3$ | 1 | $10^3$ | $10^2$ | 1 | $\begin{bmatrix} 0 \\ 0 \\ 0 \end{bmatrix}$ | $\begin{bmatrix} 4 \\ 4 \\ 4 \end{bmatrix}$ |
| Fig. 5(c) | $\begin{bmatrix} \text{3DG} \\ \text{3DG} \\ \text{3DG} \\ \text{3DG} \end{bmatrix}$ | $\begin{bmatrix} -1.0 & 0.0 & -1.5 \\ 1.0 & 0.0 & -1.5 \\ 0.0 & -1.0 & 1.5 \\ 0.0 & 1.0 & 1.5 \end{bmatrix}$ | $\begin{bmatrix} 0.3 & 0.3 & 1.5 \\ 0.3 & 0.3 & 1.5 \\ 0.3 & 0.3 & 1.5 \\ 0.3 & 0.3 & 1.5 \end{bmatrix}$ | 20 | 1 | 1 | 1 | $10^4$ | $10^3$ | 1 | $10^3$ | $10^2$ | 1 | $\begin{bmatrix} 0 \\ 0 \\ 0 \end{bmatrix}$ | $\begin{bmatrix} 4 \\ 4 \\ 4 \end{bmatrix}$ |
